# Conserved Vulnerability, Local Tunability: How Binding Proteins Read Ligand Fluorination

**DOI:** 10.64898/2026.11.17.767237

**Authors:** Paul Matthay, Amy Gooch, Yusran Abdillah Muthahari, Ben E. Clifton, Ai Shinobu, Paola Laurino

**Author notes:** These authors contributed equally (Paul Matthay, Amy Gooch).

## Abstract

While rare in nature, fluorinated compounds are ubiquitous in synthetic chemicals ranging from pharmaceuticals to agrochemicals, yet how proteins recognize these anthropogenic molecules remains poorly understood. Here, we use three structurally and evolutionarily divergent solute-binding proteins (SBPs) specific for short-chain dicarboxylic acids as a comparative model system to dissect the principles governing protein-fluorocompound recognition. X-ray crystal structures reveal that each SBP uses a distinct binding mode, ranging from backbone-mediated to side-chain-dominated hydrogen bonding. Despite their structural divergence, ITC shows that all three proteins exhibit a conserved vulnerability upon binding perfluorinated ligands: binding affinity drops up to 80-fold, or is abolished, across all three proteins regardless of fold or binding mode. In contrast, sensitivity to partial fluorination is locally tunable, with one protein even binding a fluorinated ligand more tightly than its non-fluorinated counterpart. Using X-ray crystallography, molecular dynamics simulations, and site-directed mutagenesis, we show that aromatic residues positioned near the ligand contribute to this tunability, capable of either penalizing or favoring fluorinated ligands. Together, these findings establish that ligand-fluorination sensitivity reflects a conserved chemical vulnerability layered with locally tunable, structurally encoded selectivity, providing a roadmap for predicting and engineering protein recognition of fluorinated compounds.

## Introduction

Fluorine is the most abundant halogen on earth, yet almost nonexistent in natural products, with a few exceptions such as fluoroacetate (1). Despite its rarity in nature, fluorine is widely used in synthetic chemistry due to its high electronegativity (2), low polarizability (3), and small size (atomic radius 1.47 Å) (4) making it a powerful tool for tuning molecular properties. Exchanging a C-H bond for C-F increases the metabolic stability of drugs, modulates lipophilicity and cell permeability, and perturbs the p*K*_a_ of nearby functional groups, thereby altering binding affinity and bioavailability (5, 6). These effects have made fluorine a pillar of drug design: indeed, more than 20% of pharmaceuticals now contain at least one fluorine atom (7, 8). Fluorine also finds extensive use in the family of chemicals known as PFAS (per- and polyfluoroalkyl substances), which are gaining attention due to their harmful effects on human and environmental health (9). Yet despite its widespread use, how fluorination reshapes protein-ligand interactions remains difficult to predict (3), raising the question of whether such effects follow shared principles or depend on the specific architecture of the binding site.

Previous studies have illuminated some of the important binding interactions between proteins and fluorinated compounds (10-16). Due to the small size of fluorine, selectively fluorinated ligands typically adopt a similar binding pose within proteins compared to their corresponding unmodified ligands (10), while the fluorine atom(s) may oriented towards fluorophilic environments in the protein, such as backbone peptide groups (11). Although the high electronegativity of fluorine might suggest strong hydrogen bonding potential, its low polarizability makes it a poor hydrogen-bond donor (17). Instead, fluorine frequently engages in C–F···C=O contacts with backbone or side-chain carbonyls (11-13). The geometry of this interaction is well-defined and can be used to predict fluorophilic sites in protein structures (13). In contrast, the binding mechanisms of per- and polyfluorinated compounds to proteins are less well understood. Several studies have shown that binding of perfluorinated compounds is driven by hydrophobic effects (14, 15). However, the effect of ligand perfluorination has only been studied in single or closely related proteins. Therefore, it is unknown how perfluorination affects ligand binding affinity across structurally divergent binding sites.

Here, we systematically characterize the effect of stepwise ligand fluorination, up to perfluorination, on protein-ligand interactions using three structurally and evolutionarily divergent proteins that belong to the solute-binding protein (SBP) superfamily. We selected SBPs specific for short-chain dicarboxylic acids as a model system because carboxylic acid groups are a feature of many industrially relevant PFAS (18). Despite sharing low sequence identity and distinct binding motifs, we found that for all three SBPs, the native protein-ligand interactions are similarly vulnerable to complete fluorination of the ligand’s central carbons, indicating a fold-independent chemical constraint on recognition. In contrast, tolerance to partial fluorination diverges between proteins, in one case even favoring the fluorinated ligand over its native counterpart. Combining X-ray crystallography, molecular dynamics simulations, and site-directed mutagenesis, we identify aromatic residues near the ligand binding site as a key contributor to this divergence, offering a structural basis for predicting when fluorination will be tolerated, penalized or preferred.

## Results

### Selection of solute-binding proteins to study the effect of ligand fluorination on ligand recognition

To systematically characterize the effect of ligand fluorination on protein-ligand interactions, we combined isothermal titration calorimetry (ITC) and X-ray crystallography with molecular dynamics (MD) simulations. Three representative SBPs that show binding to dicarboxylates were selected: B5M07_04440 from *Sulfitobacter* sp. (unpublished results), SAR11_1361 from *Pelagibacter ubiqueversans* (19), and Sdel_0447 from *Sulfurospirillum deleyianum* (20). B5M07_04440 and SAR11_1361 are both the soluble extracytoplasmic subunit of an ATP binding cassette (ABC) transporter, whereas Sdel_0447 is the soluble extracytoplasmic subunit of a tripartite ATP-independent periplasmic (TRAP) transporter. Notably, these three SBPs belong to distinct subfamilies of the SBP superfamily (the distinct PFAM codes of which are listed in Table S8) and share low sequence identity of 16 to 27% (Figure S1). These features allow assessment of fluorination effects across structurally and evolutionarily distinct binding sites.

X-ray crystallographic analysis was first performed for each SBP with their native ligand, succinate. We determined the X-ray crystal structures of B5M07_04440 and SAR11_1361 at resolutions of 1.36 Å and 1.52 Å, respectively, while the 1.8 Å structure of Sdel_0447 has been previously reported (20). B5M07_04440 and Sdel_0447 both have the periplasmic binding protein-like II fold, whereas SAR11_1361 has the periplasmic binding protein-like I fold (Figure 1a-c). Further analysis revealed distinct ligand-binding modes for each protein. B5M07_04440 stabilizes succinate predominantly through hydrogen bonds involving the protein backbone, whereas SAR11_1361 and Sdel_0447 rely primarily on side-chain-mediated interactions (Figure 1d-f). In B5M07_04440, one carboxylate group of succinate forms hydrogen bonds with the backbone amide and sidechain hydroxyl of Thr151, while the second carboxylate is stabilized by the backbone amides of Thr41 and Tyr42, and the sidechain amine of Trp174 (Figure 1e). In SAR11_1361, one carboxylate interacts with the backbone amide and sidechain hydroxyl of Ser114, as well as with the imidazole of His239 and backbone of Ala137. The other carboxylate is stabilized by hydrogen bonds with the side chains of Tyr112, Tyr159, His264, Tyr318, and Gln319 (Figure 1d). In contrast, Sdel_0447 binds succinate exclusively via side chain interactions: one carboxylate is stabilized by Lys38, Asn211, and Tyr234, and the other by Lys91 and two hydrogen bonds with Arg167 (Figure 1f). Thus, the three SBPs show structurally distinct binding sites and different binding modes for their native ligand.

**Figure 1.**
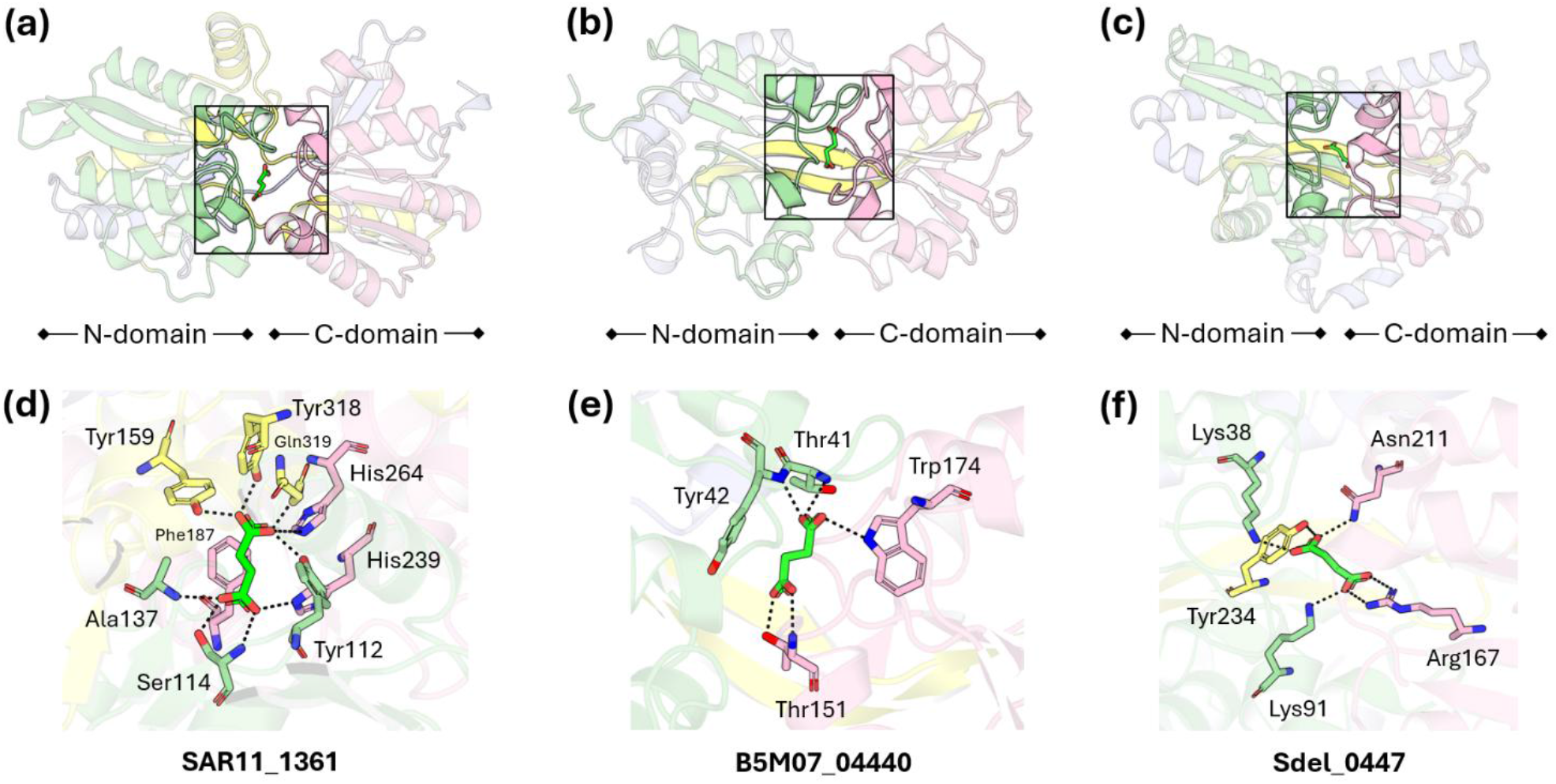
Structures of solute-binding proteins specific for dicarboxylates. X-ray crystal structure of succinate bound to (a) SAR11_1361, (b) B5M07_04440, and (c) Sdel_0447 (PDB code: 4OVS). Binding mode of succinate within the crystal structure of (d) SAR11_1361, (e) B5M07_04440, and (f) Sdel_0447 (PDB code: 4OVS).

### Partial ligand fluorination reveals protein-specific tunability

To evaluate the effect of ligand fluorination on binding affinity for a variety of dicarboxylates, we performed ITC using the saturated C_4_ dicarboxylate succinate, the unsaturated C_4_ derivative fumarate, and the analogous C_5_ dicarboxylate glutarate, together with their stepwise fluorinated derivatives (Figure 2a). We expected the SBPs to bind tightly to the dicarboxylates present in the expression host during protein expression (19), therefore, the proteins were denatured and refolded prior to ITC measurements in order to obtain the apo proteins and allow accurate determination of binding parameters. B5M07_04440 and SAR11_1361 displayed higher binding affinity than Sdel_0447 across the range of dicarboxylates tested. For instance, the *K*_d_ values of succinate binding to B5M07_04440, SAR11_1361, and Sdel_0447 were <10 nM, 31.1 nM, and 329 nM, respectively (Figure 2b). A complete overview of the ITC measurements and derived binding parameters are provided in the supporting information (Table S4 and Figures S2-7).

**Figure 2.**
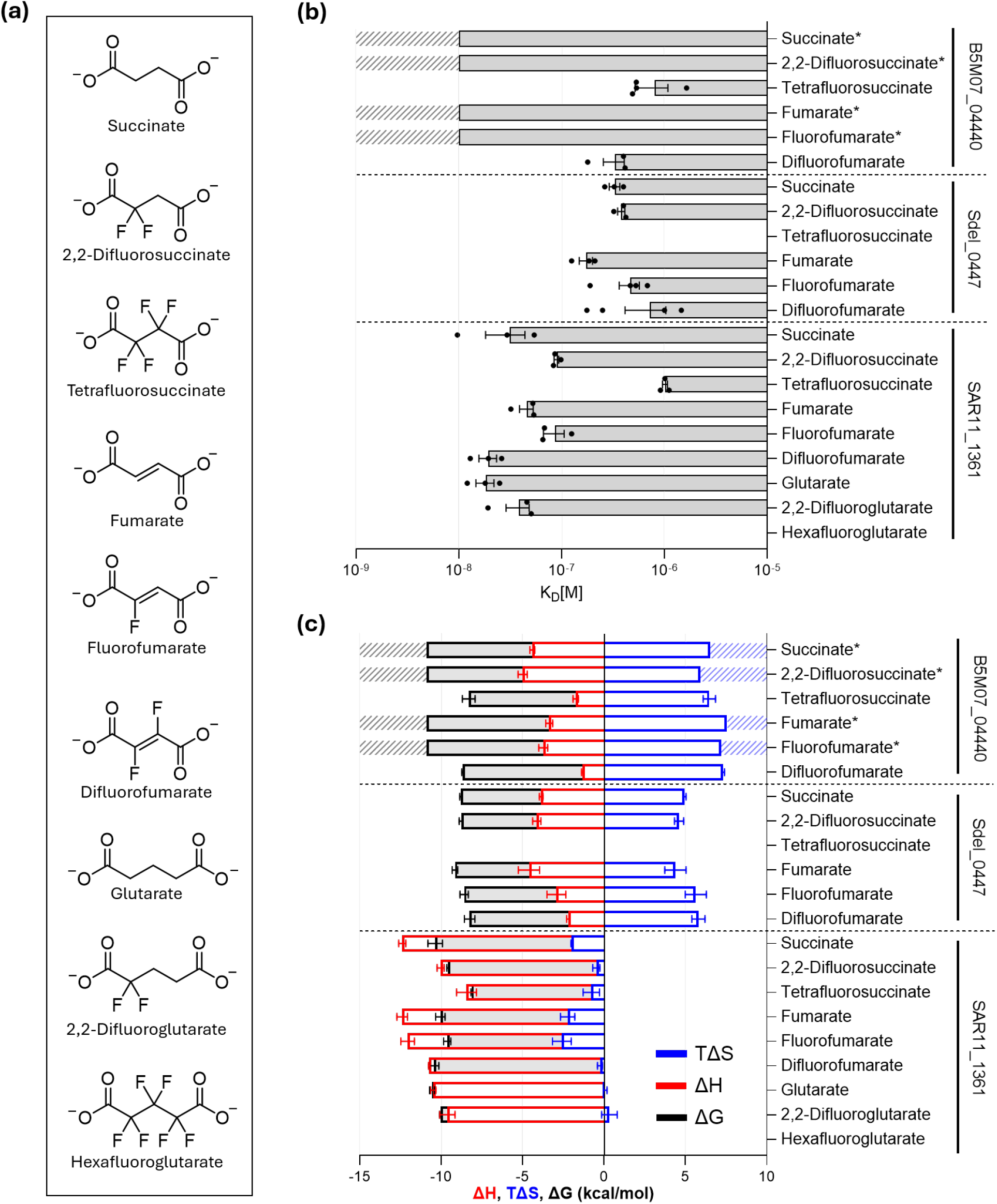
Binding affinity of solute binding proteins with dicarboxylate ligands and their fluorinated derivatives. (a) Structures of the ligands used in ITC experiments. (b) *K*_d_ values and (c) thermodynamic parameters measured using ITC. Protein samples were unfolded, refolded, purified and concentrated to 32 µM with freshly prepared ligands at a concentration of 280 µM. Values were taken in technical replicates, which were combined with inverse-variance weighting, and n≥3 biological replicates. Asterisks mark the ligands for which the binding affinity was above the limit of quantitation of ITC, and for which the *K*_d_ is reported as <10 nM.

Stepwise fluorination of succinate revealed the following effects across the three SBPs. Difluorination of one of the central carbons (i.e. 2,2-difluorosuccinate) did not significantly affect the binding affinity for Sdel_0447; however, for SAR11_1361 there was a 2.9-fold decrease in binding affinity, and for B5M07_04440, the binding affinity remained high (<10 nM). Difluorination of both central carbons to give the perfluorinated derivative tetrafluorosuccinate strongly impaired binding across the three SBPs, resulting in a greater than 80-fold decrease in binding affinity for B5M07_04440, a 33-fold decrease in binding affinity for SAR11_1361, and abolishing detectable binding for Sdel_0447 (Figure 2b).

For each SBP, the binding affinity for fumarate was similar to that for succinate, yet stepwise fluorination produced more variable effects. Fluorination of one of the central carbons (i.e. fluorofumarate) did not significantly affect binding affinity for SAR11_1361 or Sdel_0447, and for B5M07_04440, the binding affinity remained high (<10 nM). However, fluorination of both central carbons (i.e. difluorofumarate) resulted in a greater than 33-fold decrease in binding affinity for B5M07_04440 and no significant change in binding affinity for Sdel_0447. Interestingly, for SAR11_1361 there was a slight, yet significant, 4.4-fold increase in binding affinity for difluorofumarate compared with fluorofumarate (Figure 2b).

Glutarate binding was only observed in SAR11_1361. Difluorination at one central carbon to give 2,2-difluoroglutarate, resulted in a 2.1-fold decrease in binding affinity, whilst perfluorination to hexafluoroglutarate abolished detectable binding (Figure 2b).

The thermodynamic driving force of binding for each SBP-ligand combination can be characterized by the following equation:

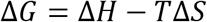

Where *ΔG* represents the change in Gibbs free energy, *ΔH* the change in enthalpy, *T* the temperature, and *ΔS* the change in entropy. ITC measures the heat released or absorbed upon ligand binding. Fitting the resulting binding isotherms provides estimates of *ΔH* and the association constant *K*_a_. *ΔG* can be calculated from *K*_a,_ and the -*TΔS* term can then be solved by *ΔG* - *ΔH*. Our data reveal that ligand binding to SAR11_1361 is enthalpically driven: the favorable *ΔH* contributes most to *ΔG*, while the positive -*TΔS* serves as a penalty. In contrast, for B5M07_04440 and Sdel_0447, ligand binding is more entropically driven (Figure 2c), with the negative -*TΔS* contributing favorably to *ΔG*. This is also mirrored in the crystal structures, where SAR11_1361 can form up to nine hydrogen bonds to succinate, whereas B5M07_04440 and Sdel_0447 can only form a maximum of five and six bonds each, respectively (Figure 1d-f).

Together, these results reveal a conserved vulnerability to complete fluorination of the ligand’s central carbons, consistently reducing or abolishing binding across all three structurally divergent SBPs. In contrast, sensitivity to partial fluorination is locally tunable: B5M07_04440 and SAR11_1361 differ sharply in their response to fluorination of fumarate, with SAR11_1361 showing increased affinity for difluorofumarate relative to fluorofumarate — the only case of fluorination-enhanced binding we observed. This divergence suggests that tolerance to partial fluorination is governed by local structural features rather than the protein’s global fold. To further investigate this behavior, we focused on SAR11_1361 and B5M07_04440 as they show both high affinity and fold changes in binding affinities across ligands.

### Biophysical characterization of the binding mode of fluorinated ligands

To investigate how ligand fluorination influences binding at the structural level, we compared the binding modes of succinate, 2,2-difluorosuccinate, and tetrafluorosuccinate to B5M07_04440 using X-ray crystallography and MD simulations. In addition to the succinate-bound structure, we successfully solved the crystal structure of B5M07_04440 in complex with 2,2-difluorosuccinate (Figure 3a). The structure shows that 2,2-difluorosuccinate adopts essentially the same overall binding mode as succinate, indicating that the fluorinated ligand is accommodated within the same binding pocket without any major conformational shifts. Indeed, the binding mode appears highly similar except for a slight increase in the distance between the ligand and Trp174 from approximately 3.1 Å to 3.5 Å (Figure 3b,c), explaining why binding affinity is largely maintained upon fluorination.

**Figure 3.**
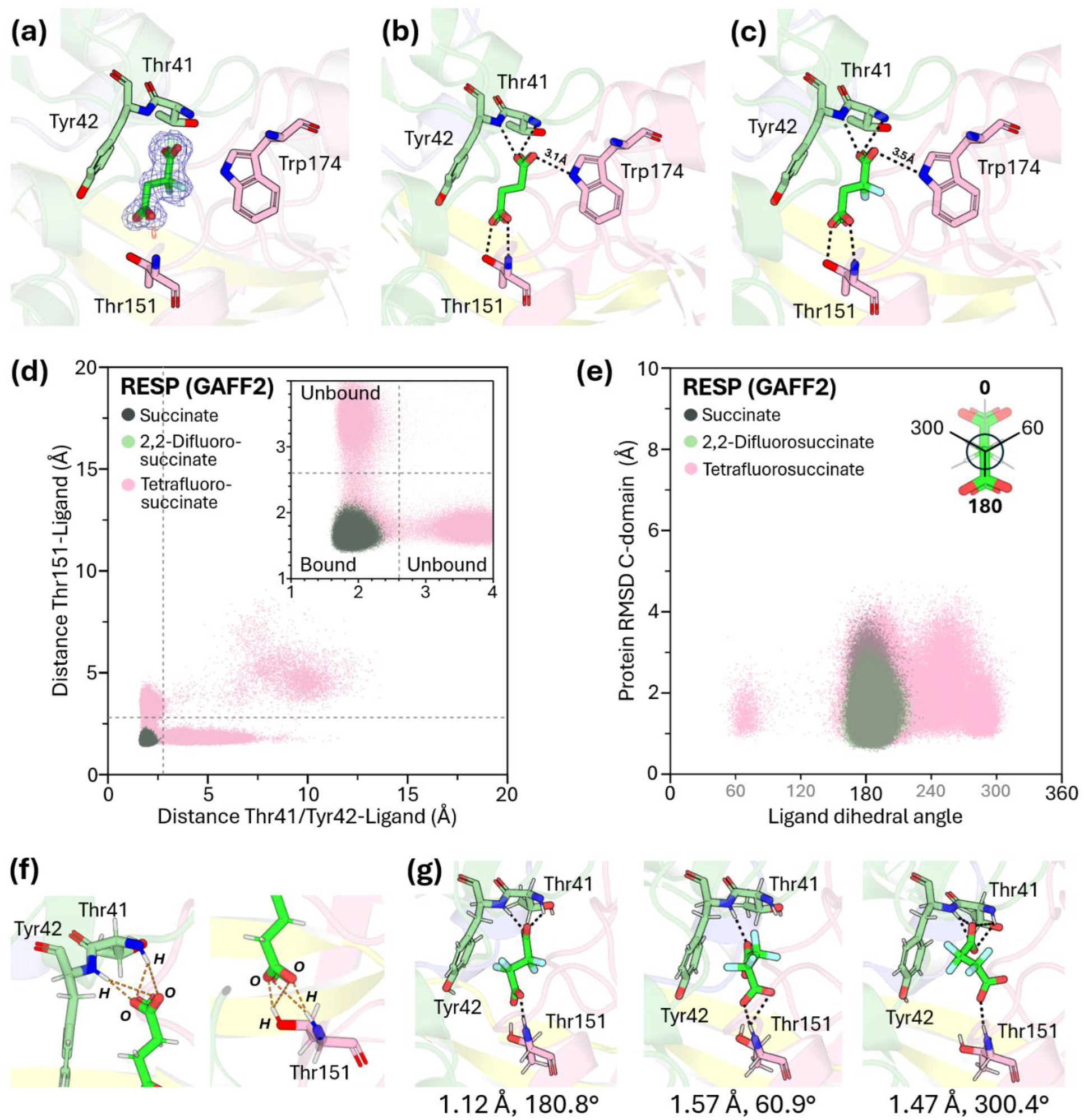
Tetrafluorination but not difluorination of succinate results in unproductive binding poses. (a) The electron density of 2,2-difluorosuccinate bound in the X-ray crystal structure of B5M07_04440 reveals the preferred binding orientation, whereby the fluorine atoms face toward Trp174. The 2F_o_–F_c_ electron density map is represented by blue mesh (σ = 1.2), and the Fo-Fc weighted difference map by green and red mesh (σ = ±2.5). (b-c) Binding modes of (b) succinate and (c) 2,2-difluorosuccinate within the crystal structure of B5M07_04440. (d) Distribution of conformations sampled during MD simulations of the B5M07_04440 complexes with succinate, 2,2-difluorosuccinate, and tetrafluorosuccinate in the two-dimensional space defined by two protein–ligand distances. The x-axis represents the minimum distance between the ligand carboxylate oxygen atoms and the backbone amide hydrogen atoms of Thr41/Tyr42, while the y-axis represents the corresponding minimum distance to the backbone amide hydrogen atom of Thr151. The grey dashed lines indicate a 2.6 Å distance cutoff used to define the bound state. Three independent replicates were performed for each condition, with 1 μs per replicate. (e) Distribution of bound-state conformations in the two-dimensional space defined by the C-domain RMSD and ligand dihedral angle. The C-domain RMSD was calculated after superimposing the backbone atoms (C, CA, N, and O) of the N-domain onto the initial closed-state structure (Figure S13). (f) Definition of the protein–ligand distances used to characterize the bound state. Orange dashed lines indicate the atom pairs considered for each distance calculation, with the minimum distance used for each simulation frame. (g) Representative snapshots of tetrafluorosuccinate conformations with different dihedral angles sampled during simulation of the protein– tetrafluorosuccinate complex. Polar contacts between the protein and ligand are shown as black dashed lines. The tetrafluorosuccinate dihedral angle and corresponding C-domain RMSD are indicated below each snapshot. Polar contacts were identified using PyMOL (21) with the default settings (3.2 Å cutoff). Corresponding analyses using AM1-BCC charges and the GAFF force field for the ligands are shown in Figure S9.

Analysis of the electron density also revealed a preferred orientation of the two fluorine atoms within the binding pocket, facing towards Trp174 rather than Tyr42 (Figure 3a). The observed structure reveals that the binding site can accommodate the difluorinated ligand without introducing significant steric clashes or unfavorable electrostatic interactions. Considering the binding affinity for 2,2-difluorosuccinate is largely unaffected, whereas the binding affinity for tetrafluorosuccinate is strongly disrupted, we hypothesize that additional fluorination may lead to unfavorable SBP-ligand interactions.

To further understand the structural basis for the observed differences in binding affinity, we performed MD simulations for the succinate-, 2,2-difluorosuccinate-, and tetrafluorosuccinate-bound complexes of B5M07_04440. The experimental tetrafluorosuccinate-bound structure could not be obtained, so instead we generated the complex by replacing the two methylene hydrogen atoms of the 2,2-difluorosuccinate-bound structure with fluorine atoms (Figure S12). MD simulations showed that the tetrafluorosuccinate complex sampled more time in the unbound states than the succinate or 2,2-difluorosuccinate complexes (Figure 3d,f and S10). In the unbound states, tetrafluorosuccinate cannot maintain stable interactions between its two carboxylates and the protein, as indicated by their long distances over the course of the simulation (Figure S9 and S10). In these tetrafluorosuccinate-unbound states, the protein sampled a more open conformation, as shown by a higher RMSD of the C-domain (Figure S13). Moreover, in the bound state, succinate and 2,2-difluorosuccinate populated mostly anti-conformations (180°) of the dihedral angle, forming stable closed-state conformations, whereas tetrafluorosuccinate sampled more non-anti conformations (Figure 3e,g and S8). In summary, our simulations explain the ITC-derived affinities (Figure 2b) and further reveal how the loss of binding manifests conformationally; tetrafluorosuccinate’s inability to maintain stable carboxylate contacts drive the protein toward more open, unbound-like states, whereas succinate and 2,2-difluorosuccinate remain confined to the stable closed-state, anti-conformation ensemble.

### Adjacent aromatic rings play a vital role in ligand recognition

The aliphatic chain of succinate forms a hydrophobic interaction with an aromatic ring in both B5M07_04440 and SAR11_1361. Thus, we hypothesized that these aromatic rings might be important for discrimination of the native ligand from its fluorinated derivatives. To address this question, we tested the effect of a Tyr42Ala substitution in B5M07_04440 (Figure 4a) and a Phe187Ala substitution in SAR11_1361 (Figure 4d) on binding fluorinated vs unfluorinated ligands. In both mutants, there was a significant decrease in binding affinity for the unfluorinated ligands compared with the wild-type proteins. This is consistent with the importance of hydrophobic interactions with the aliphatic chain for binding affinity. In the B5M07_04440 Tyr42Ala mutant, the decrease in binding affinity was generally a result of a reduction in the favorable entropic contribution to binding, with minimal effect on enthalpy, compared to wild type (Figure 4c). In contrast, for the SAR11_1361 Phe187Ala mutant, the decrease in binding affinity was largely a consequence of a reduction in favorable binding enthalpy rather than entropy (Figure 4f).

**Figure 4.**
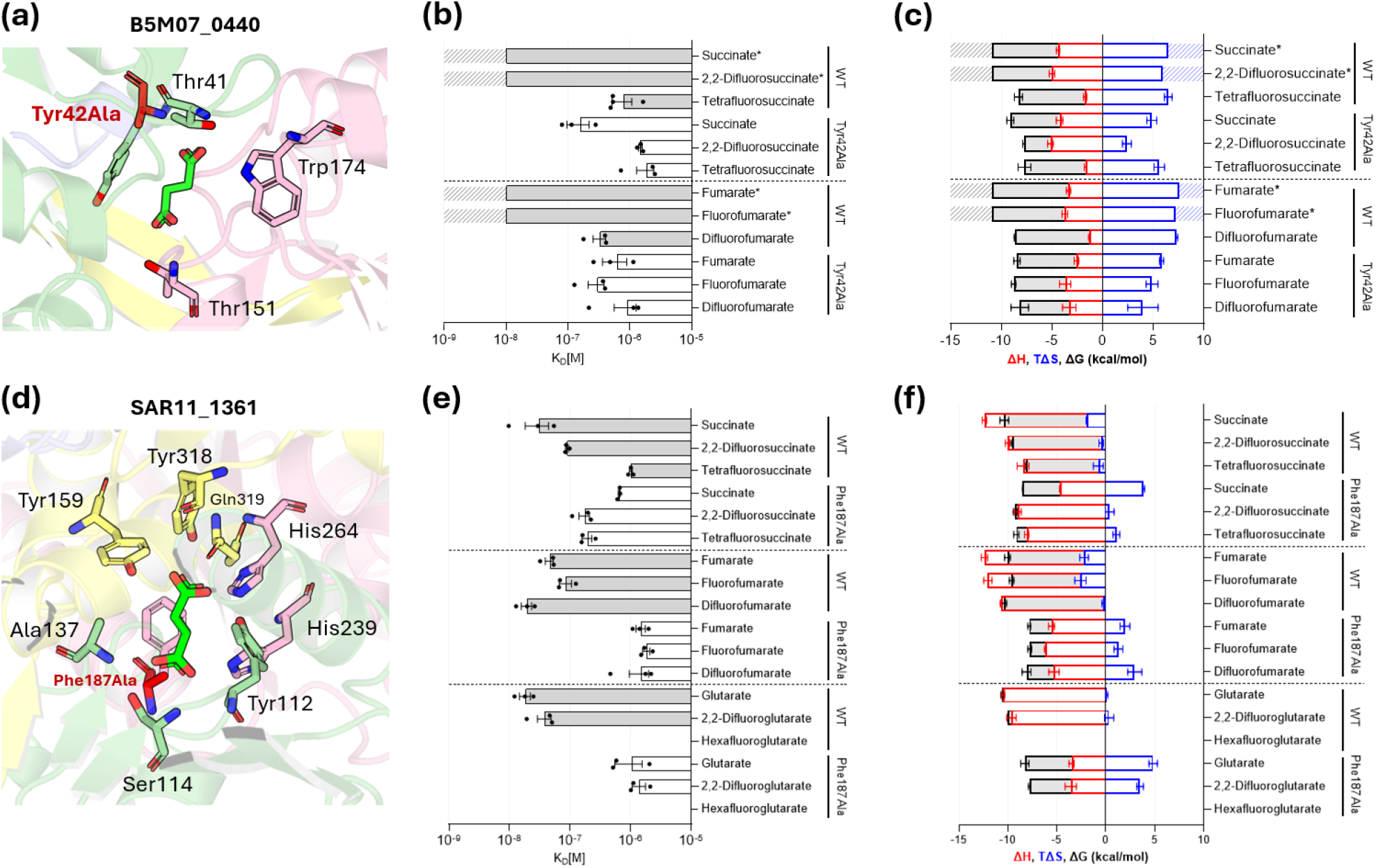
Mutagenesis validates the importance of an aromatic ring for ligand binding in B5M07_04440 and SAR11_1361. B5M07_04440 Tyr42Ala (a) ligand binding site, (b) *K*_d_ values, and (c) thermodynamic parameters. SAR11_1361 Phe187Ala (d) ligand binding site, (e) *K*_d_ values, and (f) thermodynamic parameters. *K*_d_ values were measured using ITC. Protein samples were unfolded, refolded, purified, concentrated to 32 µM, and titrated with freshly prepared ligands at a concentration of 280 µM. Values were taken in technical replicates, which were combined with inverse-variance weighting, and n≥3 biological replicates. Asterisks mark the ligands for which the binding affinity was above the limit of quantitation of ITC, and for which the *K*_d_ is reported as <10 nM. For (b), (c), (e), and (f), data from the wild-type proteins is reproduced from Figure 2 for comparison with the mutants.

The B5M07_04440 Tyr42Ala mutant showed similar binding affinity for fumarate, fluorofumarate, and difluorofumarate (*K*_d_ values 628 nM, 298 nM, and 912 nM, respectively). This indicates a minimal effect of fumarate fluorination on binding in the mutant B5M07_04440, unlike what we observe for the wild type. Yet, with succinate, the B5M07_04440 mutant maintains its vulnerability to the fluorinated derivatives, showing a significant decrease in binding affinity for both 2,2-difluorosuccinate and tetrafluorosuccinate, compared to succinate (Figure 4b). In the case of SAR11_1361, the Phe187Ala substitution resulted in a significant decrease in binding affinity for almost all ligands, except for 2,2-difluorosuccinate which had no significant change in binding affinity, and tetrafluorosuccinate which had a 5.2-fold increase in binding affinity compared to wild type (Figure 4e). This is particularly surprising considering that the SAR11_1361 Phe187Ala mutant displays a 21-fold decrease in binding affinity for succinate compared to wild type SAR11_1361.

## Discussion

In this work, we compared three structurally and evolutionary divergent SBPs to ask whether the effects of ligand fluorination on protein-ligand recognition follow conserved chemical principles or depend on protein-specific architecture. We conclude that ligand fluorination has both conserved and structure-specific effects, but at different levels. Complete ligand fluorination is a conserved vulnerability; it diminishes or abolishes binding regardless of the fold or binding mode. In contrast, partial fluorination revealed the protein’s local tunability, as it affects binding in a more varied way: fluorination at the same position can have a neutral effect on binding affinity for one protein, penalize affinity in another, or enhance affinity for a third. Here, we investigated which factors contribute to fluorine sensitivity in SBPs, as well as what features allow for a certain level of tunability to accommodate binding of fluorinated ligands.

Our results show that SBPs achieve high binding affinity through different thermodynamic strategies and binding-pocket configurations. For SAR11_1361, ligand binding is enthalpically driven but with an entropic penalty (Figure 2c); it has a tight binding pocket that, when succinate is bound, does not contain any ordered water molecules (Figure S15). Here, succinate adopts a twisted conformation (Figure S16). These observations point to the ligand being drawn into a highly bonded yet constrained pose, enforced by hydrophobic packing of the nearby aromatic ring of Phe187. B5M07_04440 and Sdel_0447 instead rely on a favorable entropy of binding; they maintain a looser binding pocket with more water molecules bound and where succinate adopts the canonical anti-conformation (Figure S15-16). These differences in binding pocket constraints may help explain why the same level of ligand fluorination can have opposite consequences in distinct proteins. A tight binding pocket that holds the ligand in a rigid, highly bonded conformation has more to lose from the further conformational restriction that ligand fluorination imposes, whereas a binding pocket with a looser fit may be more easily disrupted instead when fluorination changes ligand geometry or electrostatics.

We identified aromatic residues in the binding pocket (Tyr42 in B5M07_04440 and Phe187 in SAR11_1361) that are highly conserved across protein homologues (Figure S14) indicating a significant role in binding. Their contribution is hydrophobic and likely also π-mediated (CH-π interactions with succinate or π-π stacking interactions with fumarate). These contacts help position the ligand for hydrogen bonding by the surrounding side chains and backbone groups (Figure 4a,d), rather than directly forming hydrogen bonds to the ligand themselves. This case is relevant to B5M07_04440, as the ring provides significant entropic contributions upon binding and supports ligand positioning. The entropic contribution is likely due to the displacement of water molecules surrounding both the ligand’s aliphatic chain and the aromatic ring into the bulk solvent during ligand binding. In SAR11_1361, the ring instead imposes an entropic penalty that disproportionally disfavors binding to fluorinated ligands. Indeed, substituting this aromatic residue in SAR11_1361 relieves the penalty, but at an enthalpic cost, which results in an overall reduction in affinity for succinate. Remarkably, this mutation also shifts SAR11_1361’s binding specificity toward the fluorinated succinate derivatives. Removing the ring likely relieves the constraint on succinate binding in a highly bonded pose but allows binding of fluorinated ligands that may sample different conformational ensembles. In summary, it appears that the aromatic ring is employed in each protein to either buffer or amplify the consequences of fluorination on binding. The functional consequence of the aromatic ring for fluorination sensitivity depends on the local geometry context, but more broadly may act as a tunable feature for ligand fluorination sensitivity.

Our MD simulations show that the ligand conformational ensemble may contribute to differences in SBP binding affinity for various fluorinated ligands, similar to observations in another independent case (22). It is well known that substitution of C-H with C-F can alter ligand conformation (23, 24), and for succinate, may shift its conformational ensemble relative to that of the non-fluorinated ligand (25). Consistent with this hypothesis, we observed that the fully fluorinated tetrafluorosuccinate populated anti conformations much less frequently compared to succinate in the protein-bound simulations. However, because binding affinity is determined by the free-energy difference between the bound and unbound states, the present bound-state simulations alone cannot establish whether this altered conformational preference contributes to the experimentally observed differences in affinity (26). In conclusion, we do not yet know whether this shift of the ligand’s conformational ensemble drives the affinity loss measured by ITC. Comparative experiments of the unbound ligands and protein-ligand binding/unbinding simulations would be required to determine whether ligand conformational ensembles are a predictive determinant of fluorination tolerance.

Sensitivity of ligand fluorination to binding in SBPs is governed by a conserved chemical constraint that is nonetheless open to local and structural control. Because tolerance to partial fluorination can shift with a single aromatic substitution, the same logic could guide the design of proteins with enhanced affinity for fluorinated ligands. Given the environmental persistence of polyfluorinated compounds and the growing regulatory pressure around them, engineering SBP-based binders that exploit this tunability, while overcoming the conserved vulnerability to complete fluorination, is a concrete next step toward protein-based capture or sensing strategies for contaminants such as PFAS.

## Author contributions

P.M., B.E.C., and P.L. conceived the study. P.M. performed all the experiments. P.M., A.G., Y.A.M., P.L., and B.E.C. analyzed the ITC data. P.M., A.G., and B.E.C. analyzed the X-ray crystallography data. Y.A.M. and A.S. performed the MD simulations and analyzed the data. P.M., Y.A.M., and P.L. wrote the manuscript with input from all authors. P.L. supervised the project.

## Acknowledgements

P.L. gratefully acknowledges funding from the Okinawa Institute of Science and Technology (OIST) and COI-NEXT grant. P.M. grateful acknowledges funding from OIST’s POC program and Deutsche Forschungsgemeinschaft (DFG, German Research Foundation) under the Walter Benjamin Programme number 543786638. The authors thank OIST’s Instrument Analysis Section for providing instrument access and training. We also thank OIST’s Scientific Computing and Data Analysis Section for use of the computer clusters. We thank Prof. Assaf Vardi (Weizmann Institute of Science) for providing *Sulfitobacter* D7 DNA. We thank SPring-8 for the use of their BLXU45 beamline to perform synchrotron radiation experiments with the approval of the Japan Synchrotron Radiation Research Institute (JASRI) (proposal no. 2025A2745). We thank Prof. Nobuhiko Tokuriki and Iana Zavarzina for critical reading of the manuscript.

## Declaration of generative AI and AI-assisted technologies

In preparing this manuscript, the authors used ChatGPT (OpenAI) and Claude (Anthropic) to assist with scripting for plotting simulation data and to identify grammatical mistakes in the text. The suggestions were manually reviewed and selectively incorporated into the text. The authors take full responsibility for the content of the publication.

## Materials and methods

### Strains and plasmids

The gene encoding Sdel_0447 (20) was codon optimized and synthesized by Twist Bioscience, while the gene encoding B5M07_04440 was amplified from *Sulfitobacter* D7 genomic DNA (27), which was a gift from Prof. Assaf Vardi (Weizmann Institute of Science). These genes were cloned into a pET-28a (+) vector backbone using the In-Fusion HD Cloning Kit (Takara Bio) for expression under control of a T7 promoter with an N-terminal hexahistidine tag. The primers utilized can be found in the supporting information (Table S2). The plasmid for expression of SAR11_1361 with an N-terminal hexahistidine tag has been reported previously (19). The sequence for each gene can be found in the supporting information (Table S1).

For cloning and growth purposes, lysogeny broth (LB) medium was used containing 10 g/L tryptone, 10 g/L NaCl, 5 g/L yeast extract, and 7.5 g/L agar if needed. For protein expression, terrific broth (TB) medium was used, containing 12 g/L tryptone, 24 g/L yeast extract, 4 g/L glycerol, 25 g/L K_2_HPO_4_ and 4.62 g/L KH_2_PO_4_. Kanamycin was used at a concentration of 50 µg/mL.

### Protein expression and purification

*Escherichia coli* BL21-DE3 cells were transformed with the expression vector and incubated on LB agar with 50 µg/mL kanamycin at 30 °C. After an overnight incubation, the cells were resuspended in 2 mL of LB medium, then transferred to 500 mL of TB medium and grown at 37 °C until an OD_600_ of 0.7. Next, isopropyl-β-D-thiogalactopyranoside (IPTG) was added to a final concentration of 0.1 mM and the cultures were incubated at 17 °C for 16 h. Cells were harvested by centrifugation and stored at -20 °C.

For protein purification, frozen cultures were thawed on ice and resuspended in 100 mL of ice-cold Ni-wash buffer (pH 8, 20 mM Tris, 0.5 M NaCl, 20 mM imidazole) containing lysozyme at a concentration of 0.1 mg/mL. The resuspended cells were lysed by sonication, after which 500 U Benzonase Nuclease (Sigma-Aldrich) was added. Next, cells were centrifuged at 10,000 × g for 45 min. The supernatant was collected and loaded on a gravity-flow column containing His60 Ni Superflow Resin (Takara Bio), prewashed with Ni-wash buffer. To obtain proteins in native conditions, the column was washed twice with Ni-wash buffer and the protein eluted with Ni-elution buffer (pH 8, 20 mM Tris, 0.5 M NaCl, 0.5 M imidazole). Next, protein solutions were concentrated to approximately 400 µL using a 10 kDa molecular weight cut-off (MWCO) Amicon Ultra-4 centrifugal spin concentrator (Sigma Aldrich). Concentrated protein solutions were purified by size-exclusion chromatography using a Superdex 200 Increase 10/300 GL column (Cytiva) and eluted in DSF buffer (pH 7.50, 20 mM HEPES, 0.3 M NaCl). For crystallography, proteins were concentrated up to 10–20 mg/mL. Protein purity was evaluated by SDS-PAGE.

For protein purification under denaturing conditions, the His60 Ni Superflow Resin containing the protein was washed for 30 min with denaturing Ni-wash buffer (pH 8, 8 M urea, 20 mM Tris, 0.25 M NaCl, 20 mM imidazole) and eluted with denaturing Ni-elution buffer (pH 8, 8 M urea, 20 mM Tris, 0.25 M NaCl, 0.5 M imidazole). Denatured protein solutions were diluted to 0.5 mg/mL and stored at -20 °C for future refolding.

### Protein refolding

Approximately 25 mL of protein stored at -20 °C at a concentration of 0.5 mg/mL was thawed and transferred to a 10 kDa MWCO SnakeSkin dialysis tubing (Thermo Fisher Scientific). Proteins were dialyzed against 2 L of dialysis buffer (pH 8, 20 mM Tris, 150 mM NaCl) at 4 °C over a period of 24 h and a total of three buffer exchanges. Proteins were recovered, filtered through a 0.22 µm syringe filter, and the buffer exchanged to DSF buffer and concentrated to approximately 400 µL. The concentrated protein was purified with size-exclusion chromatography as described above.

### Isothermal titration calorimetry

A MicroCal PEAQ-ITC system (Malvern Panalytical) was used to perform ITC measurements. The protein samples were freshly purified following denaturation and refolding. Ligand and protein samples were prepared with the same batch of DSF buffer used for size exclusion chromatography to minimize extraneous heat effects. Titrations were performed at 25 °C with stirring at 750 rpm, and a reference power of 10 µcal/s. Titrations were performed through titration of 32 µM protein with 1 × 0.4 µL, then 19 × 1.8 µL injections of 280 µM ligand. Binding parameters were obtained by fitting the data to the one-set-of-sites model in MicroCal PEAQ-ITC Analysis Software. ITC experiments were performed in technical duplicates and at least three biological replicates.

### X-ray crystallography and model refinement

For the B5M07_04440-succinate complex, the protein was purified under native conditions and concentrated to 15 mg/mL as described above, and a 30-fold molar excess of succinate was then added. For the SAR11_1361-succinate and B5M07_04440-2,2-difluorosuccinate complexes, to avoid contamination of the protein with endogenous ligands bound during protein expression, the protein was refolded and purified as described above, and then a 30-fold molar excess of ligand was added. Initial screenings were carried out with the commercially available kits SG1 HT-96 Eco Screen (Molecular Dimensions) and PEGRx (Hampton Research). Droplets were created by mixing 0.5 µL of concentrated protein solution (10-15 mg/mL) with an equal volume of crystallization reagent. Droplets were prepared in 96-well sitting drop protein crystallization plates VC-1 (Az-One Corporation) and diffused against a reservoir of 50 µL crystallization reagent at 20 °C. Based on initial screens, crystal quality was optimized using the hanging-drop vapor-diffusion method. 1 µL of concentrated protein sample was mixed with 1 µL of freshly prepared crystallization reagent and equilibrated against 1 mL of reagent in 24-well VDX crystallization plates (Hampton research). Well-formed crystals were cryoprotected and flash frozen in liquid nitrogen. Crystallization and cryoprotectant conditions are listed in the supporting information (Table S7).

X-ray diffraction data was collected on the BL45XU beamline at the SPring-8 synchrotron facilities (Harima, Japan) using the automated data collection package ZOO (28) and processed using XDS (29) and KAMO (30). Each structure was solved with molecular replacement using MOLREP (31) and the initial model structure obtained from AlphaFold3 (32). Structures were refined by iterative real-space and reciprocal-space refinement in REFMAC (33) and COOT (34). Details of data collection, processing, and refinement are given in Table S3. Structures were visualized in PyMOL (21).

### Molecular dynamics simulations

The coordinates of B5M07_04440 and the ligands (succinate and 2,2-difluorosuccinate) were derived from the crystal structure data. Because no experimental structure of tetrafluorosuccinate was available, tetrafluorosuccinate was modeled by substituting the two methylene hydrogen atoms of 2,2-difluorosuccinate with fluorine atoms (Figure S12). Ligand parameters were generated with GAFF2 (35) and atomic charges derived using the standard two-stage RESP (36) procedure. Geometry optimization and electrostatic potential calculations were performed at the HF/6-31+G(d) level using Gaussian09 (37). An alternative parameterization using GAFF (38) with AM1-BCC (39) charges was evaluated as a robustness check and is described in the supporting information (Method S2). The assigned charges from both the two-stage RESP (36) and AM1-BCC fitting are included in the supporting information (Tables S5 and S6, respectively).

Protein protonation states of B5M07_04440 were assigned using the H++ webserver (40) at pH 7.5, and the protein was parameterized using the AMBER ff14SB force field (41). Conserved crystallographic waters were retained, and each complex was then solvated in a TIP3P (42) water box with dimensions of 12 Å on each side. To match the experimental conditions, Na^+^ and Cl^-^ions were subsequently added to simulate 300 mM NaCl salt concentration.

Molecular dynamics simulations were performed using AMBER 24 (43). Each system was first subjected to a three-stage energy minimization protocol with harmonic positional restraints applied to the heavy atoms of the ligand and protein residues. All minimization steps were performed with periodic boundary conditions and a 10 Å nonbonded cutoff. Long-range electrostatic interactions were treated using the Particle Mesh Ewald (PME) method. Following minimization, three independent replicates were generated by assigning different initial velocities using independent random seeds. Each system was gradually heated to 298 K under the NVT ensemble in three consecutive 100 ps stages, using Langevin dynamics with a collision frequency of 2.0 ps⁻¹. Bonds involving hydrogen atoms were constrained using the SHAKE algorithm (44). The heated system was then equilibrated under the NPT ensemble at 298 K and 1 bar. Harmonic restraints were gradually reduced from 2.0 to 1.0, and then 0.5 kcal mol⁻¹ Å⁻², followed by a final 500 ps unrestrained equilibration. Pressure during equilibration was maintained using the Berendsen weak coupling barostat with isotropic pressure scaling and a pressure relaxation time of 2.0 ps.

Production simulations were subsequently carried out under the NPT ensemble at 298 K and 1 bar, using a 2 fs integration time step for a total duration of 1 µs per replicate. Temperature was maintained using Langevin dynamics with a collision frequency of 0.5 ps⁻¹, and pressure was maintained using a Monte Carlo barostat with isotropic pressure scaling. The detailed simulation protocols are provided in the supporting information (Methods S1-4).

## Notes

### Competing Interest Statement

The authors have declared no competing interest.

