## Supporting Information for "Conserved Vulnerability, Local Tunability: How Binding Proteins Read Ligand Fluorination"

[#]These authors contributed equally.

Table S1. Gene sequences used in this study.

Table S2. Primers used in this study

Table S3. X-ray crystal data collection and refinement statistics.

Table S4. Binding parameters for protein-ligand interactions.

Table S5. Single point charge fitting table for RESP two-stage.

Table S6. Single point charge fitting table for AM1-BCC.

Table S7. Crystallization and cryoprotectant conditions.

Table S8. PFAM codes and family names for Sdel_0447, B5M07_04440, and SAR11_1361.

Figure S1. Multiple sequence alignment and percent identity matrix of Sdel_0447, B5M07_04440, and SAR11_1361.

Figure S2. Determination of binding parameters of B5M07_04440 with ITC.

Figure S3. Determination of binding parameters of SAR11_1361 with ITC.

Figure S4. Determination of binding parameters of Sdel_0447 with ITC.

Figure S5. Determination of binding parameters of B5M07_04440 Tyr42Ala with ITC.

Figure S6. Determination of binding parameters of SAR11_1361 Phe187Ala with ITC.
Figure S7. Control ITC experiments.
Figure S8. Results from MD simulations of B5M07_04440 using AM1-BCC charges and the GAFF force field.
Figure S9. Protein-ligand distances used to characterize the bound state in simulation of B5M07_04440 using AM1-BCC charges and the GAFF force field.
Figure S10. Protein–ligand distances used to characterize the bound state of B5M07_04440 in simulations using RESP charges and the GAFF2 force field.
Figure S11. Protein–ligand distances between the ligand H/F atoms and the aromatic ring of Tyr42 in MD simulations of B5M07_04440.
Figure S12. Ligand poses used to prepare the B5M07_04440-ligand MD simulation systems.
Figure S13. RMSD of the C-domain of B5M07_04440 in the tetrafluorosuccinate complex during MD simulations.
Figure S14. Aromatic residues in the binding pocket are conserved in SBPs.
Figure S15. Water molecules at the ligand binding site of B5M07_04440 and SAR11_1361.
Figure S16. The conformation of succinate bound in B5M07_04440 vs SAR11_1361.
Method S1. Protein-ligand system preparation for MD simulations of B5M07_04440.
Method S2. Ligand parameterization for MD simulations of B5M07_04440.
Method S3. System preparation for MD simulations of B5M07_04440.
Method S4. Molecular dynamics simulations of B5M07_04440.

**Table S1. Gene sequences used in this study.** Start codon highlighted in blue, hexahistidine tag in green, and position of mutation in red.

| Gene name | DNA sequence |
|---|---|
| B5M07_04440 | ATGGGCAGCAGCCATCATCATCATCATCACAGCAGCGGCCAGGACGTTGATAAATCC<br>GACTGGCCCAGCAGCTTTACCGTCGGCACAGCCAGCCAAGGCGGCACCTACTTCGCC<br>TATGGCTCCGGCTGGGCGAACCTCGTGGCCAAAGAACTGGGCCTGACAGGCGGCGGC<br>GAAGTCACCGGCGGCCCGATGCAGAACATGGCGCTGGTCCACACCGGCGAGGCGGCC<br>TTCGGCATGACCACGATGGGTCCGGCGGCGGAATCGCTGGCTGGCACCAACCCCATC<br>GCACCCGGTCTGGAAATGACCCAAGCCTGCGCGATGTTCCCGATGTACCAGACACCT<br>TTCTCGGTGACTGCGCTTGCGTCCTCGGGCATCGAAACCGTGGCTGACATTCCGGCA<br>GGCGCCAAGATCGGCTTTGGCCCGGCAGGCTCCACTTCTGACACTTATTTCCCGCGC<br>ATGATGGACACGCTCGGCGTGGAATATGAGCGCCGCAACGGCGGTTGGACCGATCTG<br>GGTGGCCAGTTGCAGGATGGTCTGCTGGACGTGATCGCTTTTGCCGCAGGCGTGCCT<br>GTGCCCGCGGTCAGCCAGTTGGAAGTGCAGACCGACGTGAACATCATCGAGTTCACC<br>GAGGAAGAGCAGGCCAAACTGCTCGAAGAATTCCCGGTATCTCAGTTCGACATCGCA<br>TCCAGCACCTACACCACGCTTGAAGCAGATGCGCGATCGGTCTCCATGTGGAACTTC<br>GCCATCGCCAACTGCGATCTGCCGGAATCCTTCGTCAAAGCGGCTGTGGATGTGGTC<br>ATGTCCGACAATGAGCGGATGAAGGGCATTCACAAGGCGGCTGCGTCCTCCCTGCCT<br>GAGAACTGGGACAAGAACAAAGTTCTGAAATGGCACCCCGGTGCGGCGGCATGGTTC<br>AAAGAGAACGCGGATGCCGATATTTCCGACGATATGATCTACGGTAACTAA |
| SAR11_1361 | ATGGGCAGCAGCCATCATCATCATCATCACAGCAGCGGCGAGAACCTGTATTTTCAG<br>AGCGGCAGCAGCCATATGAAAGTGGTGGGCGATAAAATCATTCTGGGTGCAGCAATT<br>AGCCTGACCGGTAAATATAGCAGCAATGGTGTTCATACGCAGAATGGTTATAATCTG<br>GCAGTGGATCGCATTAATGATATGGGTGGTATTAAAGTTGGTGGCAAGACCTATAAA<br>TTCGAGATCATCTATTATGATGATGAAAGCAATAGCGGTCGTGCAGCACAGCTGGCA<br>GAACGTCTGATTCAGCAGGATGGTGTGCAGTATATGCTGGGTCCGTATAGCAGCGGT<br>CTGACCAAAGCAATGGCACCGGTTACCGAAAAATATGGTATTCCGATGGTTGAAGCA<br>AATGGTGCAAGCCGTAGCCTGTTTACCAAAGGCTATAAATACCTGTTTGCAGTTCTG<br>GCACCGGCAAATCTGTATCTGGATGTTGCAATTGATCTGGCCGTTGAAAAAAATGGT<br>GGTAAACCGGTTAAAATCGCCATGGCCTTTGAACAGGATGCATTTAGCCAGGATGTT<br>CGTCTGGGTATTCTGGATGCCGCAAAACGTACCGGTAGCAAAATTATCATTGATGAC<br>AAACTGCCGAAAGAACTGAACGATATGGCAGCAACCCTGAGCAAAGTTAAAGCAGTT<br>AAACCGGATGTTCTGGTTGTTAGCGGTCATACCAAAGGTGCACTGACCGCAGTTCGT<br>CAGATTAGCGAAATGAAAGTTGATGTTCCGATGCTGGCAATGACCCATTGTGATGCA<br>GCCAAACTGAGCAAAATGCATGGTAAAGCAAGCGAATATGCACTGTGTGCAAGCCAG<br>TGGCATAAAAGCCTGAGCTATAAAGATGACTTCTTCAAAGATGGCACCACCTATGAT<br>GCAGATTTTACAAAAGCATTTGGTTATGCACCGCCTTATCAGGCAGCAGAAAGCAGC<br>GCAGCACTGCTGGTTTTTAAAGATGCATTTGAACGTGCACAGAGCTTCGATACCAAA<br>AAAGTTCGTGATGCACTGGCAGCCACAGATATGCAGACCTTTTATGGCAATATCAAA<br>TTTGCCGAAGGTGGTCAGAATGTTAGCAAACCGATGGTTCTGTTTCAGGTTAGCTGT<br>AATGATGATGGTAGCAAATGCGAAAACAAAGTTGTTGCACCGACCAAATGGGCCAGC<br>GCAGAACTGGTTCATCCGATTCCGAGCTGGTCAAAACGTTAA |

| | |
|---|---|
| Sdel_0447 | ATGGGCAGCAGCCATCATCATCATCATCACAGCAGCGGCGAGAACCTGTATTTTCAG<br>AGCGGCAGCAGCGCAGAGTATACGATCAAAGTCACCCATGTAGTGTCTCCTAACACG<br>CCGAAAGGTAAAGGGGCGGATTTTTTCGCCAAACGCGTTGGGGAGTTGACAAATGGA<br>AAAGTTGAAGTGATCGTCTTTCCAAATAGTCAACTTTATGGCGATGGTGAAGAAATG<br>AAGGCTTTAAAACTGGGCAATGCCCACATTGCGATGCCATCTTTCTCAAAATTTACT<br>TCTCTGGTGCCAGAAATGCAGCTCTTTGATCTGCCCTTTATATTTCGCGACAAGGAC<br>CATCTGTACAAAGTTCTCGATGGTGAAGTGGGTCAAATATTAAAGGATAAAGTTTCT<br>AAGAAGGGTTTCGTTGCCCTGGATTACTGGGATGCAGGCTTTAAACATCTTTCCTCA<br>AATAAAAAACCGATCTTACTGCCCGAAGATGCAGCAGGGCAGAAGTTTCGCATTATG<br>AGCAGCCATGTTCTCGAAGCCCAATTTAAAGCCGTGGGTGCAAATCCGCAAGTCCTG<br>CCATTCAGTGAAGTCTACAGTGCCCTTCAACAGGGAGTTGTTGATGGCGCGGAAAAC<br>CCGCTGTCGAATTTCTATACTAAGAAATTTAATGAAGTGCAGACGGATCTTACACTG<br>TCGAACCATGGGTATCTTGGTTACCTGGTTATTATGTCTGAGTCCTTTTGGAAGAAA<br>TTTCCCAAGGATCTTAAGCCAATGGTTCTGCAGGCCATGAAGGAAGCCACAGAGTAT<br>GAACGGAAAGAAGCAGCACTTGACGATGAGGACATGTTAGCTAAAATCTCGGAGTAT<br>GCGAAGGCGAGCGGTAATCTGAAAATCCATACATTGACACCCGAGCAAAAGGCAGCA<br>TGGCAGAAAGCCATGGAGGCCATATATCCTCAGTTTTACAAAACCATCGGAGAAGAT<br>CTTATTAAGAAAGTCCAGGCCGTCAAGTAA |

**Table S2. Primers used in this study**.

| **Primer name** | **Sequence** |
|---|---|
| Sdel_0447 vector preparation forward | GAATTCGAGCTCCGTCGACAAG |
| Sdel_0447 vector preparation reverse | GCTGCTGCCGCTCTGAAA |
| Sdel_0447 gene preparation forward | CAGAGCGGCAGCAGCGCAGAGTATACGATC |
| Sdel_0447 gene preparation reverse | CCAGAGTAGTTACTTGACGG |
| B5M07_04440 Tyr42Ala forward | CGAAGGCGGTGCCGCCTTGGCTGG |
| B5M07_04440 Tyr42Ala reverse | GGCGGCACCGCCTTCGCCTATGGCT |
| SAR11_1361 Phe187Ala forward | GGATGCAGCCAGCCAGGATGTTCGTC |
| SAR11_1361 Phe187Ala reverse | TCCTGGCTGGCTGCATCCTGTTCAA |

**Table S3. X-ray crystallography data collection and refinement statistics.**

| **Structure** | **B5M07_04440 succinate** | **B5M07_04440 2,2-difluorosuccinate** | **SAR11_1361 succinate** |
|---|---|---|---|
| PDB code | 48GN | 48GB | 48FW |
| **Data collection** | | | |
| Space group | $P\ 1\ 2_1\ 1$ | $P\ 2_1\ 2_1\ 2_1$ | $P\ 1\ 2_1\ 1$ |
| Cell dimensions<br>$a$, $b$, $c$ (Å)<br>α, β, γ (°) | <br>38.0, 69.8, 113.4<br>90.0, 90.0, 90.0 | <br>67.1, 68.4, 116.8<br>90.0, 90.0, 90.0 | <br>46.2, 118.8, 67.2<br>90.0, 90.0, 90.0 |
| Resolution (Å) | 44.05–1.36 (1.44–1.36) | 47.96–1.84 (1.95–1.84) | 46.26–1.52 (1.61–1.52) |
| $R_{merge}$ (%) | 8.2 (112.1) | 13.9 (134.4) | 11.2 (81.6) |
| $CC_{1/2}$ (%) | 99.8 (51.7) | 99.8 (53.8) | 99.3 (57.5) |
| $I / \sigma(I)$ | 11.75 (1.62) | 9.68 (1.13) | 7.89 (1.64) |
| Completeness (%) | 99.7 (99.3) | 99.8 (99.0) | 98.9 (98.1) |
| Redundancy | 3.49 (3.50) | 6.28 (6.15) | 2.94 (2.84) |
| **Refinement** | | | |
| Resolution (Å) | 44.05–1.36 | 47.96–1.84 | 46.26–1.52 |
| No. unique reflections | 127812 | 47508 | 111215 |
| $R_{work}$ / $R_{free}$ (%) | 12.0 / 14.6 | 18.3 / 22.2 | 15.9 / 19.0 |
| Number of atoms (chain A / chain B)<br>Protein<br>Ligand<br>Ion<br>Water<br>Other | <br>2269 / 2281<br>8 / 8<br>2<br>537<br>72 | <br>2233 / 2250<br>10 / 10<br>2<br>212<br>- | <br>2871 / 2880<br>8 / 8<br>-<br>589<br>- |
| $B$-factors (Å$^2$) (chain A / chain B)<br>Protein<br>Ligand<br>Ion<br>Water<br>Other | <br>15.9 / 15.9<br>8.5 / 8.5<br>18.3<br>31.7<br>14.6 | <br>32.5 / 34.5<br>30.3 / 31.7<br>36.5<br>36.2<br>- | <br>16.4 / 16.5<br>11.7 / 11.5<br>-<br>27.6<br>- |
| Root mean square deviations<br>Bond lengths (Å)<br>Bond angles (°) | <br>0.011<br>1.76 | <br>0.007<br>1.56 | <br>0.010<br>1.76 |

**Table S4. Binding parameters for protein-ligand interactions.** For each protein-ligand interaction, the active protein (n), dissociation constant ($K_d$), and binding enthalpy (ΔH) are given. For each interaction, at least three biological replicates and two technical replicates were performed, which were combined with inverse-variance weighting. Mean values ± standard deviation are given.

| Protein | Ligand | n | $K_d$ (nM) | ΔH (kcal/mol) |
|---|---|---|---|---|
| **B5M07_04440** | Succinate | 0.77 ± 0.17 | < 10 | -4.40 ± 0.32 |
| | 2,2-Difluorosuccinate | 0.76 ± 0.05 | < 10 | -4.99 ± 0.48 |
| | Tetrafluorosuccinate | 0.72 ± 0.16 | 799 ± 561 | -1.73 ± 0.33 |
| | Fumarate | 0.74 ± 0.12 | < 10 | -3.36 ± 0.36 |
| | Fluorofumarate | 0.84 ± 0.33 | < 10 | -3.72 ± 0.47 |
| | Difluorofumarate | 1.06 ± 0.06 | 330 ± 131 | -1.31 ± 0.12 |
| | Glutarate | No binding | | |
| **SAR11_1361** | Succinate | 0.76 ± 0.12 | 31.1 ± 22.3 | -12.4 ± 0.38 |
| | 2,2-Difluorosuccinate | 0.85 ± 0.18 | 89.2 ± 8.07 | -10.0 ± 0.37 |
| | Tetrafluorosuccinate | 0.72 ± 0.11 | 1011 ± 98.2 | -8.44 ± 1.06 |
| | Fumarate | 0.61 ± 0.16 | 46.0 ± 12.0 | -12.4 ± 0.56 |
| | Fluorofumarate | 0.84 ± 0.01 | 86.2 ± 33.8 | -12.1 ± 0.72 |
| | Difluorofumarate | 1.01 ± 0.04 | 19.5 ± 6.56 | -10.7 ± 0.08 |
| | Glutarate | 0.70 ± 0.16 | 18.3 ± 4.41 | -10.5 ± 0.27 |
| | 2,2-Difluoroglutarate | 0.78 ± 0.19 | 38.6 ± 16.9 | -9.62 ± 0.81 |
| | Hexafluoroglutarate | No binding | | |
| **Sdel_0447** | Succinate | 0.83 ± 0.03 | 329 ± 68.2 | -3.86 ± 0.18 |
| | 2,2-Difluorosuccinate | 0.93 ± 0.07 | 381 ± 53.0 | -4.13 ± 0.44 |
| | Tetrafluorosuccinate | No binding | | |

| | | | | |
|---|---|---|---|---|
| | Fumarate | 0.77 ± 0.02 | 174 ± 44.1 | -4.60 ± 1.13 |
| | Fluorofumarate | 1.13 ± 0.23 | 466 ± 206 | -2.92 ± 1.15 |
| | Difluorofumarate | 1.07 ± 0.14 | 721 ± 616 | -2.18 ± 0.23 |
| | Glutarate | No binding | | |
| **B5M07_04440 Tyr42Ala** | Succinate | 0.86 ± 0.13 | 158 ±107 | -4.25 ± 0.55 |
| | 2,2-Difluorosuccinate | 0.73 ± 0.17 | 1488 ±169 | -5.15 ± 0.49 |
| | Tetrafluorosuccinate | 0.54 ± 0.17 | 1868 ± 1003 | -1.70 ± 0.19 |
| | Fumarate | 0.95 ± 0.09 | 628 ± 460 | -2.57 ± 0.39 |
| | Fluorofumarate | 0.95 ± 0.37 | 298 ± 148 | -3.70 ± 0.95 |
| | Difluorofumarate | 0.81 ± 0.18 | 912 ± 610 | -3.29 ± 1.13 |
| **SAR11_1361 Phe187Ala** | Succinate | 0.73 ± 0.11 | 646 ± 36.9 | -4.70 ± 0.05 |
| | 2,2-Difluorosuccinate | 0.89 ± 0.11 | 175 ± 59.2 | -9.00 ± 0.62 |
| | Tetrafluorosuccinate | 0.85 ± 0.10 | 193 ± 60.5 | -8.09 ± 0.38 |
| | Fumarate | 0.67 ± 0.08 | 1493 ± 455 | -5.56 ± 0.52 |
| | Fluorofumarate | 0.72 ± 0.08 | 1830 ± 430 | -6.20 ± 0.10 |
| | Difluorofumarate | 0.77 ± 0.04 | 1466 ± 894 | -5.29 ± 0.92 |
| | Glutarate | 0.74 ± 0.15 | 1048 ± 870 | -3.50 ± 0.45 |
| | 2,2-Difluoroglutarate | 0.82 ± 0.12 | 1413 ± 608 | -3.59 ± 0.99 |
| | Hexafluoroglutarate | No binding | | |

**Table S5. Single point charge fitting table for RESP two-stage.**

| No. | **Succinate** | | **2,2-Difluorosuccinate** | | **Tetrafluorosuccinate** | |
|---|---|---|---|---|---|---|
| | Atom type | Charge | Atom type | Charge | Atom type | Charge |
| 1 | c | 0.895265 | c3 | 0.012573 | c3 | 0.328645 |
| 2 | o | -0.919321 | o | -0.909495 | o | -0.853253 |
| 3 | c3 | 0.034599 | c | 0.931266 | c | 0.912489 |
| 4 | o | -0.919321 | o | -0.909495 | o | -0.853253 |
| 5 | c3 | 0.034599 | c3 | 0.578862 | c3 | 0.328645 |
| 6 | o | -0.919321 | o | -0.881106 | o | -0.853253 |
| 7 | c | 0.895265 | c | 0.862679 | c | 0.912489 |
| 8 | o | -0.919321 | o | -0.881106 | o | -0.853253 |
| 9 | hc | -0.04561 | f | -0.358842 | f | -0.267314 |
| 10 | hc | -0.04561 | f | -0.358842 | f | -0.267314 |
| 11 | hc | -0.04561 | hc | -0.043247 | f | -0.267314 |
| 12 | hc | -0.04561 | hc | -0.043247 | f | -0.267314 |

**Table S6. Single point charge fitting table for AM1-BCC.**

| No. | Succinate | | 2,2-Difluorosuccinate | | Tetrafluorosuccinate | |
|---|---|---|---|---|---|---|
| | Atom type | Charge | Atom type | Charge | Atom type | Charge |
| 1 | c | 0.8966 | c3 | -0.2124 | c3 | 0.2426 |
| 2 | o | -0.8913 | o | -0.8738 | o | -0.8193 |
| 3 | c3 | -0.1564 | c | 0.9106 | c | 0.9136 |
| 4 | o | -0.8913 | o | -0.8738 | o | -0.8193 |
| 5 | c3 | -0.1564 | c3 | 0.2906 | c3 | 0.2426 |
| 6 | o | -0.8913 | o | -0.8388 | o | -0.8193 |
| 7 | c | 0.8966 | c | 0.8956 | c | 0.9136 |
| 8 | o | -0.8913 | o | -0.8388 | o | -0.8193 |
| 9 | hc | 0.0212 | f | -0.2753 | f | -0.2593 |
| 10 | hc | 0.0212 | f | -0.2753 | f | -0.2593 |
| 11 | hc | 0.0212 | hc | 0.0462 | f | -0.2593 |
| 12 | hc | 0.0212 | hc | 0.0462 | f | -0.2593 |

**Table S7. Crystallization and cryoprotectant conditions.**

| Structure | Crystallization conditions | Cryoprotectant conditions |
|---|---|---|
| B5M07_04440 with succinate | 1 µL of 15 mg/mL protein and 1 µL of 28% (w/v) PEG 3350, 0.1 M sodium acetate, 0.1 M MES (pH 5.5) | 30% (v/v) ethylene glycol, 19.6% (w/v) PEG 3350, 0.07 M sodium acetate, 0.07 M MES (pH 5.5) |
| B5M07_04440 with 2,2-difluoro-succinate | 1 µL of 10 mg/mL protein and 1 µL of 40% (w/v) PEG 3350, 0.1 M Bis-Tris (pH 6) | 30% (v/v) PEG 400, 28% (w/v) PEG 3350, 0.07 M Bis-Tris (pH 6) |
| SAR11_1361 with succinate | 1 µL of 10 mg/mL protein and 1 µL of 1% PEG 3350, 50% (v/v) Tacsimate (pH 4), 0.1 M Bis-Tris propane (pH 9) | 30% (v/v) PEG 400, 0.7% PEG 3350, 35% (v/v) Tacsimate (pH 4), 0.07 M Bis-Tris propane (pH 9) |

**Table S8. PFAM codes and family names for Sdel_0447, B5M07_04440, and SAR11_1361.** PFAM codes and family names were identified using InterProScan (1).

| **Protein** | **PFAM code** | **Family name** |
|---|---|---|
| Sdel_0447 | PF03480 | Bacterial extracellular solute-binding protein, family 7 |
| B5M07_04440 | PF16868 | NMT1-like family |
| SAR11_1361 | PF13458 | Periplasmic binding protein |

**(a)**

```
Sdel_0447        MTKFLKV---MAAATLLVSSAIAAEYTIKVTHVVSPNTPKGKGADFFA------------
B5M07_04440      -MKFTAI---LAASTILAGAAAAQDVDKSDWPSSFTVGTASQGGTYFAYG----SGWANL
SAR11_1361       -MKIRNIIITLVSAIALLISASTISFAKVVGDKIILGAAISLTGKYSSNGVHTQNGY-NL
                   *:  :   :.::  *  :* : .             .  .  . : :

Sdel_0447        --KRVGELTNGKVEVIVFPNSQLYGDGEEMKALK----------------LGNAHIAMPS
B5M07_04440      VAKELGLTGGGEVTGGPMQNMALVHTGEAAFGMTT---------------MGPAAESLAG
SAR11_1361       AVDRINDMGGIKVGGKTYKFEIIYYDDESNSGRAAQLAERLIQQDGVQYMLGPYSSGL--
                   . :.   . :*         :   .*   .                  :*    .:

Sdel_0447        FSKFTS-----LVPEMQL-----------FDLPFIFRDKDHLYKVLDGEV------GQIL
B5M07_04440      TNPIAP-----GLEMTQAC----AMFPM-YQTPFSVTALASSGIETVADI--------PA
SAR11_1361       TKAMAPVTEKYGIPMVEANGASRSLFTKGYKYLFAVLAPANLYLDVAIDLAVEKNGGKPV
                  . ::.      :   :            :.  * .            ::

Sdel_0447        KDKVS-----------------KKGFVALDYWDAGFKHLSSNKKPILLPEDAAGQKFRIM
B5M07_04440      GAKIGFGPAGSTSDTYFPRMMDTLGV-EYERRNGGWTDLGGQLQDGLLDVIAFAAGV---
SAR11_1361       KIAMAFEQDAFSQDV-------RLGILDAAKRTGSKIIDDKLPKELNDMAATLSKVKAV
                    :.                   *.     . ..   :..:    *    *    .

Sdel_0447        SSHVLEAQFKAVGANPQVLPFSEVYSALQQGVVDGAENPLSNFYTKKFNEVQTDLTLSNH
B5M07_04440      -------------PVPAVSQL-EVQTDVN--IIEFTEEEQAKL-LEEF-PV-SQFDIASS
SAR11_1361       KPDVLVVSGHTKGALTAVRQISEMKVDVP--MLAMTHCDAAKL-SKMHGKA-SEYALCAS
                              . . *  : *:   :   ::  :    :::  : .  . ::  :.

Sdel_0447        GYLGYLVIMSESFW--KKFPKD--------------LKPMVLQAMKEATEYERKEA----
B5M07_04440      TY--------------TTLEAD--------------ARSVSMWNF---------------
SAR11_1361       QWHKSLSYKDDFFKDGTTYDADFTKAFGYAPPYQAAESSAALLVFKDAFERAQSFDTKKV
                  :              ..   *                .  :  :

Sdel_0447        --ALDDEDMLAKISEYAKASGNLKI--------HTLTPEQKAAWQKAMEAIYPQFYKTIG
B5M07_04440      --AIANCDLPE-----SFVKAAVDV--------VMSDNERMKGIHKAAASSLPENWDK--
SAR11_1361       RDALAATDMQTFYGNIKFAEGGQNVSKPMVLFQVSCNDDGSKCENKVVA---PTKWAS--
                   *:   *:         ...  .:             :     :*.     *  : .

Sdel_0447        EDLIKKVQAVK-------------------
B5M07_04440      NKVLKKWHPGAAAWFKENADADISDDMIYGN
SAR11_1361       AELV--HP-IPSWSKR--------------
                  .::
```

**(b)**

| | Sdel_0447 | B5M07_04440 | SAR11_1361 |
|---|---|---|---|
| **Sdel_0447** | 100% | 17% | 16% |
| **B5M07_04440** | 17% | 100% | 27% |
| **SAR11_1361** | 16% | 27% | 100% |

**Figure S1. Multiple sequence alignment and percent identity matrix of Sdel_0447, B5M07_04440, and SAR11_1361.** (a) Multiple sequence alignment of Sdel_0447 (UniProt ID:

D1AZL7), B5M07_04440 (UniProt ID: A0ABW1YWG3), and SAR11_1361 (UniProt ID: Q4FKX4) generated using MUSCLE (3.8) and (b) the corresponding percent identity matrix (2).

#### (a) B5M07_04440 + Succinate*

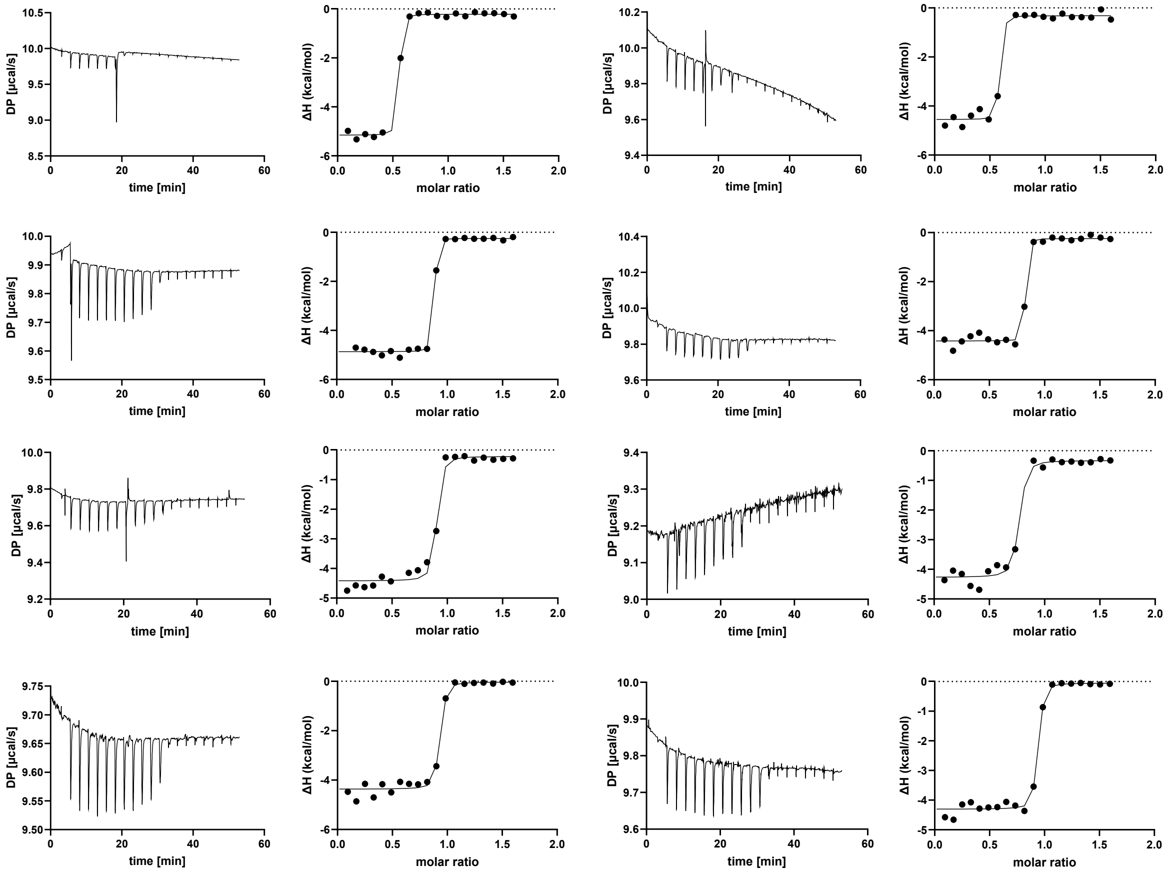


#### (b) B5M07_04440 + 2,2-Difluorosuccinate*

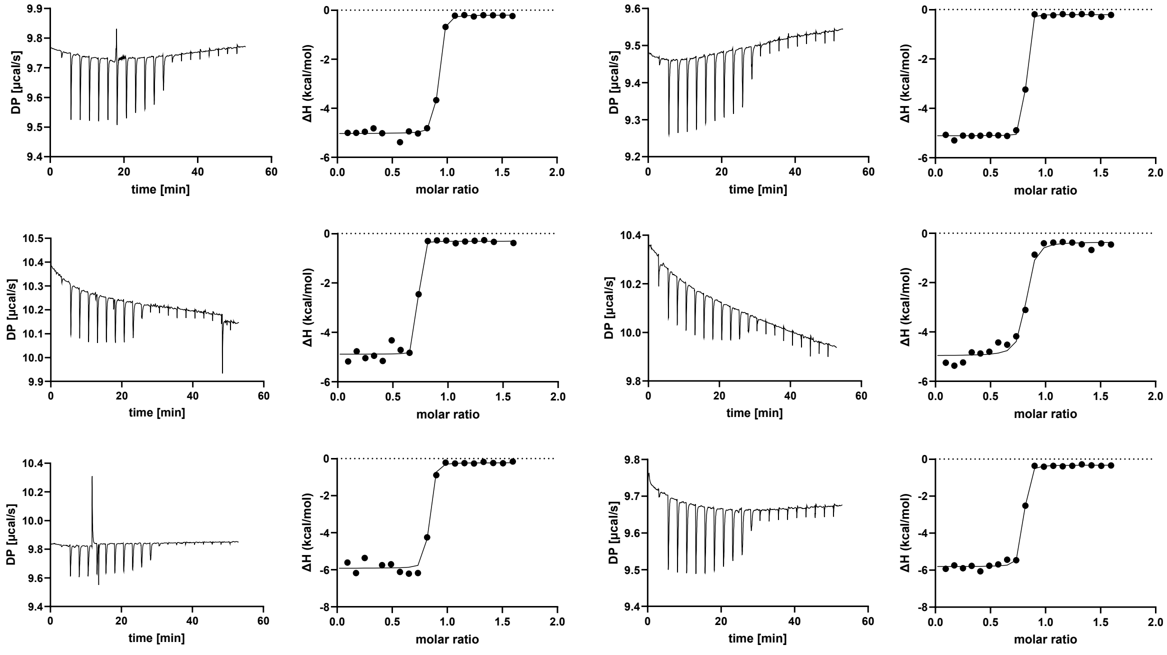

#### (c) B5M07_04440 + Tetrafluorosuccinate

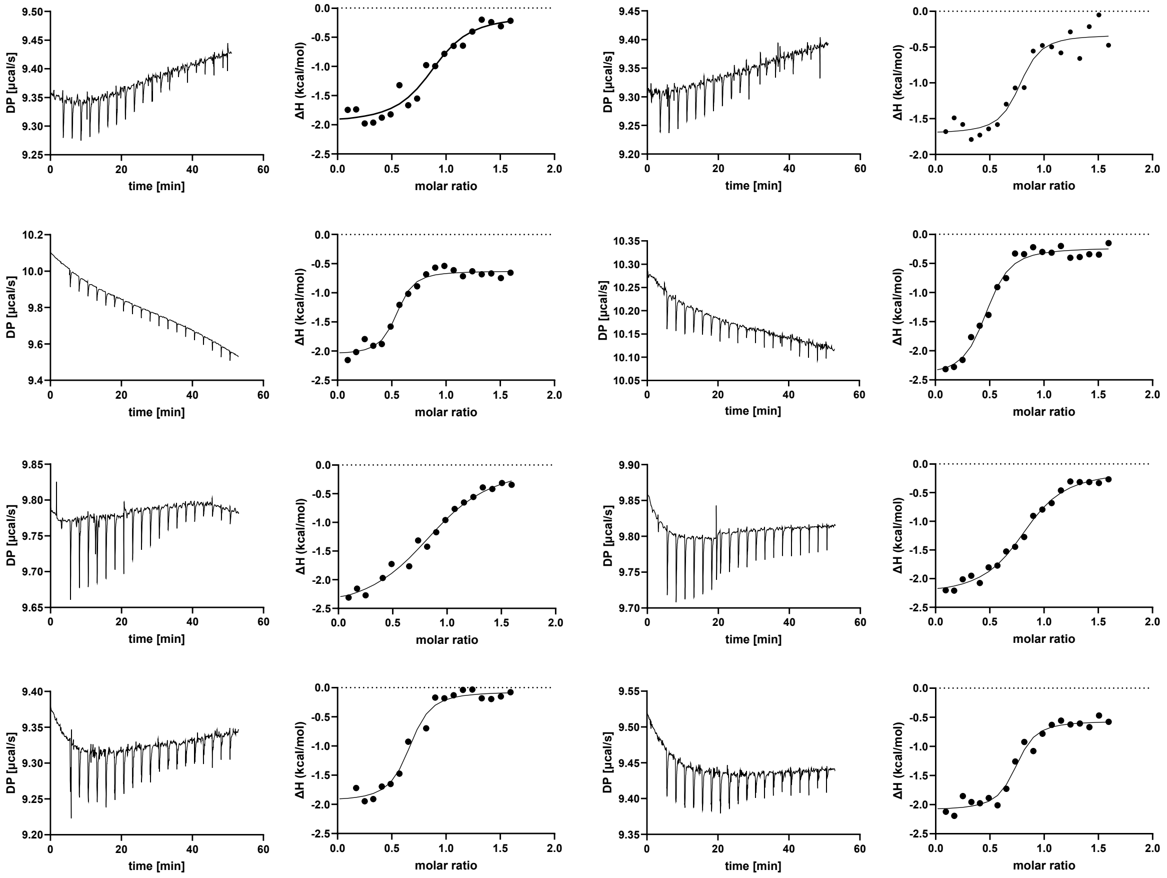


#### (d) B5M07_04440 + Fumarate*

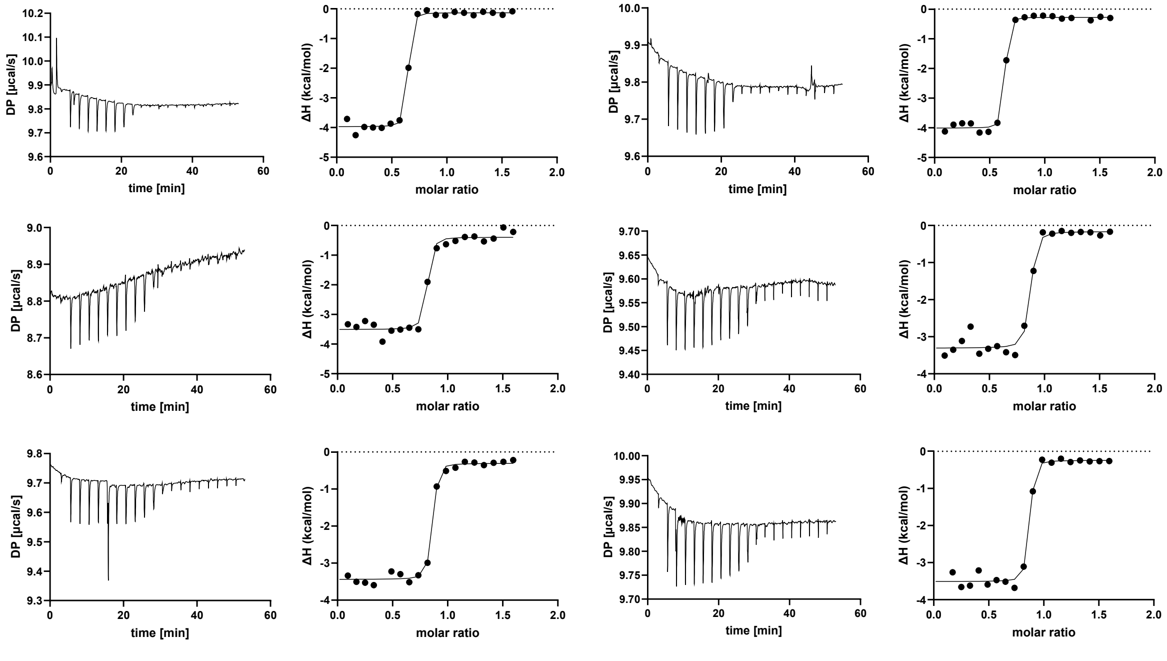

#### (e) B5M07_04440 + Fluorofumarate*

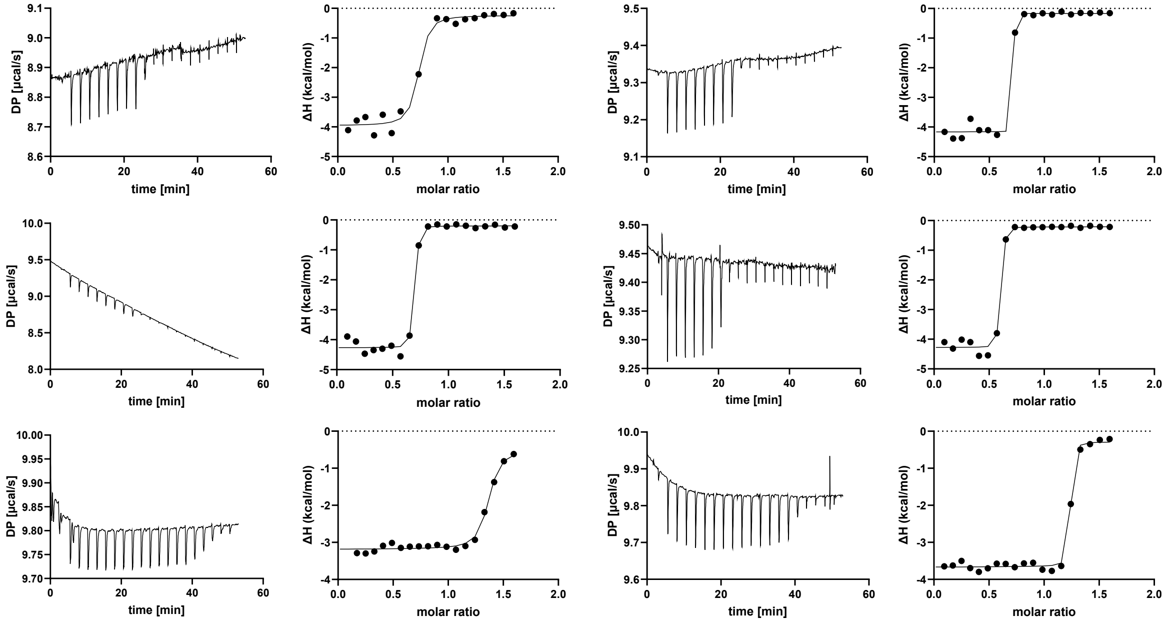


#### (f) B5M07_04440 + Difluorofumarate

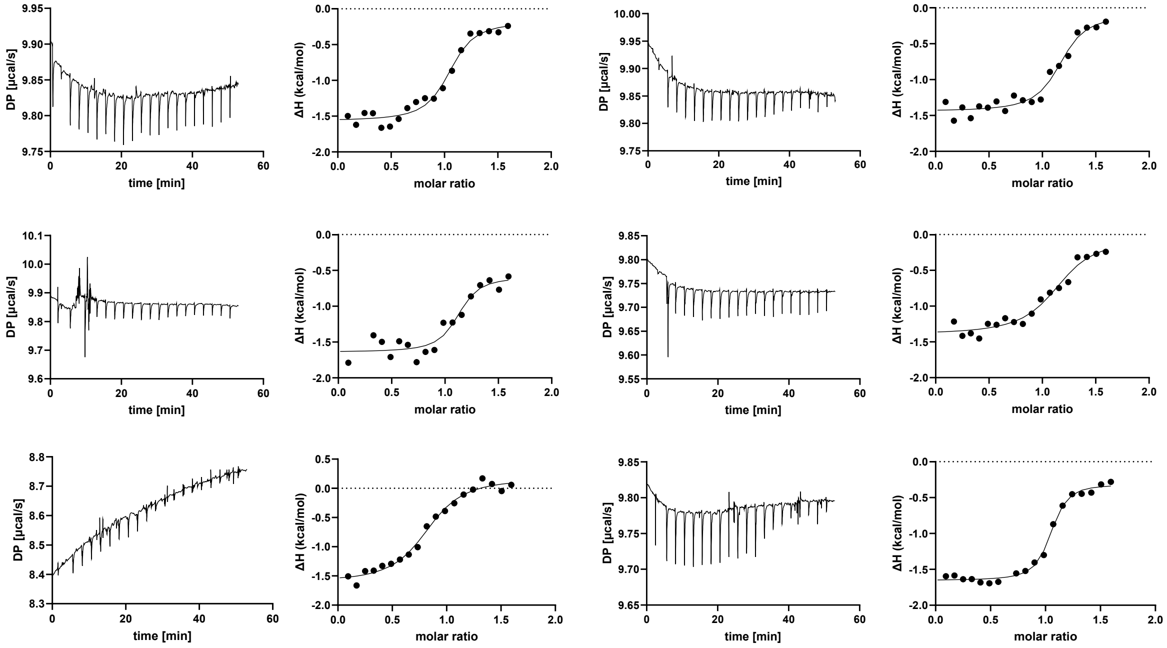

#### (g) B5M07_04440 + Glutarate

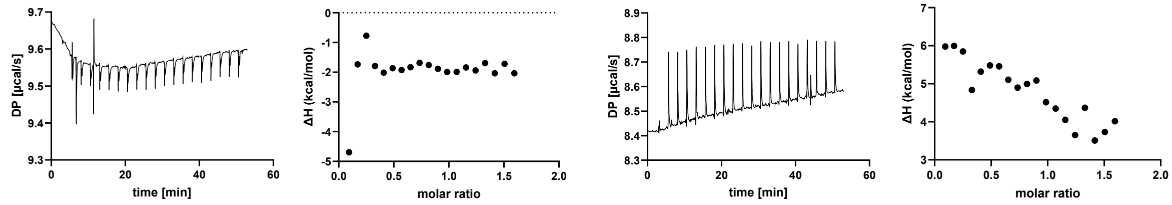


**Figure S2. Determination of binding parameters of B5M07_04440 with ITC.** Protein samples were denatured, refolded, purified, concentrated to 32 µM, and titrated with freshly prepared ligands at a concentration of 280 µM. $K_d$ values were measuring using ITC with the following ligands: (a) succinate, (b) 2,2-difluorosuccinate, (c) tetrafluorosuccinate, (d) fumarate, (e) fluorofumarate, (f) difluorofumarate, (g) glutarate. Shown is the raw and integrated data for at least three biological replicates and technical duplicates, with each row constituting a biological replicate. Asterisks mark the ligands for which the binding affinity was above the limit of quantitation of ITC, and for which the $K_d$ is reported as <10 nM.

#### (a) SAR11_1361 + Succinate

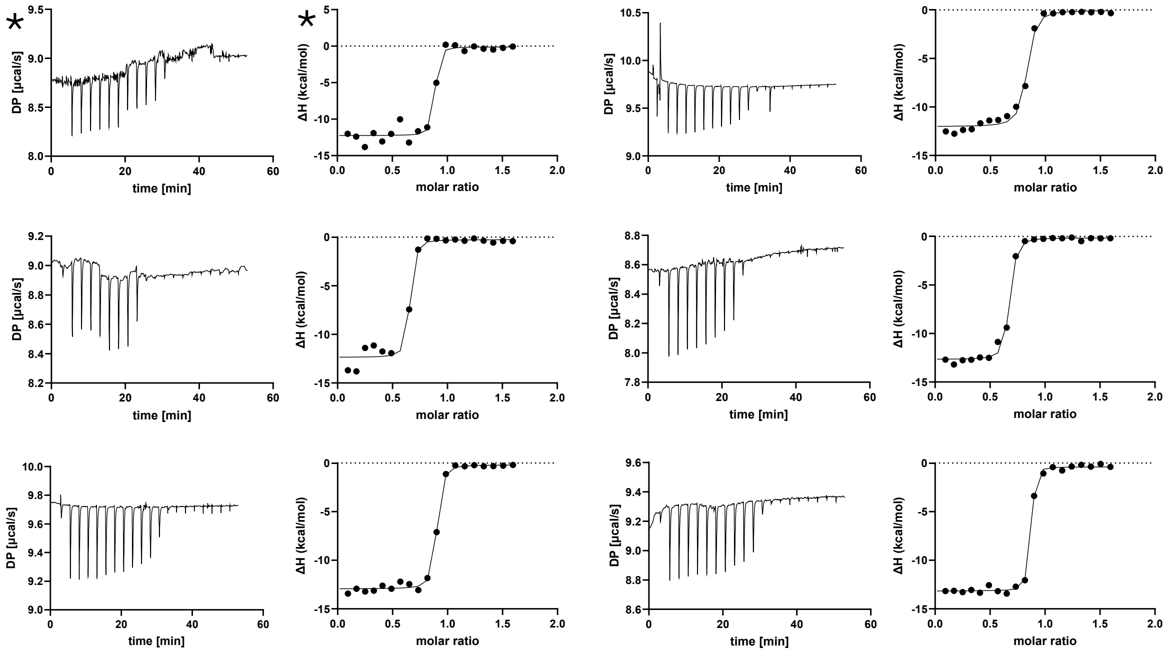


#### (b) SAR11_1361 + 2,2-Difluorosuccinate

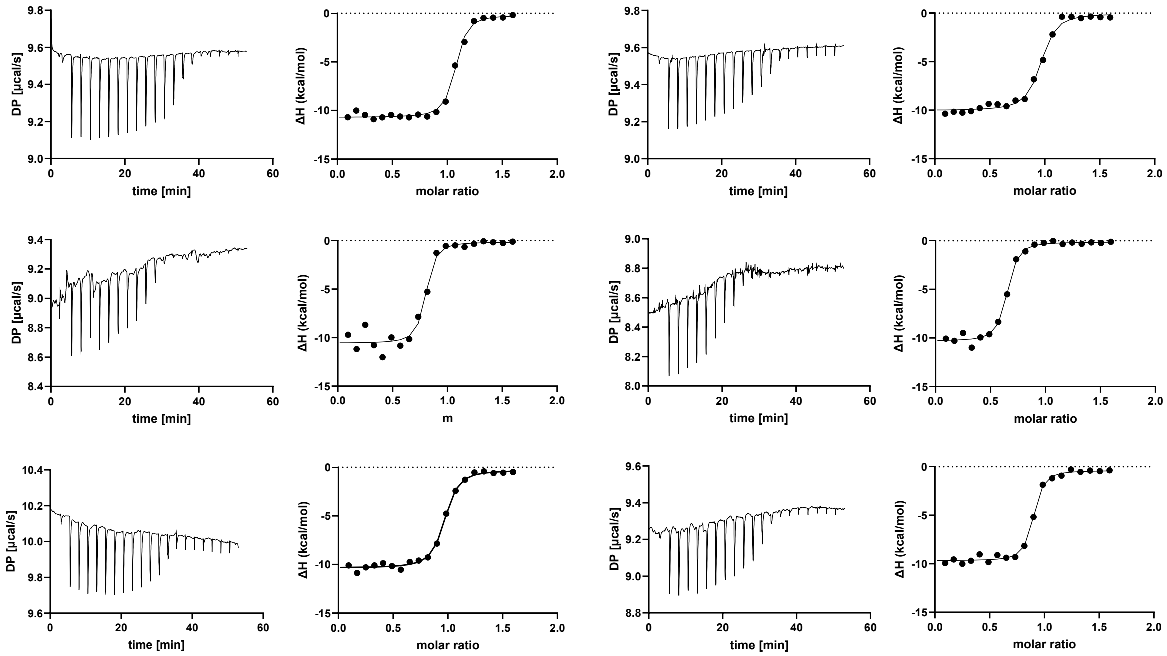

#### (c) SAR11_1361 + Tetrafluorosuccinate

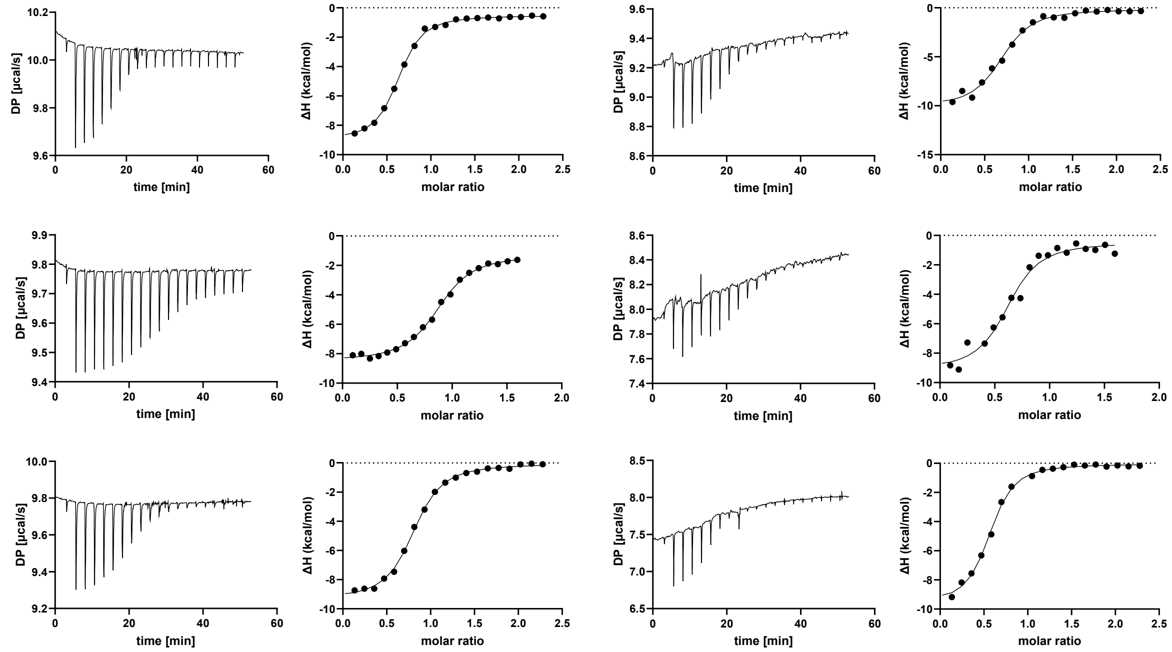


#### (d) SAR11_1361 + Fumarate

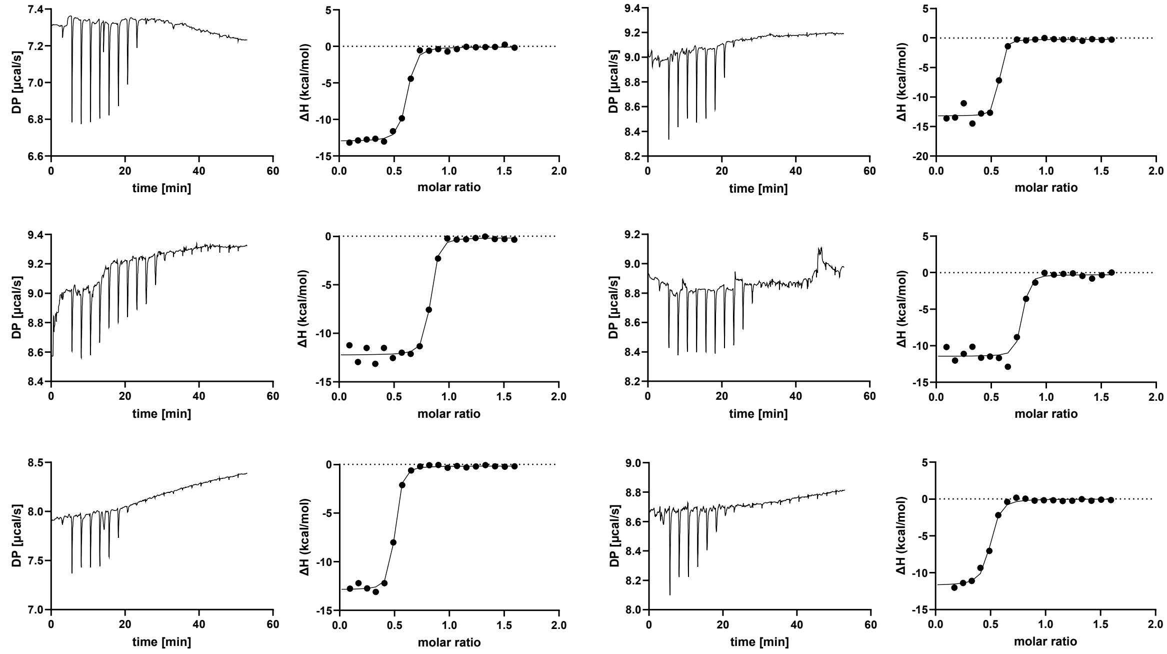

#### (e) SAR11_1361 + Fluorofumarate

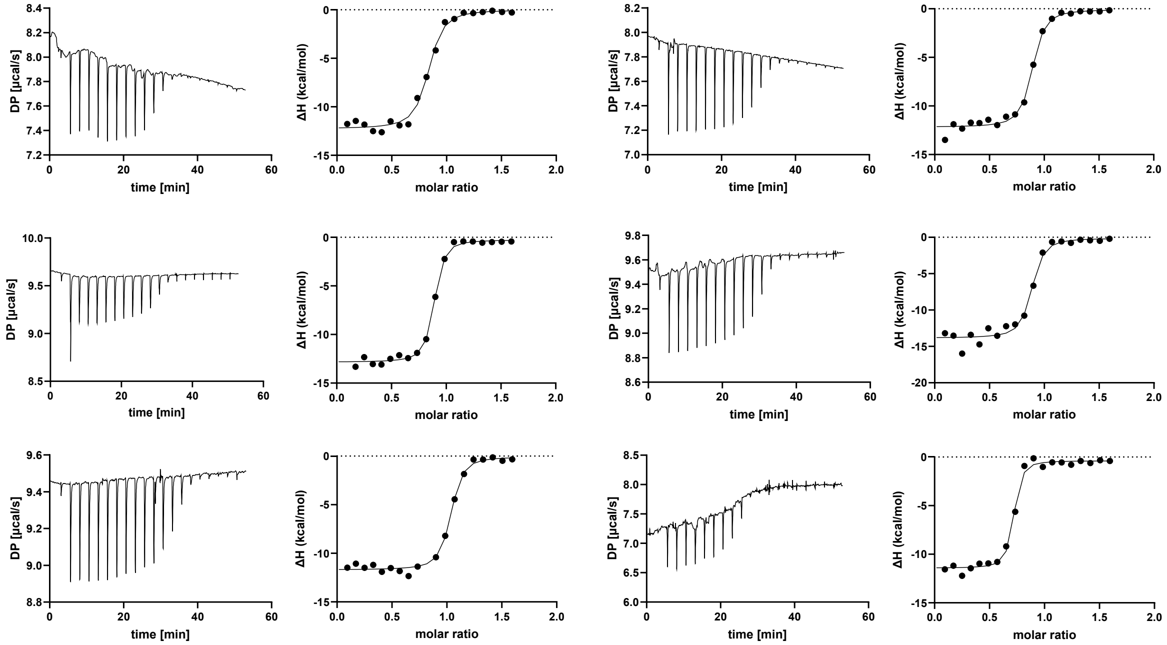


#### (f) SAR11_1361 + Difluorofumarate

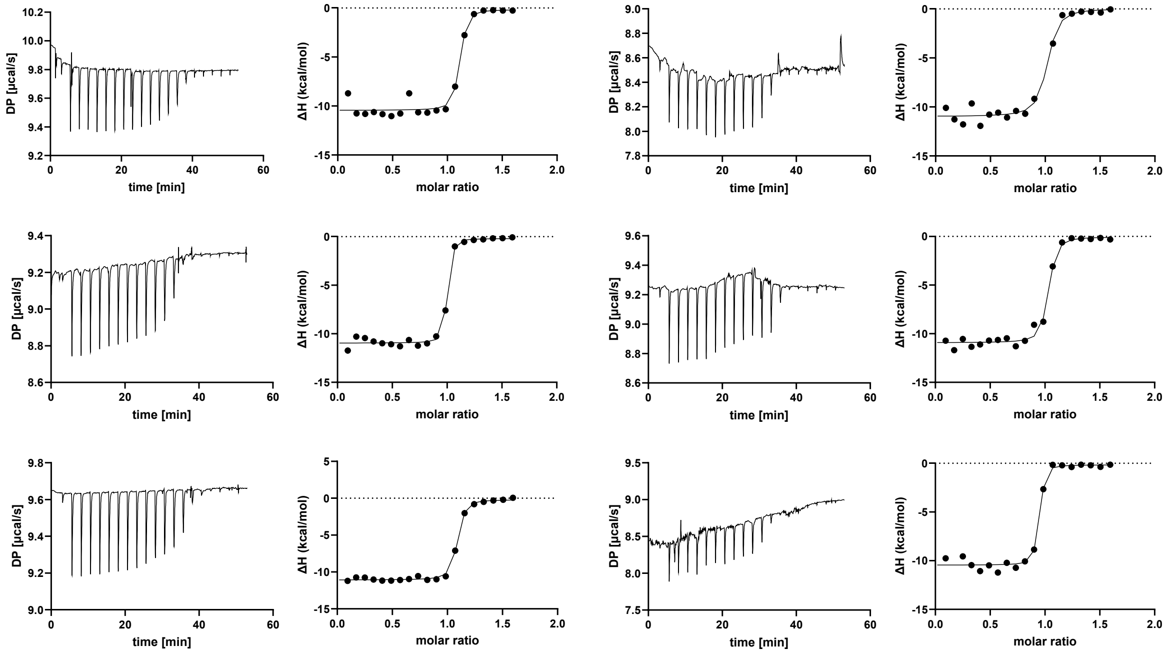

#### (g) SAR11_1361 + Glutarate

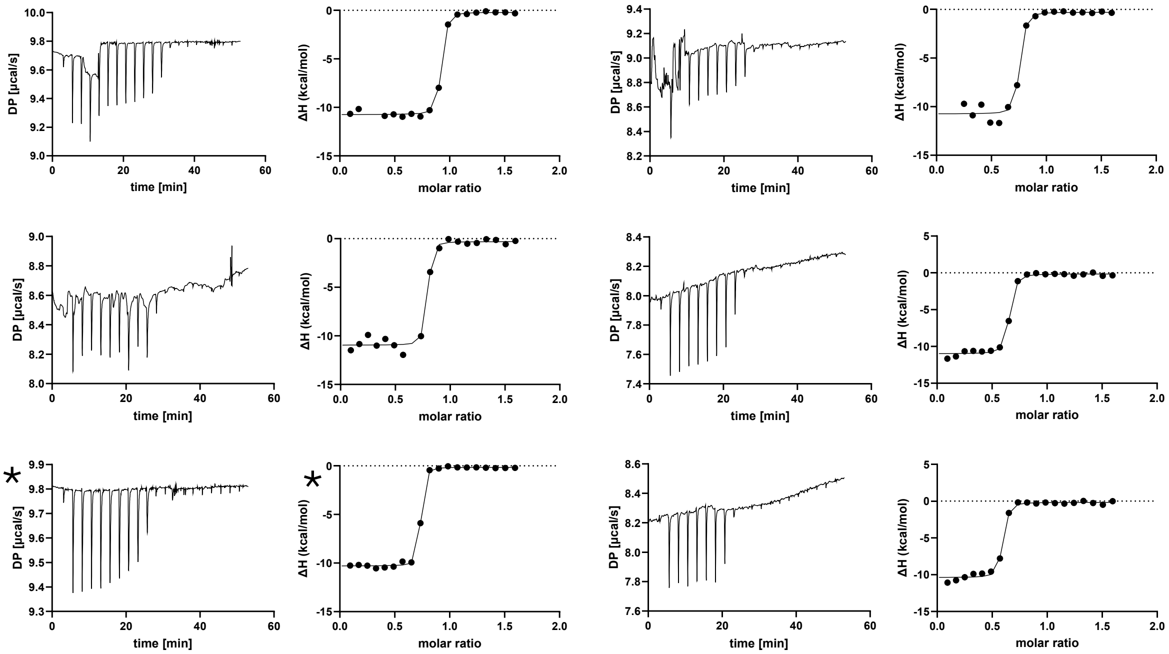


#### (h) SAR11_1361 + 2,2-Difluoroglutarate

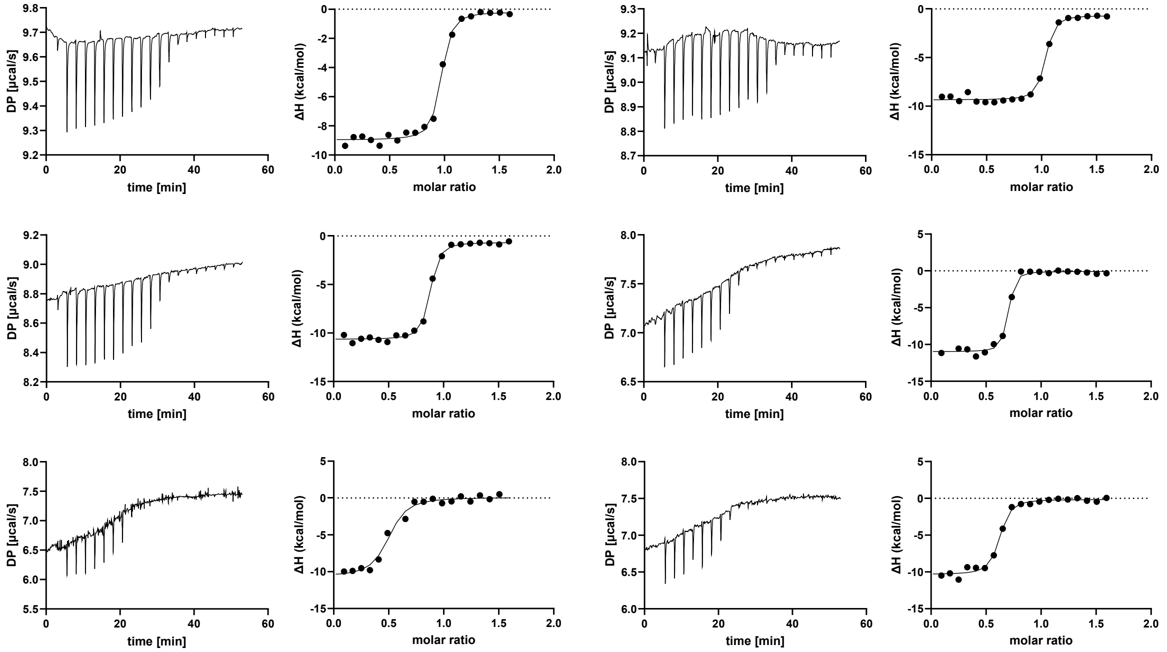

#### (i) SAR11_1361 + Hexafluoroglutarate

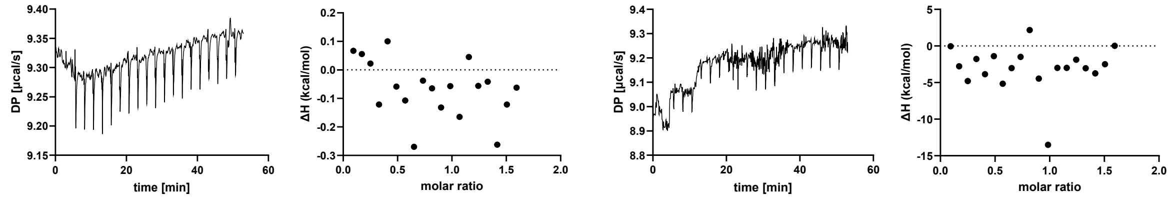


**Figure S3. Determination of binding parameters of SAR11_1361 with ITC.** Protein samples were denatured, refolded, purified, concentrated to 32 µM, and titrated with freshly prepared ligands at a concentration of 280 µM. $K_d$ values were measuring using ITC with the following ligands: (a) succinate, (b) 2,2-difluorosuccinate, (c) tetrafluorosuccinate, (d) fumarate, (e) fluorofumarate, (f) difluorofumarate, (g) glutarate, (h) 2,2-difluoroglutarate, and (i) hexafluoroglutarate. Shown is the raw and integrated data for at least three biological replicates and technical duplicates, with each row constituting a biological replicate. Asterisks mark the measurements which were excluded from calculations of $K_d$ due to being above the limit of quantitation of ITC.

#### (a) Sdel_0447 + Succinate

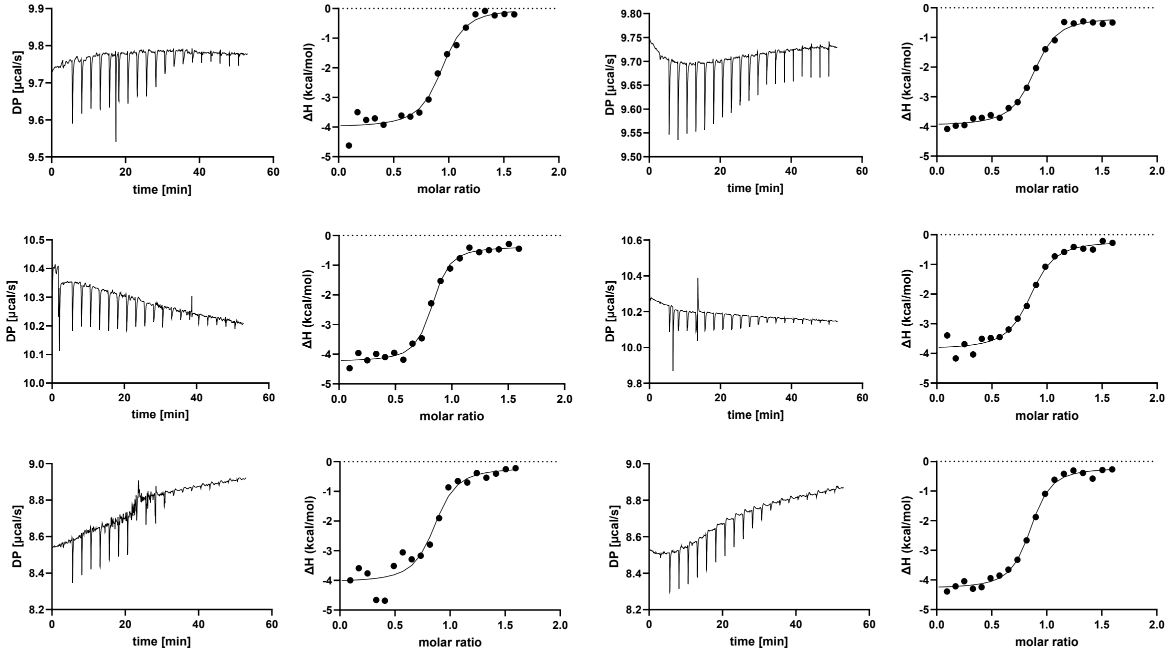


#### (b) Sdel_0447 + 2,2-Difluorosuccinate

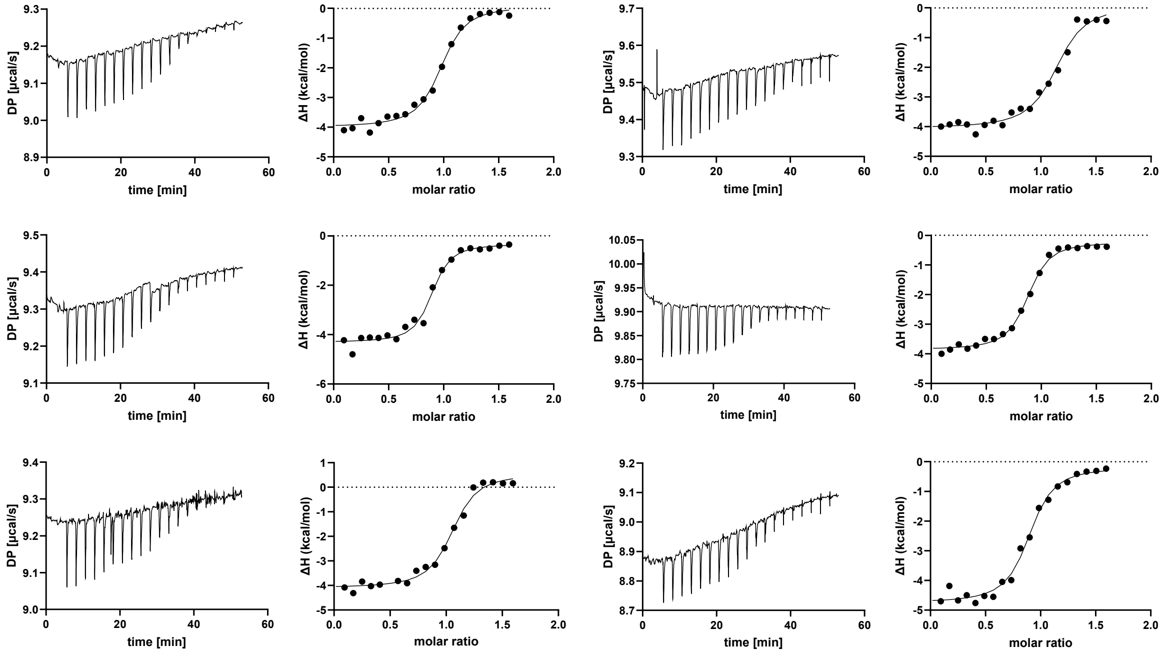

#### (c) Sdel_0447 + Tetrafluorosuccinate

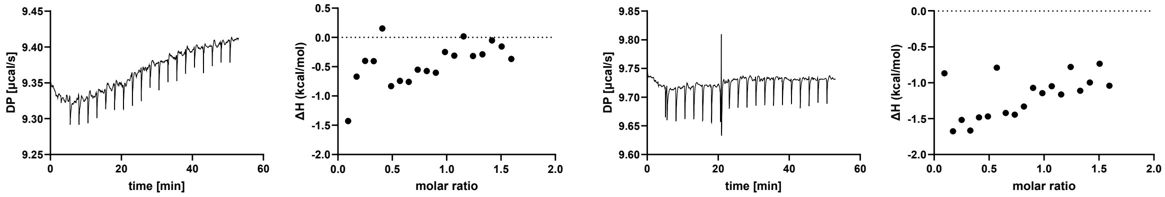


#### (d) Sdel_0447 + Fumarate

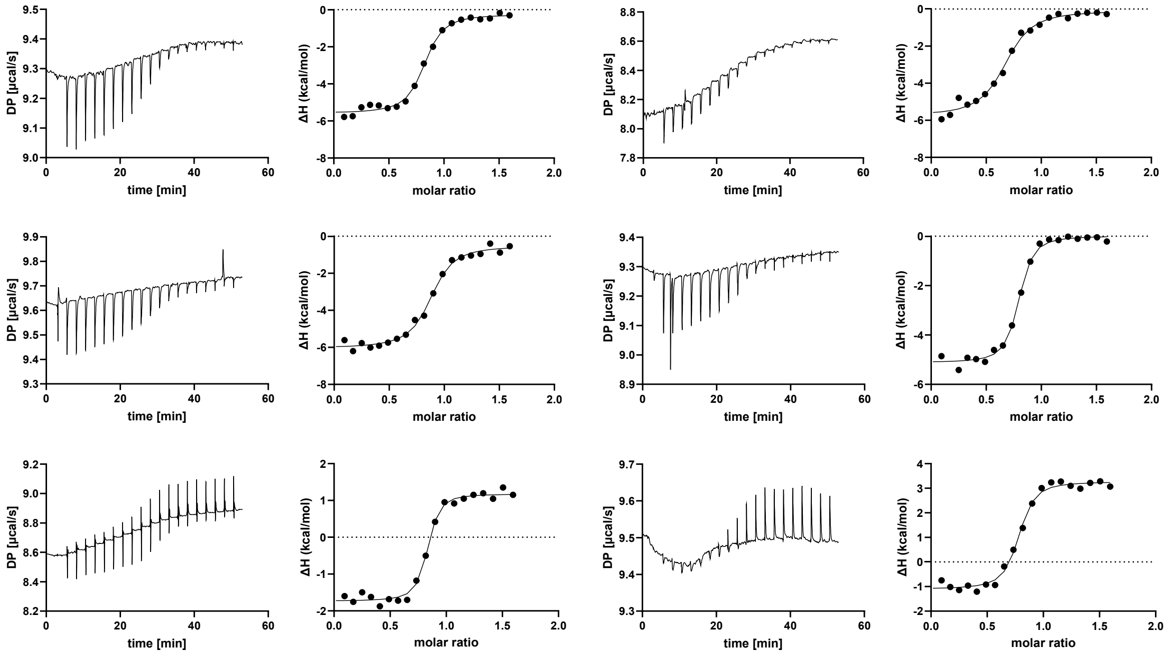


#### (e) Sdel_0447 + Fluorofumarate

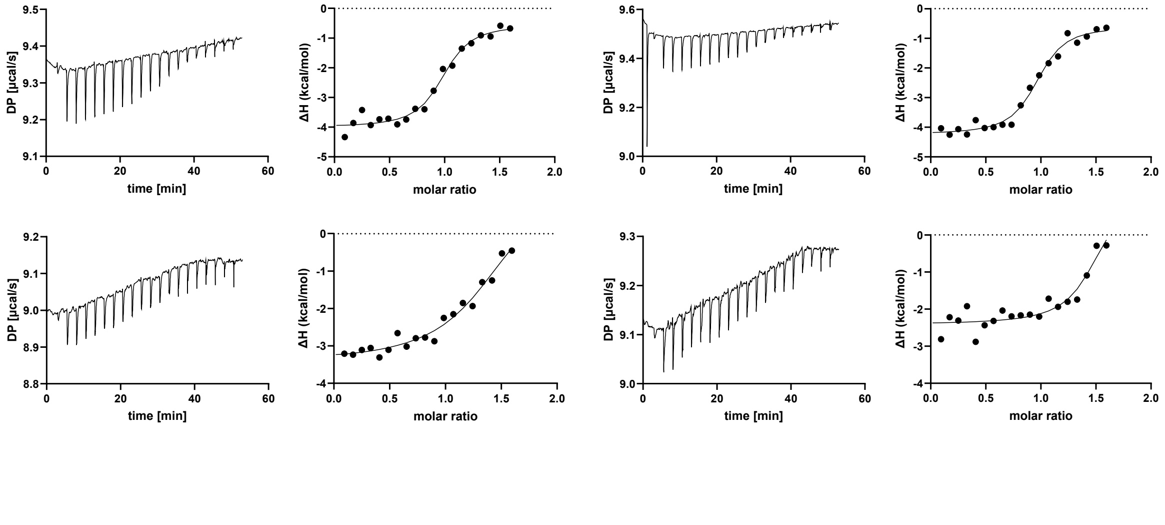

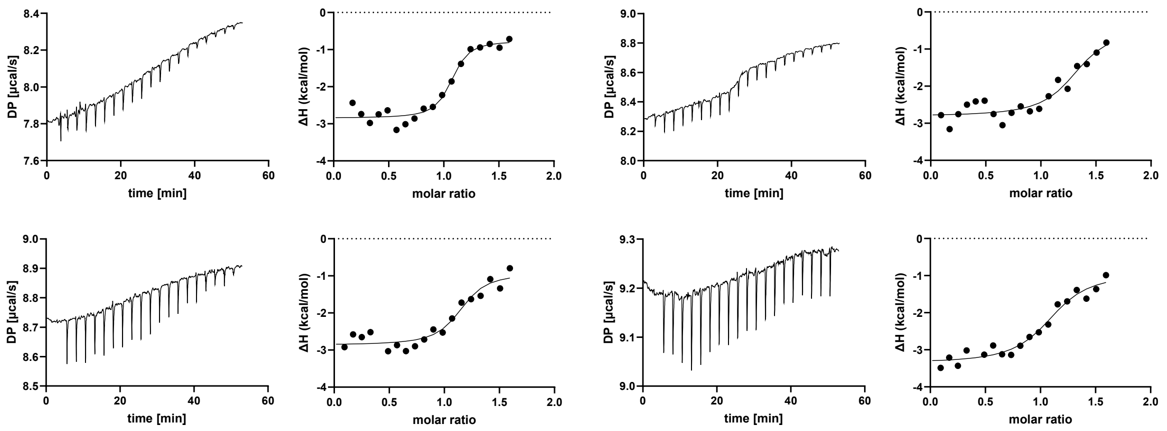


### (f) Sdel_0447 + Difluorofumarate

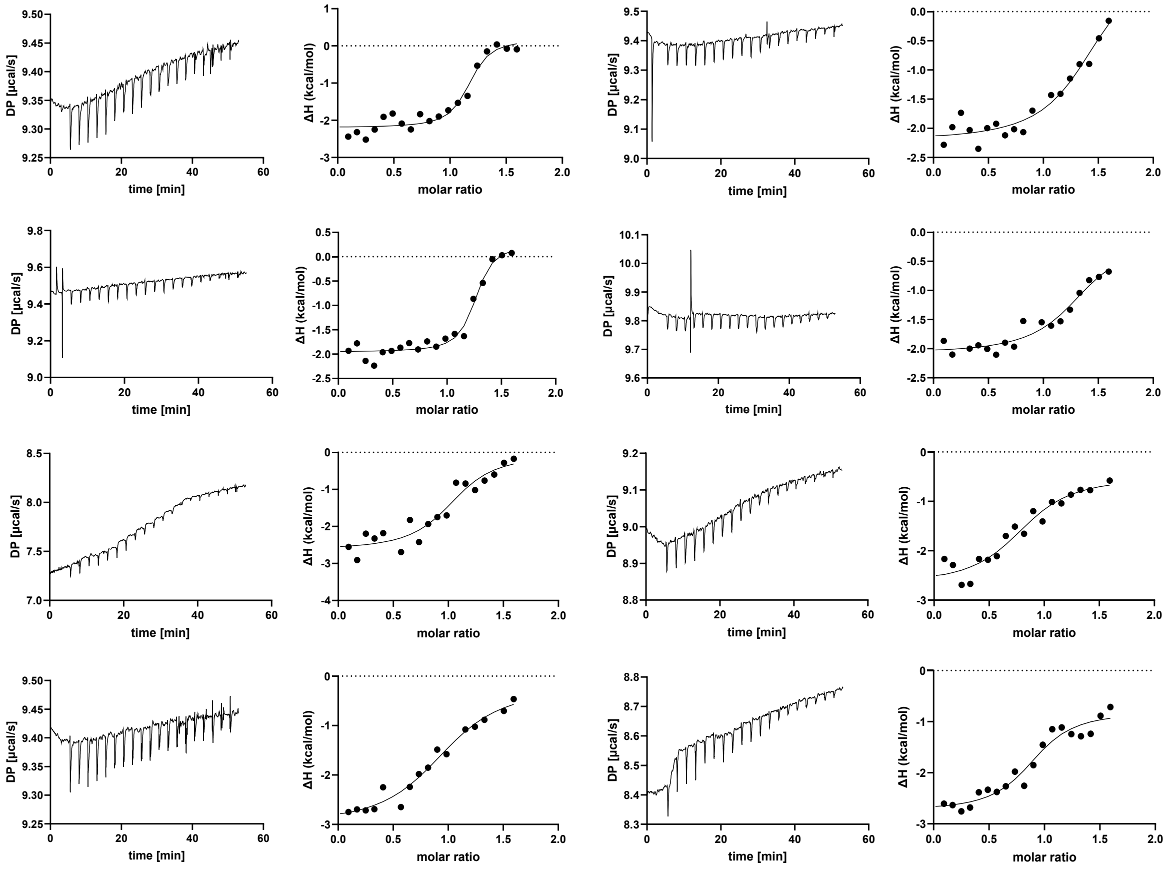


### (g) Sdel_0447 + Glutarate

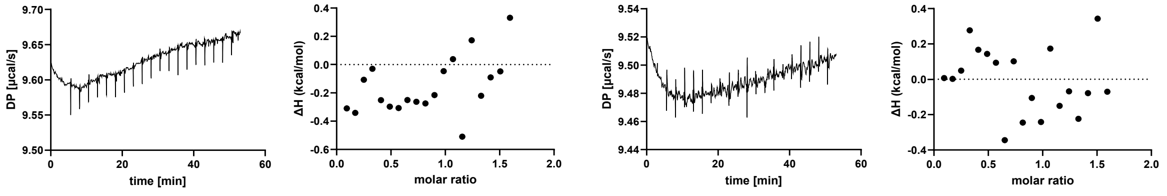

**Figure S4. Determination of binding parameters of Sdel_0447 with ITC.** Protein samples were denatured, refolded, purified, concentrated to 32 μM, and titrated with freshly prepared ligands at a concentration of 280 μM. $K_d$ values were measuring using ITC with the following ligands: (a) succinate, (b) 2,2-difluorosuccinate, (c) tetrafluorosuccinate, (d) fumarate, (e) fluorofumarate, (f) difluorofumarate, (g) glutarate. Shown is the raw and integrated data for at least three biological replicates and technical duplicates, with each row constituting a biological replicate.

#### (a) B5M07_04440 Tyr42Ala + Succinate

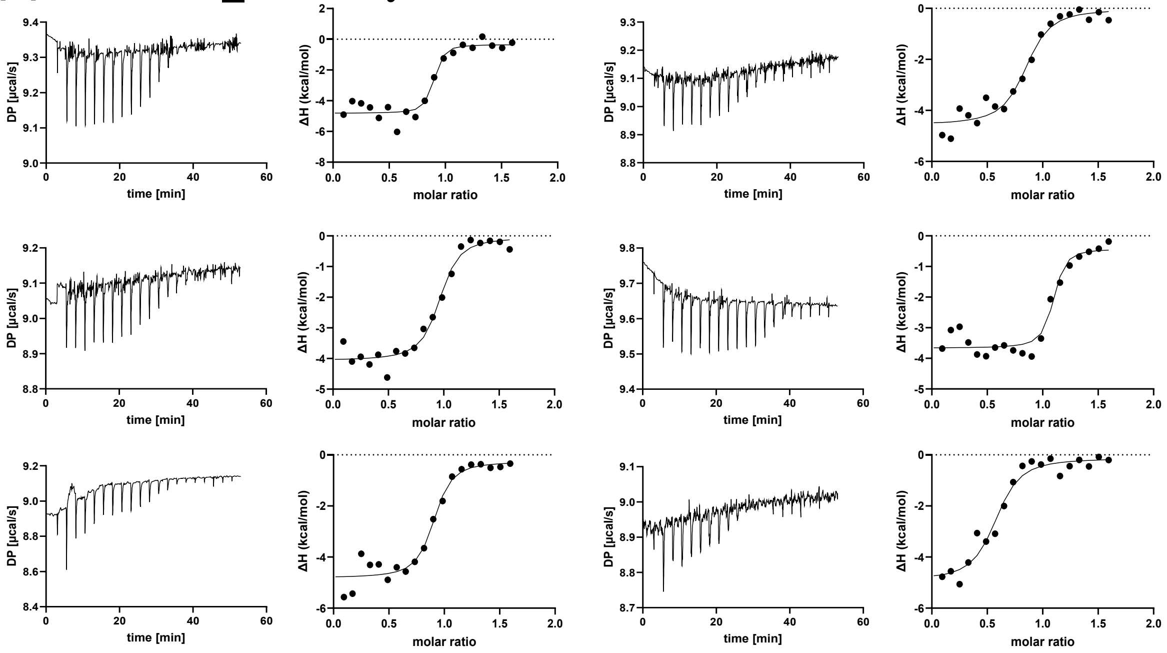


#### (b) B5M07_04440 Tyr42Ala + 2,2-Difluorosuccinate

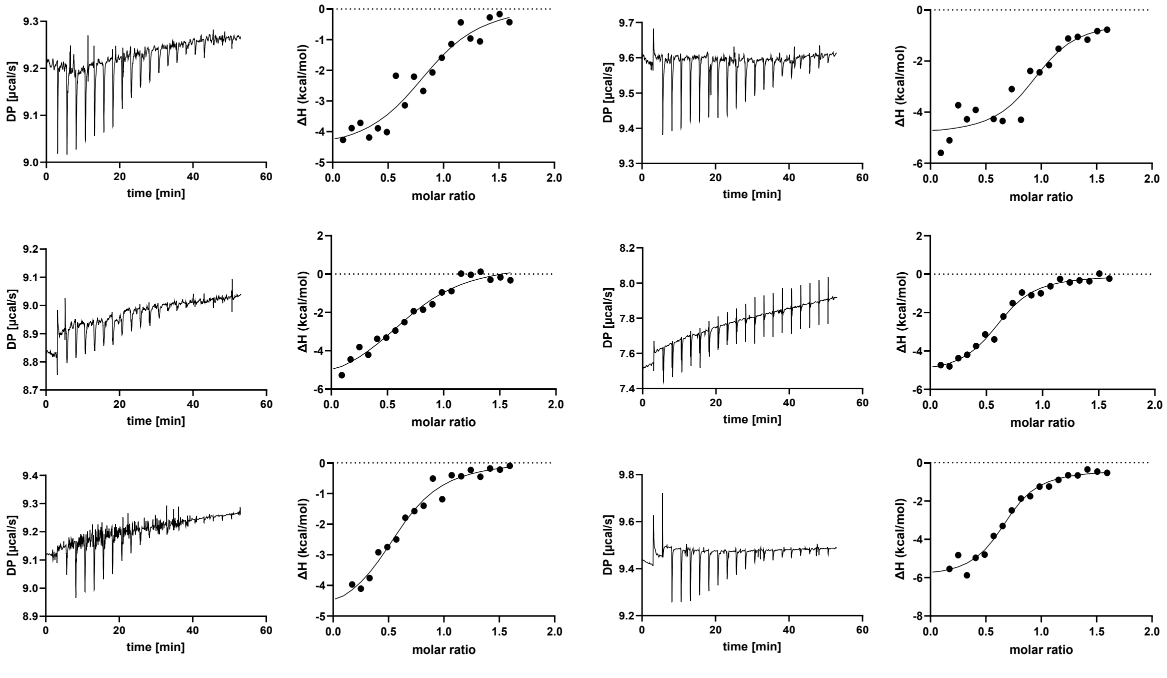

#### (c) B5M07_04440 Tyr42Ala + Tetrafluorosuccinate

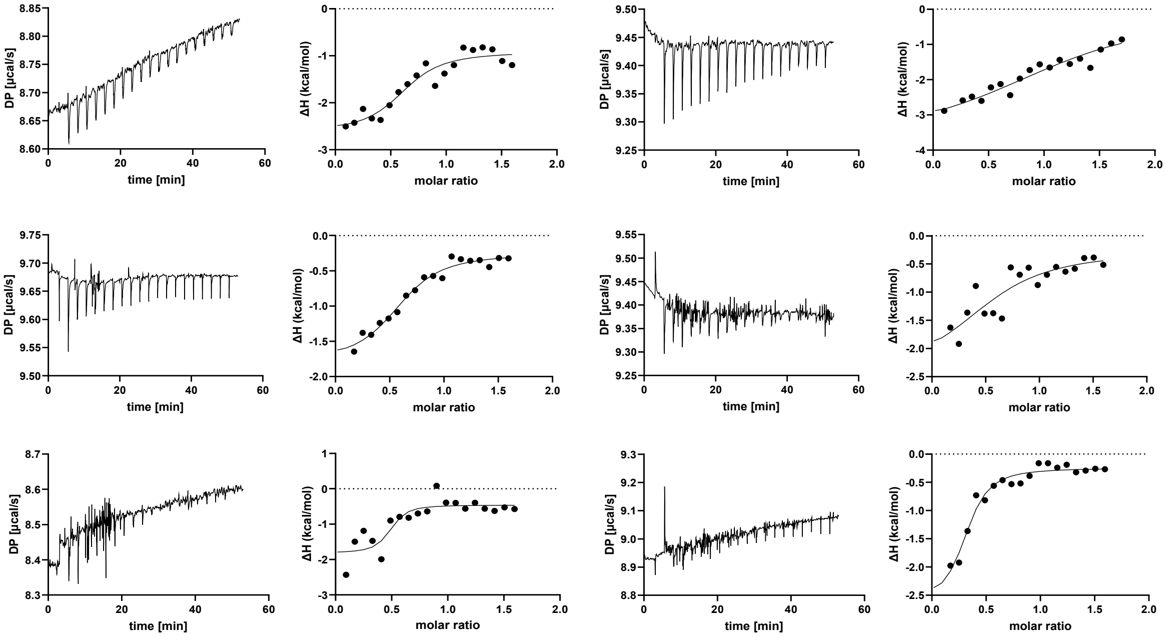


#### (d) B5M07_04440 Tyr42Ala + Fumarate

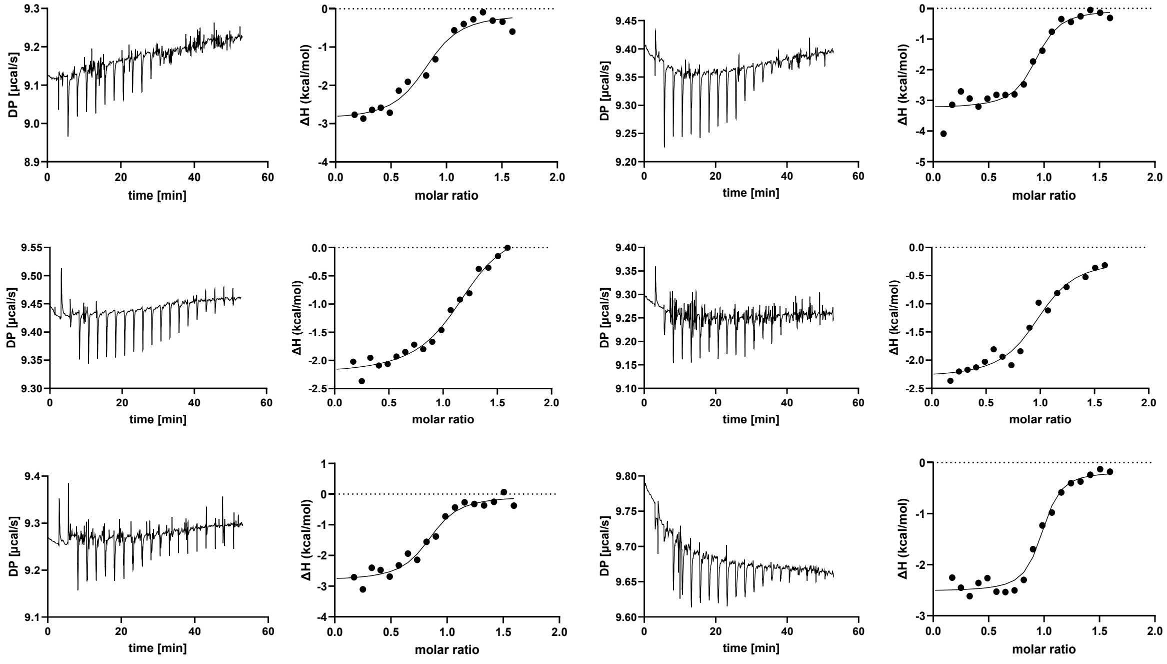

### (e) B5M07_04440 Tyr42Ala + Fluorofumarate

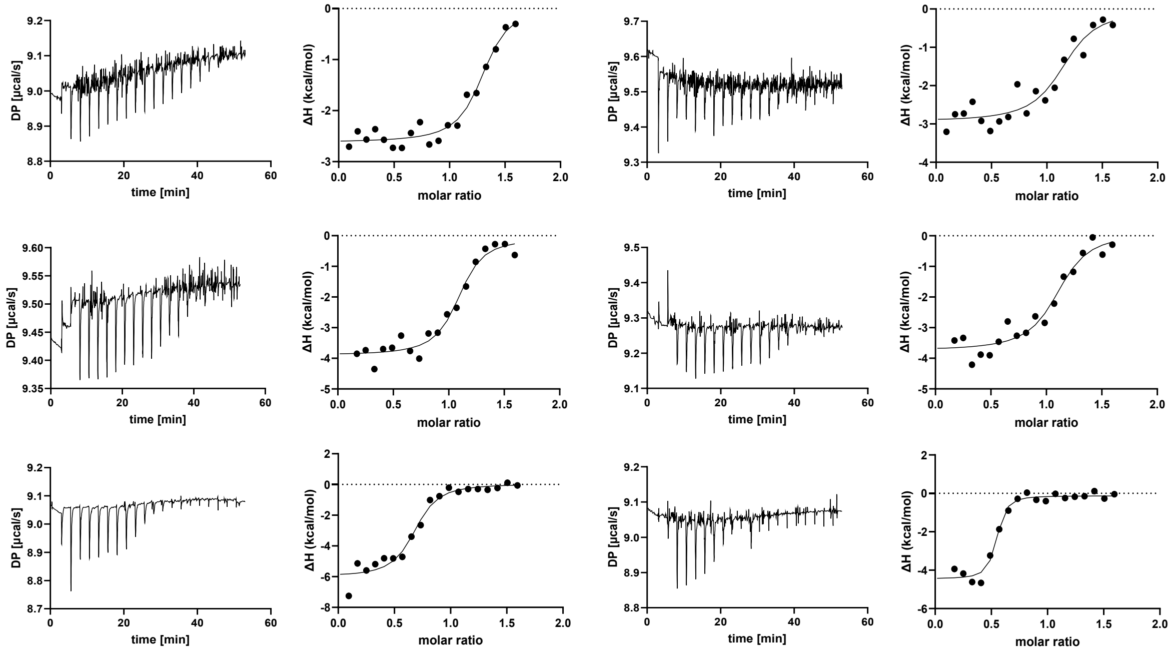


### (f) B5M07_04440 Tyr42Ala + Difluorofumarate

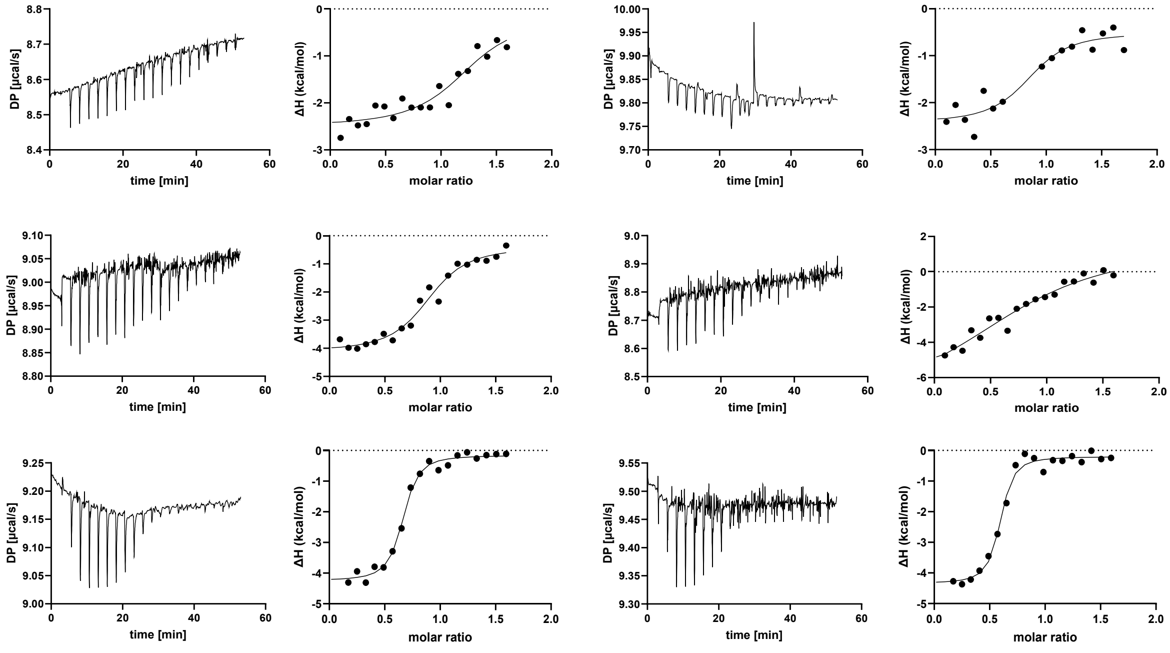

**Figure S5. Determination of binding parameters of B5M07_04440 Tyr42Ala with ITC.** Protein samples were denatured, refolded, purified, concentrated to 32 μM, and titrated with freshly prepared ligands at a concentration of 280 μM. $K_d$ values were measuring using ITC with the following ligands: (a) succinate, (b) 2,2-difluorosuccinate, (c) tetrafluorosuccinate, (d) fumarate, (d) fluorofumarate, and (e) difluorofumarate. Shown is the raw and integrated data for at least three biological replicates and technical duplicates, with each row constituting a biological replicate.

#### (a) SAR11_1361 Phe187Ala + Succinate

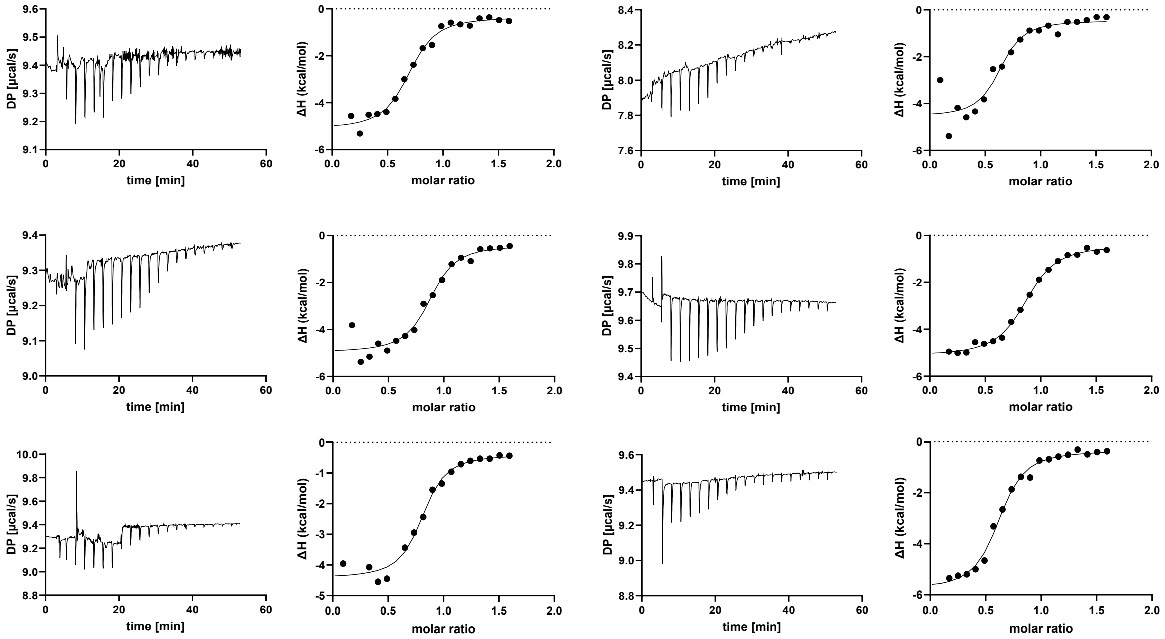


#### (b) SAR11_1361 Phe187Ala + 2,2-Difluorosuccinate

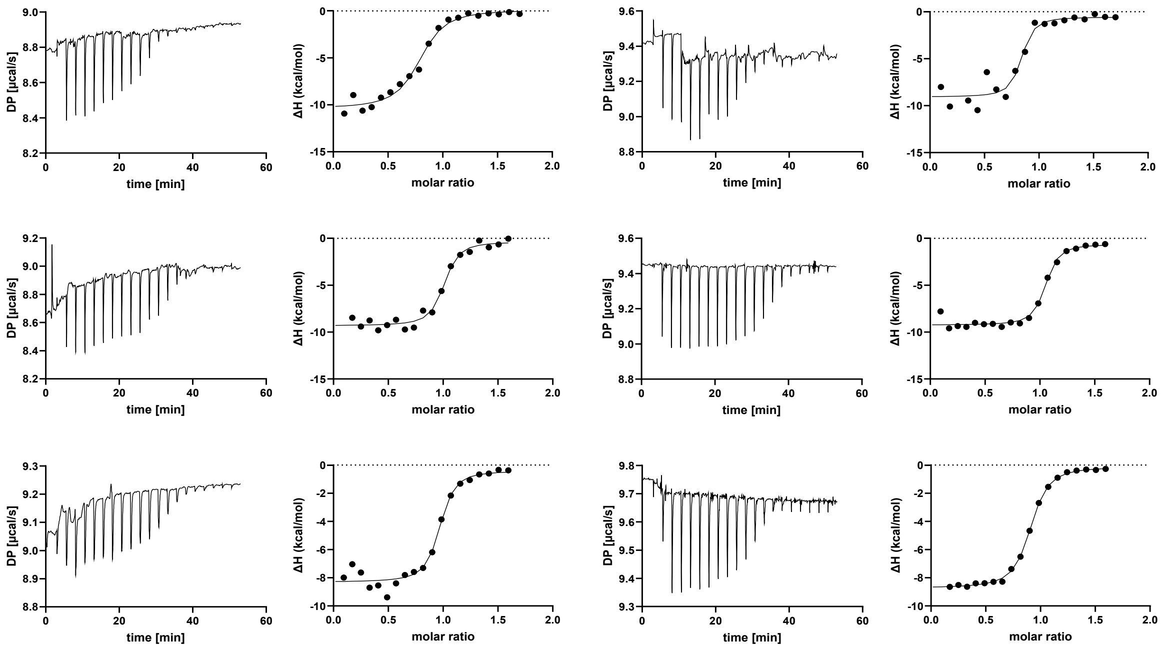

#### (c) SAR11_1361 Phe187Ala + Tetrafluorosuccinate

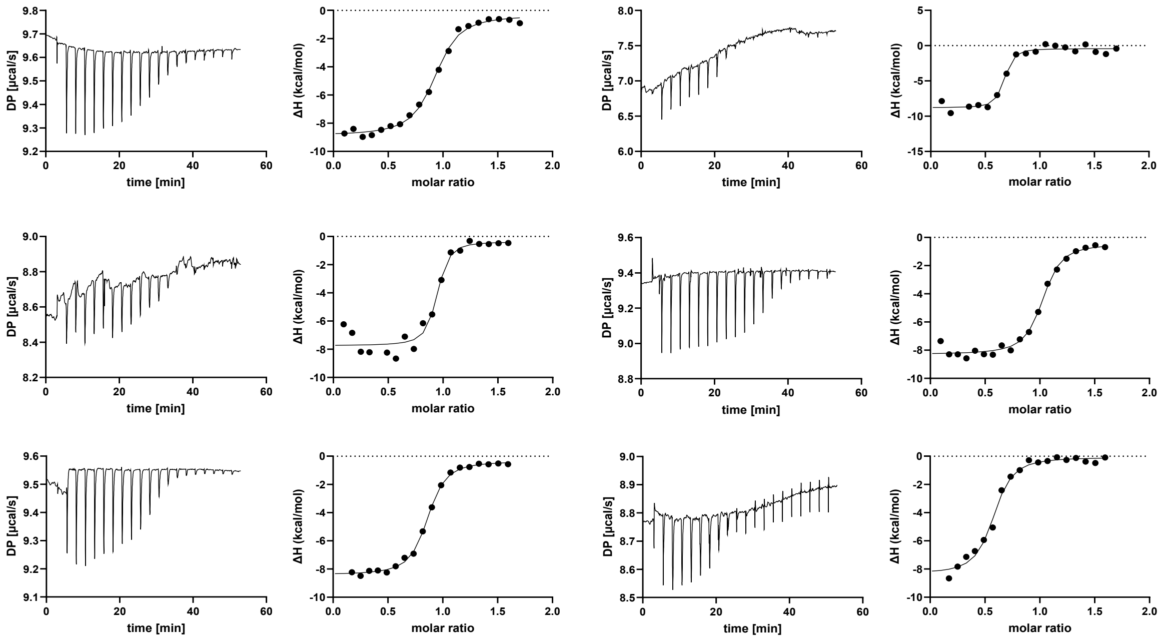


#### (d) SAR11_1361 Phe187Ala + Fumarate

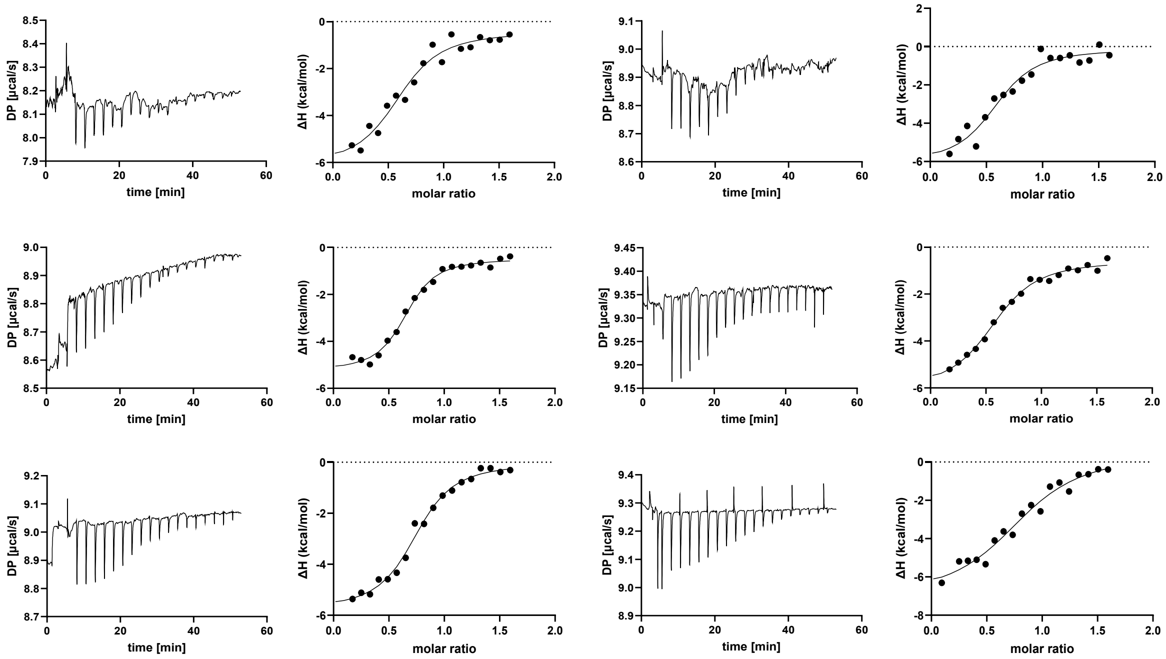

### (e) SAR11_1361 Phe187Ala + Fluorofumarate

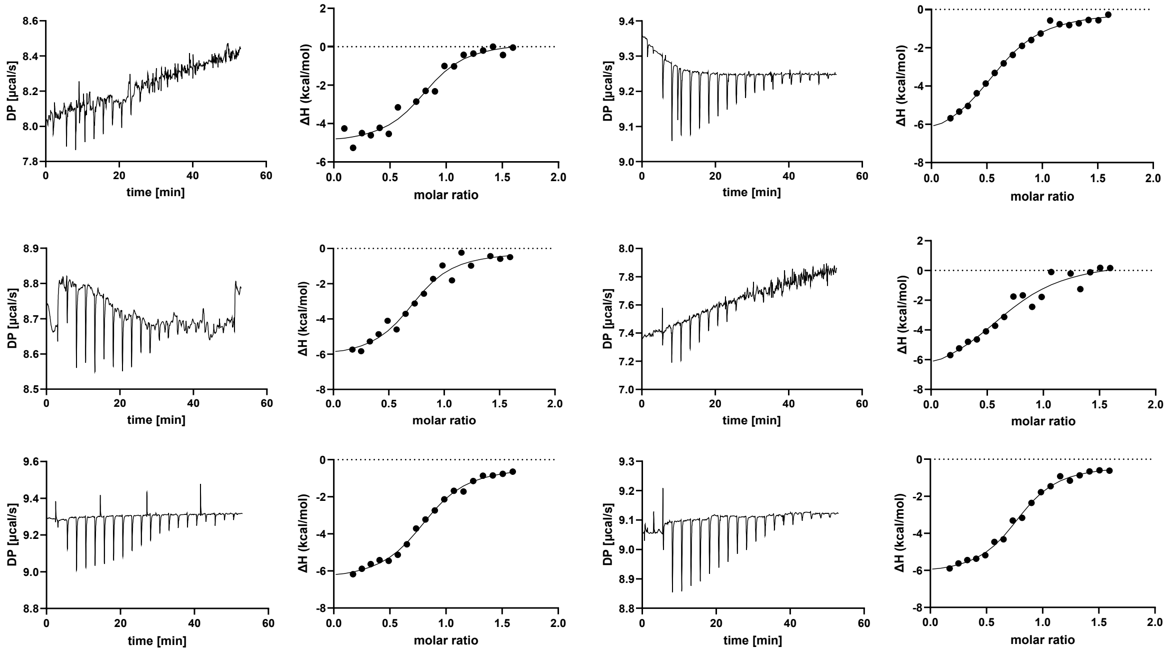


### (f) SAR11_1361 Phe187Ala + Difluorofumarate

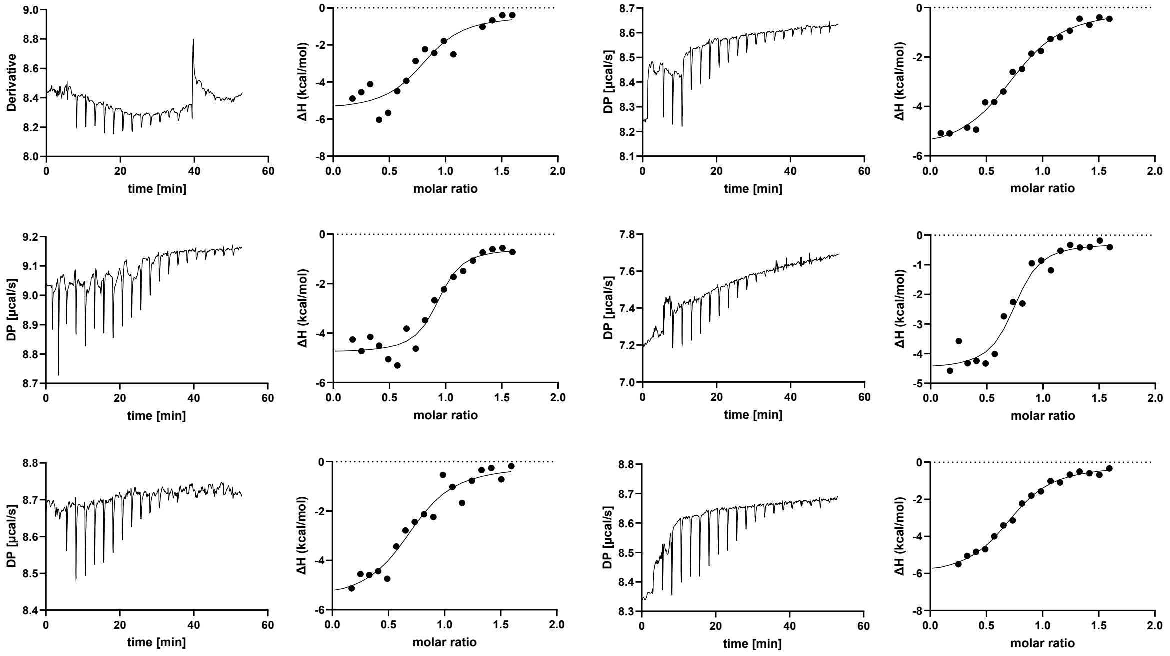

#### (g) SAR11_1361 Phe187Ala + Glutarate

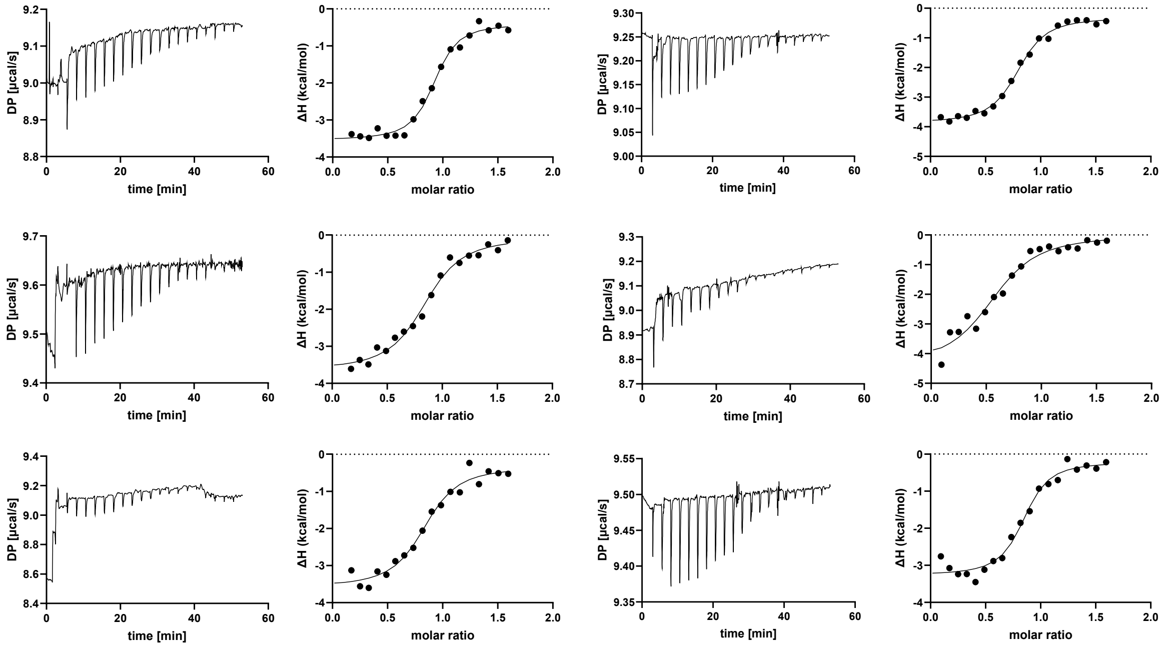


#### (h) SAR11_1361 Phe187Ala + 2,2-Difluoroglutarate

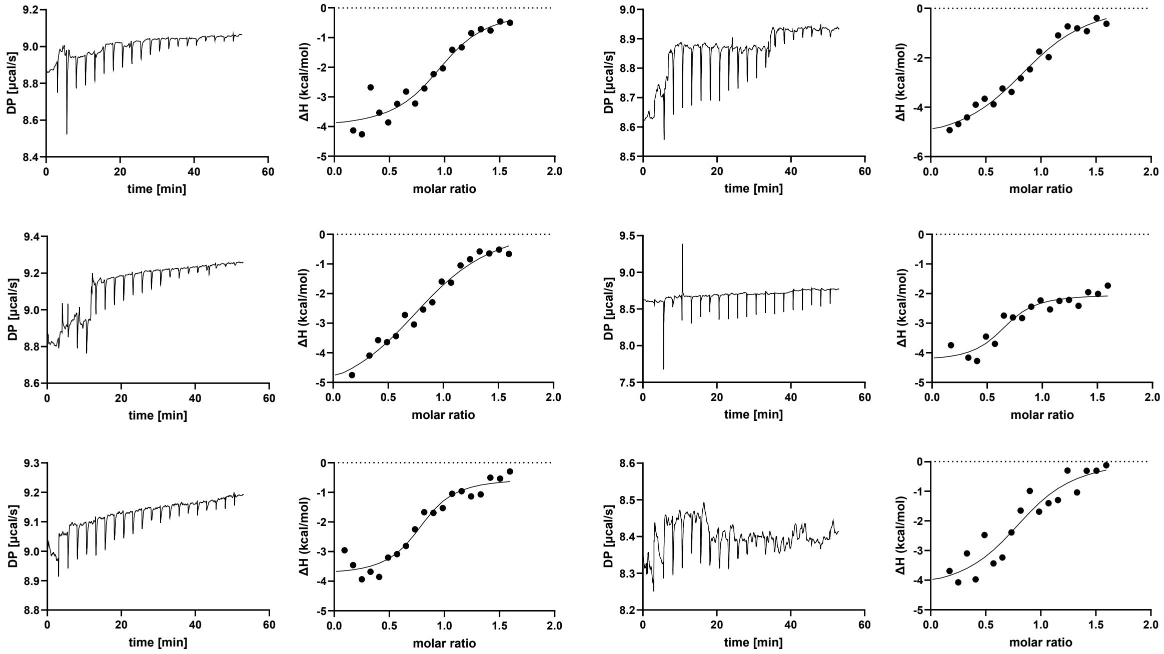

#### (i) SAR11_1361 Phe187Ala + Hexafluoroglutarate

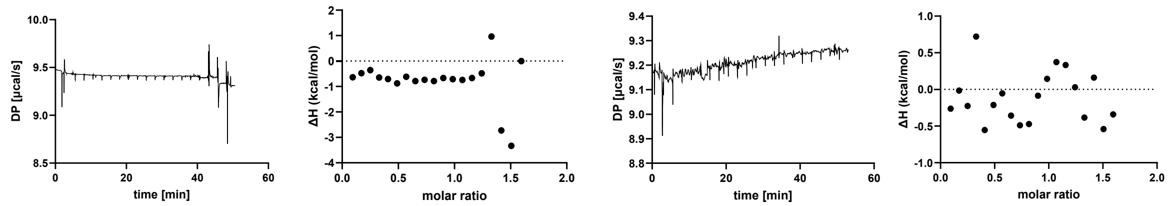


**Figure S6. Determination of binding parameters of SAR11_1361 Phe187Ala with ITC.** Protein samples were denatured, refolded, purified, concentrated to 32 µM, and titrated with freshly prepared ligands at a concentration of 280 µM. $K_d$ values were measuring using ITC with the following ligands: (a) succinate, (b) 2,2-difluorosuccinate, (c) tetrafluorosuccinate, (d) fumarate, (e) fluorofumarate, (f) difluorofumarate, (g) glutarate, (h) 2,2-difluoroglutarate, and (i) hexafluoroglutarate. Shown is the raw and integrated data for at least three biological replicates and technical duplicates, with each row constituting a biological replicate.

#### (a) Succinate

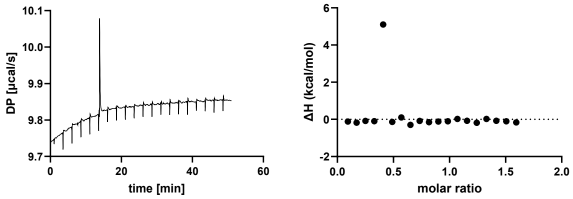


#### (b) 2,2-Difluorosuccinate

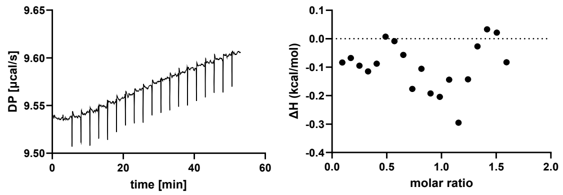


#### (c) Tetrafluorosuccinate

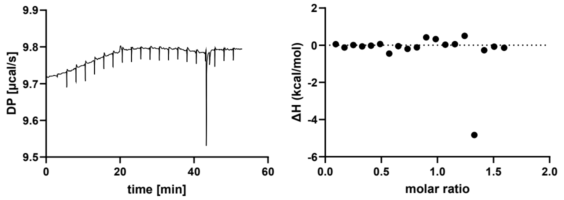


#### (d) Fumarate

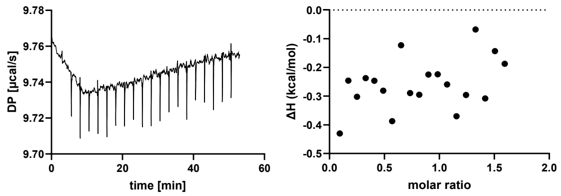


#### (e) Fluorofumarate

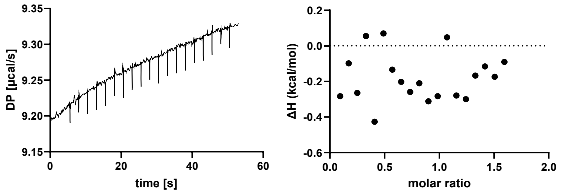


#### (f) Difluorofumarate

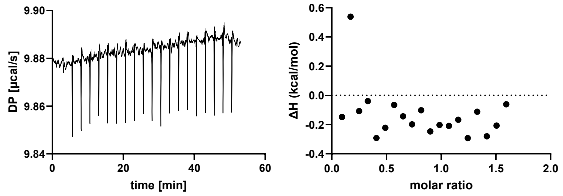


#### (a) Glutarate

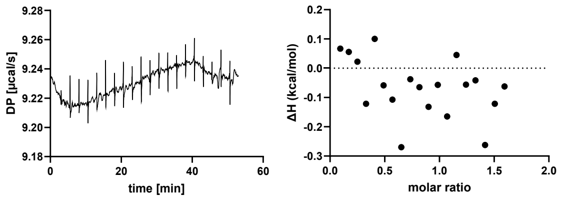


#### (b) 2,2-Difluoroglutarate

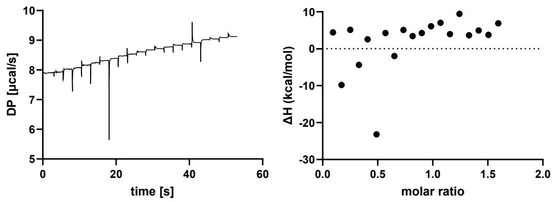


#### (a) Hexafluoroglutarate

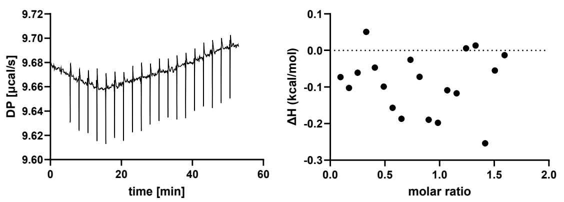

**Figure S7. Control ITC experiments.** ITC samples without protein were measured for the following ligands: (a) succinate, (b) 2,2-difluorosuccinate, (c) tetrafluorosuccinate, (d) fumarate, (e) fluorofumarate, (f) difluorofumarate, (g) glutarate, (h) 2,2-difluoroglutarate, and (i) hexafluoroglutarate.

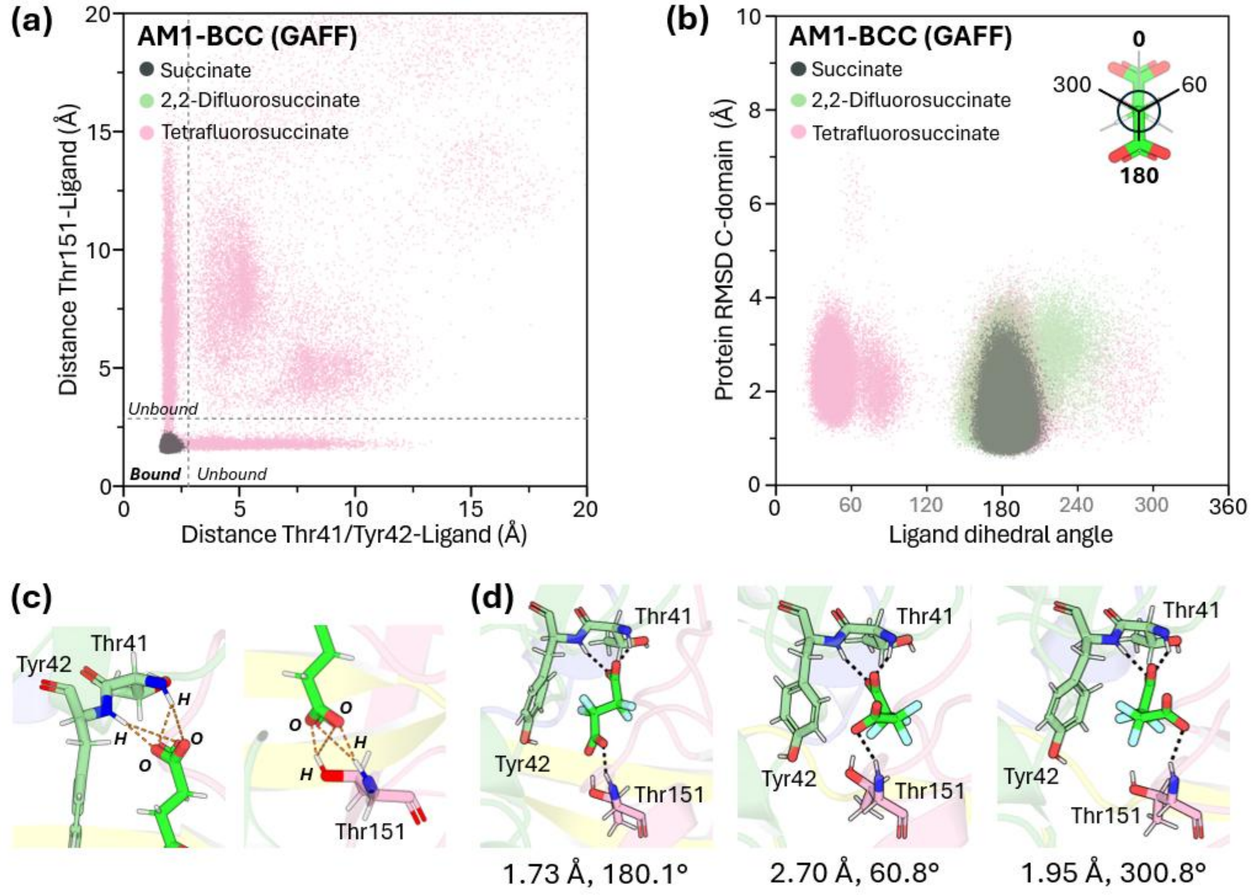


**Figure S8. Results from MD simulations of B5M07_04440 using AM1-BCC charges and the GAFF force field.** (a) Distribution of conformations sampled during the MD simulations in the two-dimensional space defined by the protein–ligand distances. The x-axis represents the minimum distance between the ligand carboxylate oxygen atoms and the backbone amide hydrogen atoms of Thr41/Tyr42, while the y-axis represents the corresponding minimum distance to the backbone amide hydrogen atom of Thr151. The grey dashed lines indicate the 2.6 Å distance cutoff used to define the bound state. (b) Distribution of bound-state conformations in the two-dimensional space defined by the C-domain RMSD and ligand dihedral angle. The C-domain RMSD was calculated after superimposing the backbone atoms (C, CA, N, and O) of the N-domain onto the initial closed-state structure (Figure S13). (c) Definition of the protein–ligand distances used to characterize the bound state. Orange dashed lines indicate the atom pairs considered for each distance calculation, with the minimum distance used for each simulation frame (Figure S9). (d) Representative snapshots of TFSA conformations with different dihedral angles sampled during the protein–tetrafluorosuccinate complex simulation. Polar contacts between the protein and ligand are shown as black dashed lines. The tetrafluorosuccinate dihedral angle and corresponding C-domain RMSD (Å) are indicated below each snapshot. Polar contacts were identified using PyMOL with the default settings (3.2 Å cutoff).

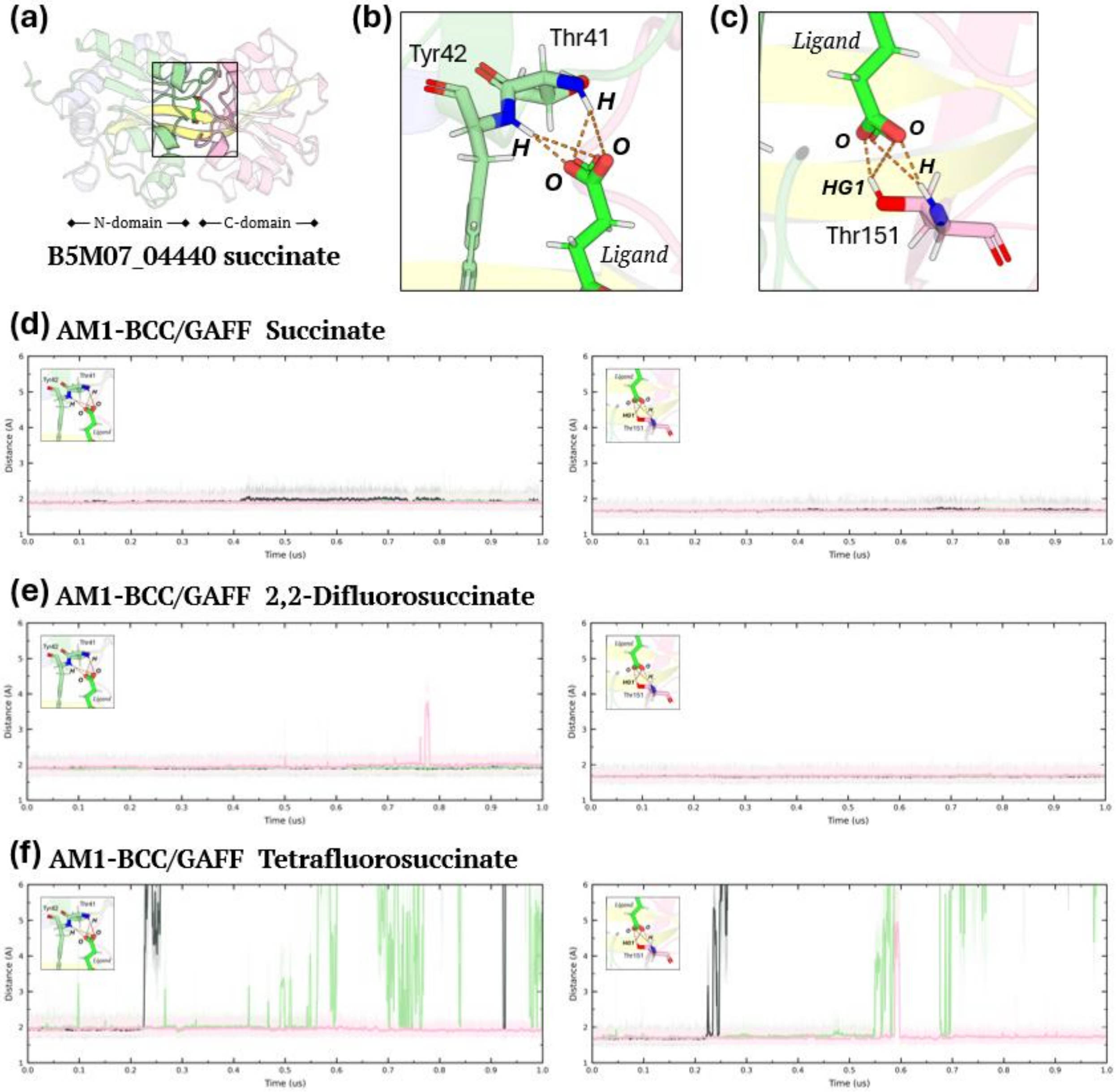


**Figure S9. Protein-ligand distances used to characterize the bound state in simulation of B5M07_04440 using AM1-BCC charges and the GAFF force field.** (a) Representative structure of the B5M07_04440-ligand complex. (b,c) Definition of the protein–ligand distances used to characterize the bound state. The minimum distances were calculated between the ligand carboxylate oxygen atoms and (b) the backbone amide hydrogen atoms of Thr41 and Tyr42, and (c) the backbone amide hydrogen and sidechain HG1 atom of Thr151. A distance cutoff of 2.6 Å was used to define the bound state. (d-f) Time series of the protein–ligand distances for three independent 1-μs MD simulations of (d) succinate, (e) 2,2-difluorosuccinate, and (f)

tetrafluorosuccinate. Replicates 1, 2, and 3 are shown in grey, green, and pink, respectively. The y-axis range was kept constant across all simulation systems to facilitate comparison.

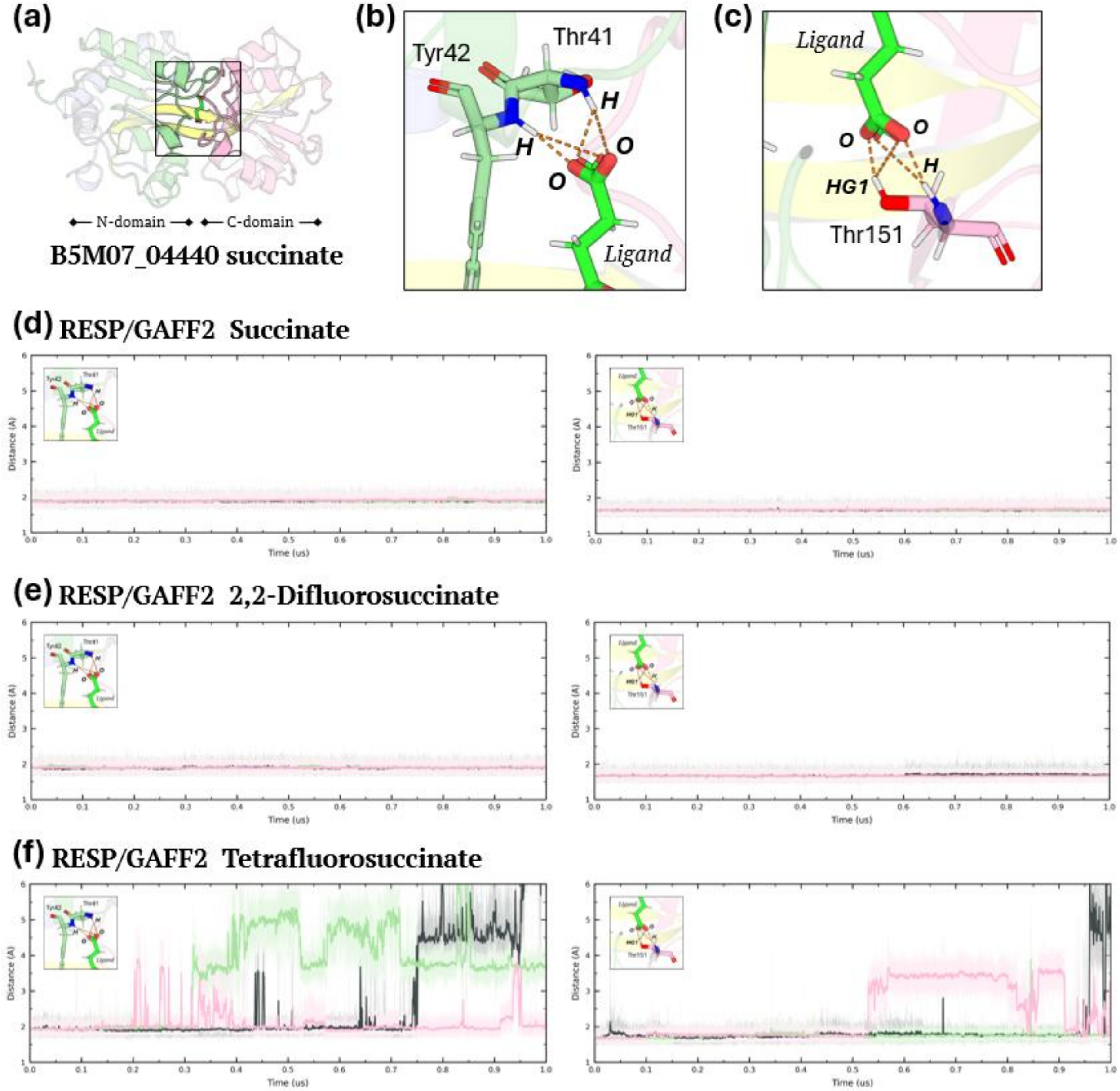


**Figure S10. Protein–ligand distances used to characterize the bound state of B5M07_04440 in simulations using RESP charges and the GAFF2 force field.** (a) Representative structure of the B5M07_04440-ligand complex. (b,c) Definition of the protein–ligand distances used to characterize the bound state. The minimum distances were calculated between the ligand carboxylate oxygen atoms and (b) the backbone amide hydrogen atoms of Thr41 and Tyr42 and (c) the backbone amide hydrogen and sidechain HG1 atom of Thr151. A distance cutoff of 2.6 Å was used to define the bound state. (d-f) Time series of the protein–ligand distances for three independent 1-μs MD simulations of (d) succinate, (e) 2,2-difluorosuccinate, and (f)

tetrafluorosuccinate. Replicates 1, 2, and 3 are shown in grey, green, and pink, respectively. The y-axis range was kept constant across all simulation systems to facilitate comparison.

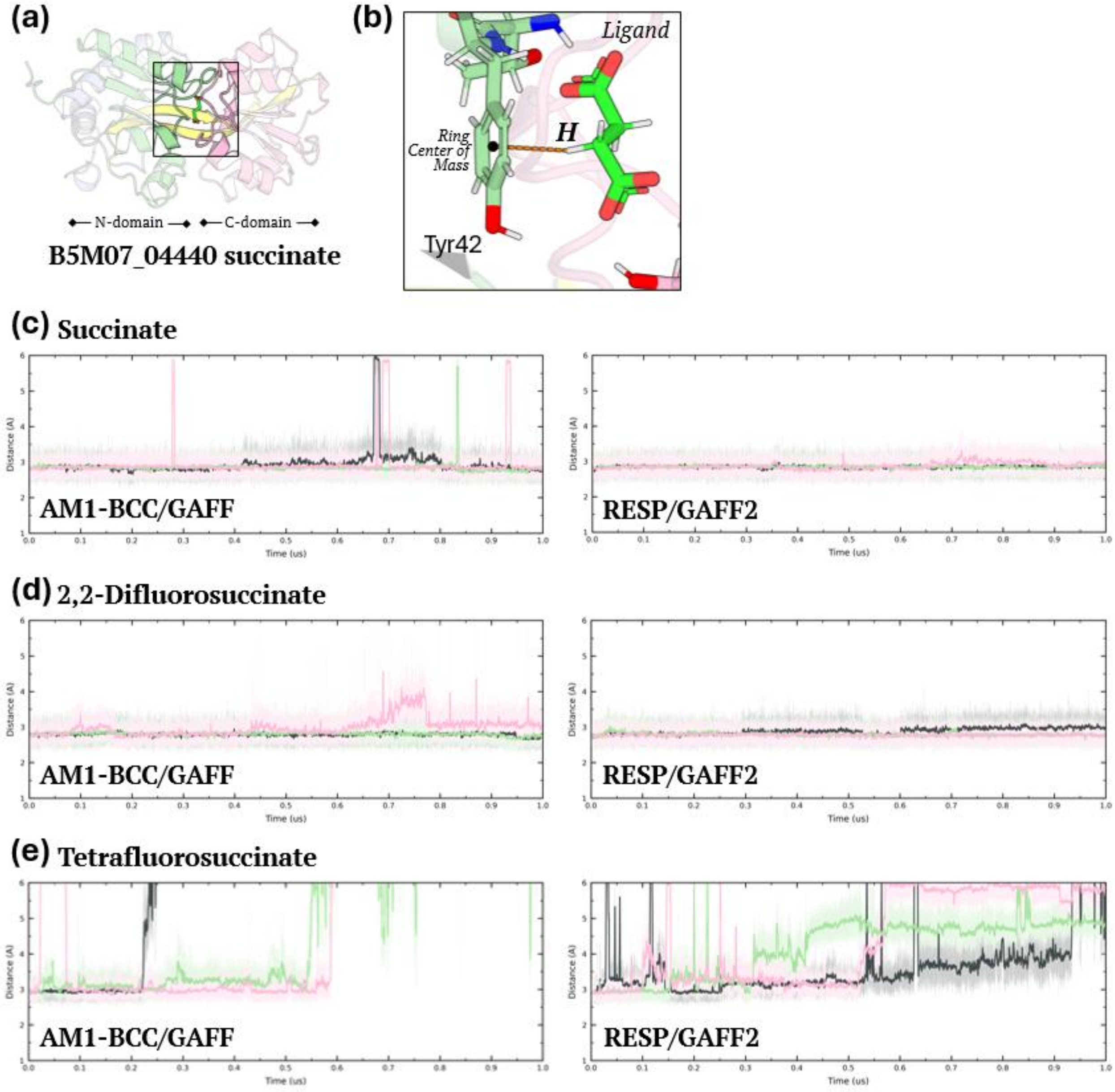


**Figure S11. Protein–ligand distances between the ligand H/F atoms and the aromatic ring of Tyr42 in MD simulations of B5M07_04440.** (a) Representative structure of the B5M07_04440–ligand complex. (b) Definition of the distance between the centre of mass of the Tyr42 aromatic ring and the ring-facing H or F atom of the ligand. (c-e) Time series of the Tyr42 aromatic-ring–ligand distances for three independent 1-μs MD simulations of (c) succinate, (d) 2,2-difluorosuccinate, and (e) tetrafluorosuccinate using AM1-BCC/GAFF (left) and RESP/GAFF2 (right). Replicates 1, 2, and 3 are shown in grey, green, and pink, respectively. The y-axis range was kept constant across all simulation systems to facilitate comparison.

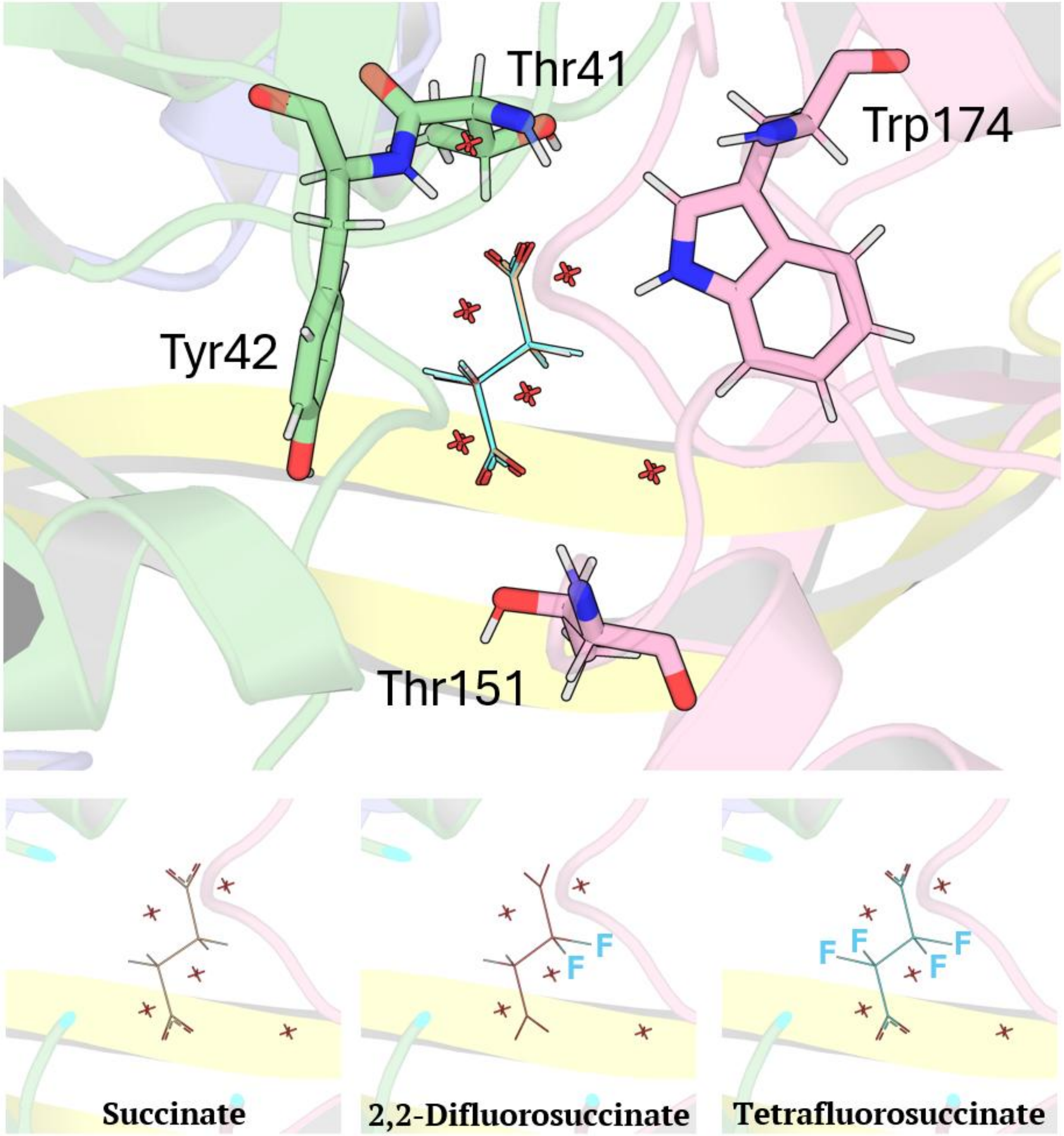


**Figure S12. Ligand poses used to prepare the B5M07_04440-ligand MD simulation systems.** Superimposition of succinate, 2,2-difluorosuccinate, and tetrafluorosuccinate in the B5M07_04440 binding pocket. Succinate, 2,2-difluorosuccinate, and tetrafluorosuccinate are shown in wheat, red, and blue, respectively. The poses for succinate and 2,2-difluorosuccinate were obtained from the crystal structures, whereas the pose for tetrafluorosuccinate was modeled as described in the methods. Key binding-site residues and conserved crystallographic water molecules are shown for reference. Individual ligand poses are shown at the bottom.

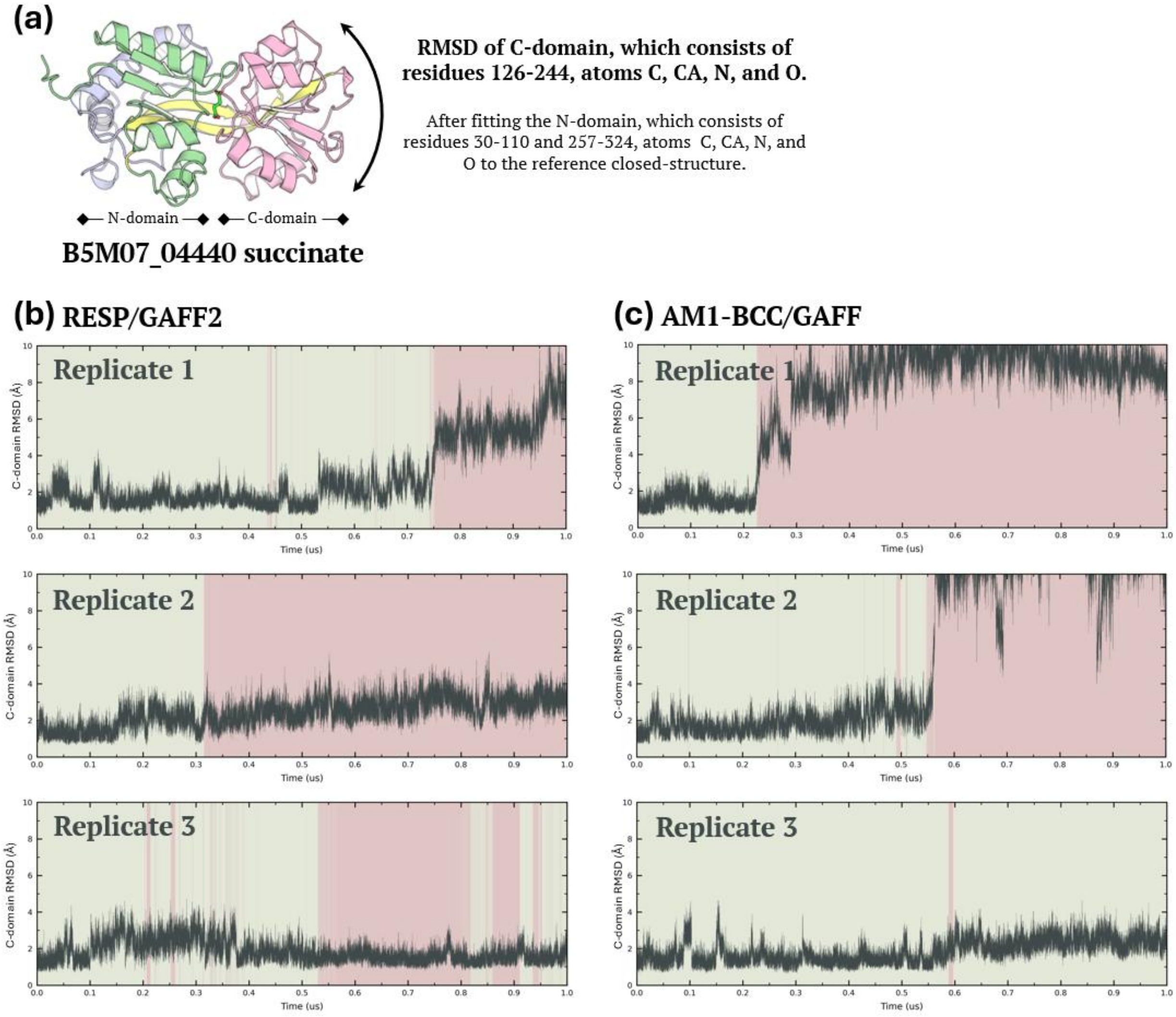


**Figure S13. RMSD of the C-domain of the B5M07_04440 in the tetrafluorosuccinate complex during MD simulations.** (a) Definition of the structural alignment used for the RMSD calculation. The RMSD of the C-domain after superimposing the N-domain to the initial closed-state structure used for the MD simulations. (b) Time series of the C-domain RMSD for three independent 1-µs MD simulations using RESP charges and the GAFF2 force field for ligand parameterization. Green and red shading indicate the ligand-bound and unbound states, respectively, as defined in Figure 3c. (c) Corresponding C-domain RMSD time series for simulations using AM1-BCC charges and the GAFF force field.

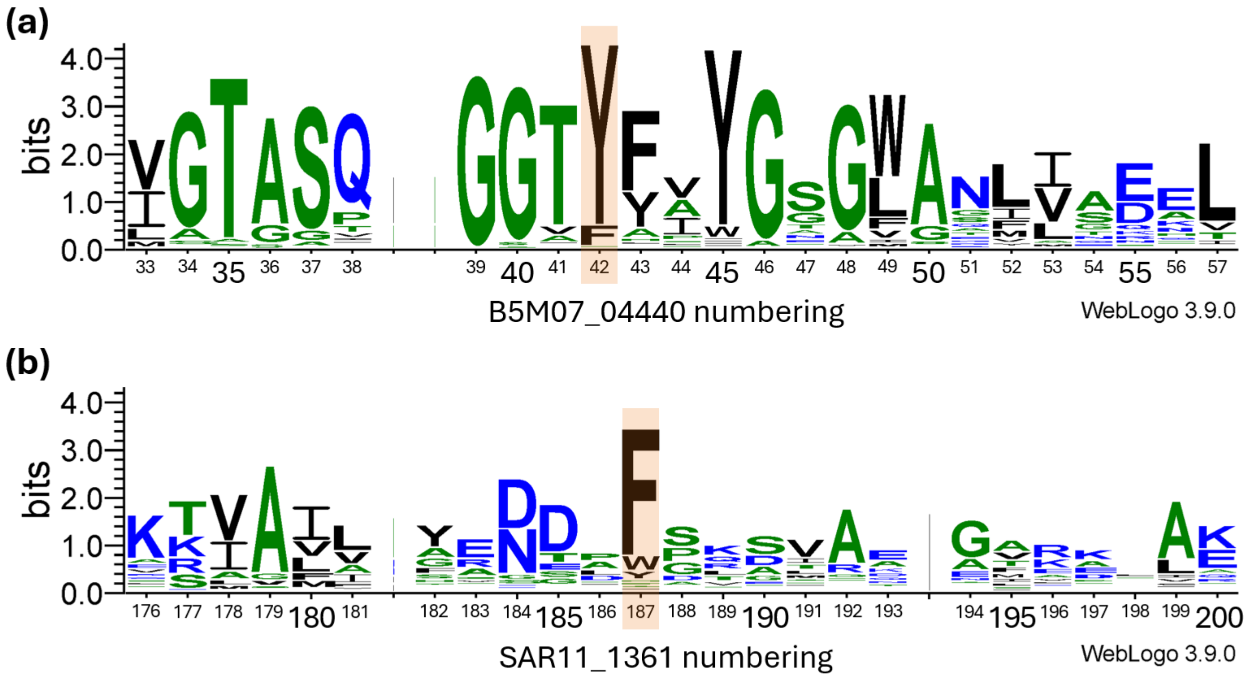


**Figure S14. Aromatic residues in the binding pocket are conserved in SBPs.** Multiple sequence alignments of (a) B5M07_04440 and (b) SAR11_1361 show that the aromatic residues Tyr42 and Phe187 are highly conserved across B5M07_04440 and SAR11_1361 homologs, respectively. BLAST searches for B5M07_04440 and SAR11_1361 homologs were performed against the UniRef90 database (3). 500 sequence hits were returned from each BLAST search and aligned using MAFFT (4). A portion of each alignment was used to create the logos shown here using the WebLogo tool (5), with additional labelling of the x-axis carried out in PowerPoint. Residue numbering for (a) and (b) is based on the sequences of B5M07_04440 and SAR11_1361, respectively. Aromatic residues of interest are highlighted in orange.

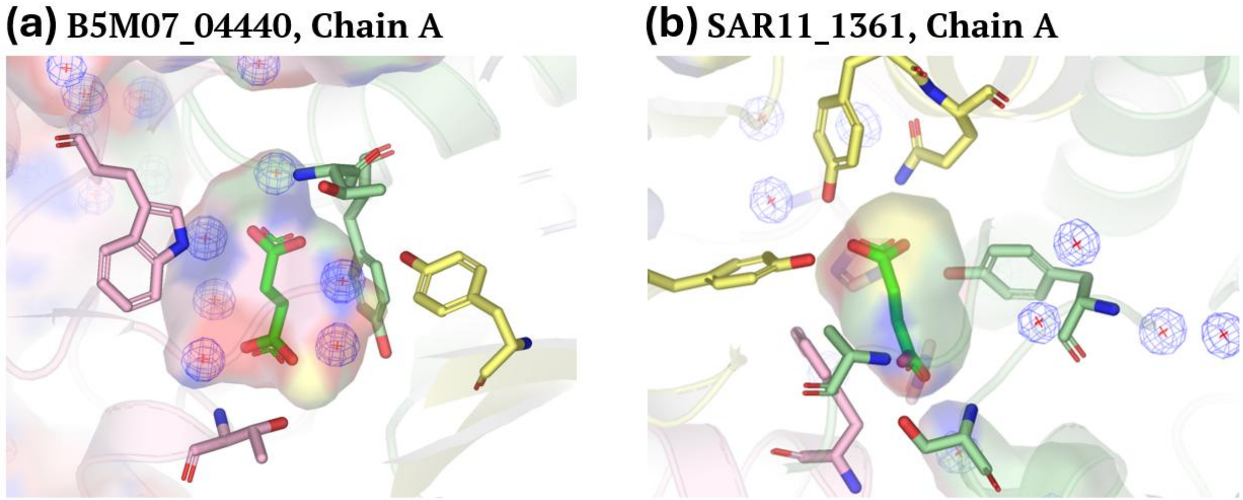


**Figure S15. Water molecules at the ligand binding site of B5M07_04440 and SAR11_1361.** (a) The X-ray crystal structure of B5M07_04440 has electron density corresponding to six water molecules at the ligand binding site when succinate is bound. (b) The X-ray crystal structure of SAR11_1361 does not show electron density corresponding to water molecules at the ligand binding site when succinate is bound. Waters are shown as red crosses and the $2F_o–F_c$ electron density map as blue mesh ($\sigma = 1.5$).

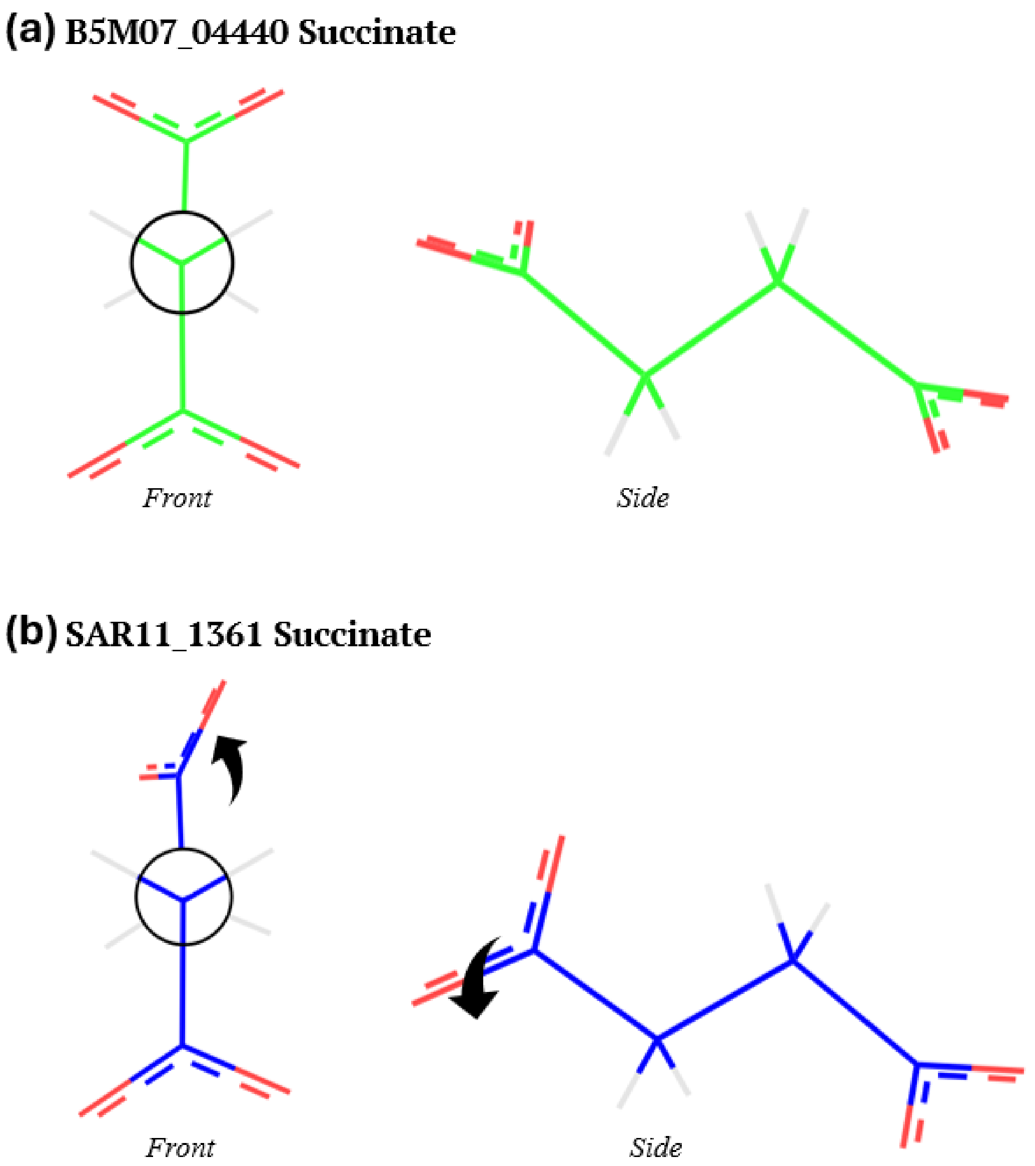


**Figure S16. The conformation of succinate bound in B5M07_04440 vs SAR11_1361.** (a) The front and side view of succinate in the conformation adopted when bound at the binding site of B5M07_04440 according to the X-ray crystal structure. Here succinate adopts the anti-conformation. (b) The front and side view of succinate in the conformation adopted when bound at the binding site of SAR11_1361 according to the X-ray crystal structure. Here succinate adopts the anti-conformation, however one carboxylate group (indicated by black arrows) is twisted relative to the axis of the carbon backbone.

**Method S1. Protein-ligand system preparation for MD simulations of B5M07_04440.**

The coordinates of B5M07_04440 and 2,2-difluorosuccinate were derived from the X-ray crystal structure, chain A. The coordinates of succinate were obtained after aligning chain A of the X-ray crystal structure to that of the B5M07_04440-2,2-difluorosuccinate crystal structure. The tetrafluorosuccinate model was prepared via addition of two fluorine atoms placed using the 2,2-difluorosuccinate pose as a structural reference. The ligand structures used for MD simulations were verified and found to be superimposable, as shown in Figure S12.

**Method S2. Ligand parameterization for MD simulations of B5M07_04440.**

Ligand modelling and parameterization were carried out using AmberTools (6), the CHARMM-GUI webserver (7), and PyMOL (8). Ligand parameters were generated using two charge-assignment strategies: (i) AM1-BCC and (ii) RESP. AM1-BCC (9) charges were assigned, and ligand parameters were generated using the GAFF force field (10). RESP charges were derived using the standard two-stage RESP protocol (11). Geometry optimization and electrostatic potential calculations were performed at the HF/6-31+G(d) level using Gaussian09 (12). The resulting charges were combined with the GAFF2 force field (13). The starting coordinates for succinate and 2,2-difluorosuccinate were derived from the X-ray crystal structures, chain A. For tetrafluorosuccinate, the ligand geometry was generated from the 2,2-difluorosuccinate coordinates by substituting two hydrogen atoms with fluorine atoms. The assigned charges from both the two-stage RESP and AM1-BCC fitting are included in Tables S5 and S6, respectively. The resulting ligand parameters were then used for the MD simulation.

**Method S3. System preparation for MD simulations of B5M07_04440.**

Protonation states of B5M07_04440 from chain A of the X-ray crystal structures, were assigned using the H++ webserver (14) at pH 7.5. Six conserved water molecules present in the ligand-binding site in the X-ray crystal structures (Figure S15) were included before solvation, as the removal of these waters led to the formation of persistent hydrogen bonding between the ligand and Tyr117's side chain in the simulations, which was not observable in the crystal structure. The B5M07_04440–ligand complex was then solvated in a TIP3P (15) water box with dimensions of 12 Å on each side. To match the experimental conditions, $Na^+$ and $Cl^-$ ions were subsequently added to simulate 300 mM NaCl salt concentration, as calculated by the SLTCAP (16) server, after neutralizing the system. The B5M07_04440 protein was parameterized using the AMBER ff14SB force field (17).

**Method S4. Molecular dynamics simulations of B5M07_04440.**

Molecular dynamics simulations were performed using AMBER 24 (6). Periodic boundary conditions were applied throughout the simulations, with a 10 Å nonbonded cutoff. Long-range electrostatic interactions were treated using the Particle Mesh Ewald (PME) method with a real-space cutoff of 10 Å and a B-spline interpolation order of 4. The system was first subjected to a three-stage energy minimization protocol. In the first minimization step, 5000 cycles of minimization were performed, consisting of 3000 cycles of steepest descent followed by 2000 cycles of conjugate-gradient minimization, with harmonic positional restraints of 10 kcal $mol^{-1}$ $Å^{-2}$ applied to the heavy atoms of the ligand and protein residues. A second restrained minimization was then performed using 3000 cycles of steepest descent followed by 2000 cycles of conjugate-gradient minimization, with the restraint force constant reduced to 2 kcal $mol^{-1}$ $Å^{-2}$. Finally, an unrestrained minimization of 2000 cycles was carried out. All minimization steps were performed with periodic boundary conditions at constant volume and a 10 Å nonbonded cutoff.

After minimization, three independent replicas were generated by assigning different initial velocities using independent random seeds. Each simulation system was gradually heated under the NVT ensemble in three consecutive 100 ps stages, using a 2 fs time step. Bonds involving hydrogen atoms were constrained using the SHAKE algorithm (18). The temperature was gradually increased from 10 to 100 K, then from 100 to 200 K, and finally from 200 to 298 K using Langevin dynamics with a collision frequency of 2.0 $ps^{-1}$. During heating, harmonic restraints of 2 kcal $mol^{-1}$ $Å^{-2}$ were applied to the heavy atoms of the protein and ligand.

The heated system was then equilibrated under the NPT ensemble at 298 K and 1 bar. During the first three NPT equilibration stages, restraints were gradually released from 2.0 to 1.0 and then 0.5 kcal $mol^{-1}$ $Å^{-2}$. These restraints were applied to the ligand heavy atoms and the protein backbone atoms. Each restrained NPT equilibration stage was run for 100 ps with a 2 fs timestep. A final unrestrained NPT equilibration was then performed for 500 ps. Langevin dynamics with a collision frequency of 2.0 $ps^{-1}$ was used for temperature control, and pressure was controlled using the

Berendsen weak-coupling barostat with isotropic pressure scaling and a pressure relaxation time of 2.0 ps. Trajectories were written in NetCDF format with coordinate wrapping enabled.

Production simulations were subsequently carried out as consecutive 100 ns segments under the NPT ensemble at 298 K and 1 bar using a 2 fs time step for a total duration of 1 μs for each replicate. Bonds involving hydrogen atoms were constrained using SHAKE. Temperature was maintained using Langevin dynamics with a collision frequency of 0.5 $ps^{-1}$, and pressure was maintained using a Monte Carlo barostat with isotropic pressure scaling. No positional restraints were applied during production. Coordinates, restart files, and energy information were written every 5000 steps, corresponding to every 10 ps of the simulation.
